# Simulating 500 million years of evolution with a language model

**DOI:** 10.1101/2024.07.01.600583

**Authors:** Thomas Hayes, Roshan Rao, Halil Akin, Nicholas J. Sofroniew, Deniz Oktay, Zeming Lin, Robert Verkuil, Vincent Q. Tran, Jonathan Deaton, Marius Wiggert, Rohil Badkundri, Irhum Shafkat, Jun Gong, Alexander Derry, Raul S. Molina, Neil Thomas, Yousuf A. Khan, Chetan Mishra, Carolyn Kim, Liam J. Bartie, Matthew Nemeth, Patrick D. Hsu, Tom Sercu, Salvatore Candido, Alexander Rives

## Abstract

More than three billion years of evolution have produced an image of biology encoded into the space of natural proteins. Here we show that language models trained on tokens generated by evolution can act as evolutionary simulators to generate functional proteins that are far away from known proteins. We present ESM3, a frontier multimodal generative language model that reasons over the sequence, structure, and function of proteins. ESM3 can follow complex prompts combining its modalities and is highly responsive to biological alignment. We have prompted ESM3 to generate fluorescent proteins with a chain of thought. Among the generations that we synthesized, we found a bright fluorescent protein at far distance (58% identity) from known fluorescent proteins. Similarly distant natural fluorescent proteins are separated by over five hundred million years of evolution.

## Introduction

The proteins that exist today have developed into their present forms over the course of billions of years of natural evolution, passing through a vast evolutionary sieve. In parallel experiments conducted over geological time, nature creates random mutations and applies selection, filtering proteins by their myriad sequences, structures, and functions.

As a result, the patterns in the proteins we observe reflect the action of the deep hidden variables of the biology that have shaped their evolution across time. Gene sequencing surveys of Earth’s natural diversity are cataloging the sequences (1–3) and structures (4, 5) of proteins, containing billions of sequences and hundreds of millions of structures that illuminate patterns of variation across life. A consensus is developing that underlying these sequences is a fundamental language of protein biology that can be understood using language models (6–11).

A number of language models of protein sequences have now been developed and evaluated (5–10, 12–17). It has been found that the representations that emerge within language models reflect the biological structure and function of proteins (6–8, 18), and are learned without any supervision on those properties (19, 20), improving with scale (5, 21). In the field of artificial intelligence, scaling laws have been found that predict the growth in capabilities with increasing scale, describing a frontier in compute, parameters, and data (22–24).

We present ESM3, a frontier multimodal generative model that reasons over the sequences, structures, and functions of proteins. ESM3 is trained as a generative masked language model over discrete tokens for each modality. Structural reasoning is achieved by encoding three-dimensional atomic structure as discrete tokens rather than with the complex architecture and diffusion in three-dimensional space employed in recent predictive (25) and generative models (26– 28) of proteins. All-to-all modeling of discrete tokens is scalable, and allows ESM3 to be prompted with any combination of its modalities, enabling controllable generation of new proteins that respect combinations of prompts.

ESM3 at its largest scale was trained with 1.07 × 10^24^ floating point operations (FLOPs) on 2.78 billion proteins and 771 billion unique tokens, and has 98 billion parameters. Scaling ESM3 to this 98 billion parameter size results in improvements in the representation of sequence, structure, and function, as well as on generative evaluations. We observe that ESM3 is highly responsive to prompts, and finds creative solutions to complex combinations of prompts, including solutions for which we can find no matching structure in nature. Models at all scales can be aligned to better follow prompts, and larger models are far more responsive to alignment, showing greater capability to solve the hardest prompts after alignment.

We report the generation of a new green fluorescent protein (GFP) with ESM3. Fluorescent proteins are responsible for the glowing colors of jellyfish and corals (29) and are important tools in modern biotechnology (30). They share an elegant structure: an eleven stranded beta barrel with a helix that threads its center, which scaffolds the formation of a light-emitting chromophore out of the protein’s own atoms. This mechanism is unique in nature—no other protein spontaneously forms a fluorescent chromophore out of its own structure—suggesting that producing fluorescence is hard even for nature.

Our new protein, which we have named esmGFP, has 36% sequence identity to *Aequorea victoria* GFP, and 58% sequence identity to the most similar known fluorescent protein. Despite GFP’s intense focus as a target for protein engineering over several decades, as far as we are aware, new GFPs this distant have only been found through their discovery in nature.

Similar amounts of diversification among natural GFPs have occurred over predictable timescales. Understood in these terms, the generation of a new fluorescent protein at this distance from existing proteins appears to be equivalent to simulating over 500 million years of evolution.

### ESM3

ESM3 achieves a scalable generative model of the three fundamental properties of proteins, sequence, structure, and function, through language modeling. Previous generative modeling efforts for proteins have focused primarily on individual modalities, leveraging complex architectures and training objectives for structure that represent proteins as three-dimensional objects. To date, the only language models that have been scaled are for protein sequences. In ESM3 sequence, structure, and function are represented through alphabets of discrete tokens. The modalities are input and output as separate sequence tracks that are fused into a single latent space within the model. This simplicity enables ESM3 to leverage a scalable transformer architecture to train up to 98 billion parameters and more than one trillion teraflops of compute, demonstrating the emergence of complex reasoning capabilities over sequence, structure, and function.

ESM3 is trained with a generative masked language modeling objective across all its tracks:

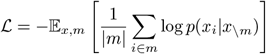

A random mask *m* is applied to the tokens *x* describing the protein, and the model is supervised to predict the identity of the tokens that have been masked. During training, the mask is sampled using a noise schedule that varies the fraction of positions that are masked so that ESM3 sees many different combinations of masked sequence, structure, and function, and predicts completions of any combination of the modalities from any other. This differs from classical masked language modeling (31) in that the supervision is applied across all possible masking rates rather than a single fixed masking rate. This supervision factorizes the probability distribution over all possible predictions of the next token given any combination of previous tokens, ensuring that tokens can be generated in any order from any starting point (32–34).

To generate from ESM3, tokens are iteratively sampled. Starting from a fully or partially masked context, tokens can be sampled one at a time, or in parallel, in any order, until all positions are fully unmasked (Fig. 1A). In addition to enabling generation, ESM3’s training objective is also effective for representation learning. High masking rates improve the generative capability, while lower masking rates improve representation learning. We chose to train ESM3 with a noise schedule that balances generative capabilities with representation learning (Appendix A.2.2).

**Figure 1.**
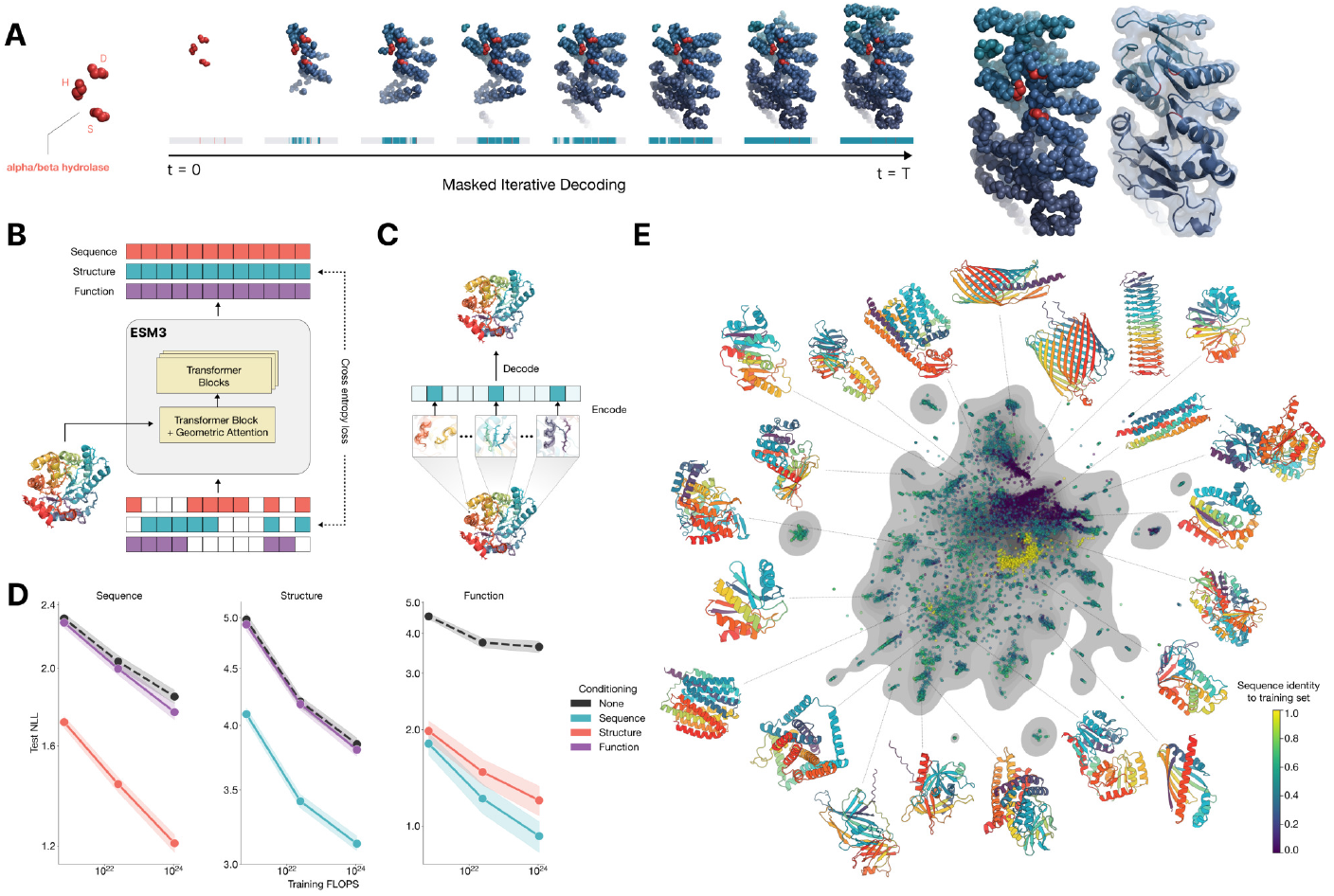
ESM3 is a generative language model that reasons over the sequence, structure, and function of proteins. (A) Iterative sampling with ESM3. Generation of an alpha/beta hydrolase. Sequence, structure, and function can all be used to prompt the model. At each timestep t, a fraction of the masked positions are sampled until all positions are unmasked. (B) ESM3 architecture. Sequence, structure, and function are represented as tracks of discrete tokens at the input and output. The model is a series of transformer blocks, where all tracks are fused within a single latent space; geometric attention in the first block allows conditioning on atomic coordinates. ESM3 is supervised to predict masked tokens. (C) Structure tokenization. Local atomic structure around each amino acid is encoded into tokens. (D) Models are trained at three scales: 1.4B, 7B, and 98B parameters. Negative log-likelihood (averaged across mask rates) on test set as a function of training FLOPs shows response to conditioning on each of the input tracks, improving with increasing FLOPs (95% c.i.). (E) Unconditional generations from ESM3 98B (colored by sequence identity to the nearest sequence in the training set), embedded by ESM3, and projected by UMAP alongside randomly sampled sequences from UniProt (in gray). Generations are diverse, high quality, and cover the distribution of natural sequences.

ESM3 is a bidirectional transformer. Sequence, structure, and function tokens are embedded and fused at the input, then processed through a stack of transformer blocks (Fig. 1B). At the output of the model, shallow multi-layer perceptron (MLP) heads project the final layer representation into token probabilities for each of the tracks. ESM3 uses tokenization, rather than specialized architectural components, to represent the complexity of proteins in a learned multimodal feature space. This enables efficient and highly scalable training.

Protein structures are tokenized by a discrete autoencoder (35), which is trained to compress three-dimensional structure into discrete tokens (Fig. 1C). We propose an invariant geometric attention mechanism to efficiently process three-dimensional structure. The mechanism operates in local reference frames defined by the bond geometry at each amino acid, and allows local frames to interact globally through a transformation into the global frame (Appendix A.1.6). The local structural neighborhoods around each amino acid are encoded into a sequence of discrete tokens, one for each amino acid.

When predicting or generating protein structure, the structure tokens output by ESM3 are passed through the decoder, which reconstructs the full atomic structure. The autoen-coder is trained to encode and reconstruct atomic coordinates with a geometric loss that supervises the pairwise distances and relative orientations of bond vectors and normals (Appendix A.1.7.3.1). This tokenization delivers near-perfect reconstruction of protein structure (<0.5Å RMSD on CAMEO, Fig. S3).

Since the local neighborhoods of each structure token contain information about neighboring parts of the structure, we also provide the model with a mechanism to condition on backbone atomic coordinates directly via geometric attention in the first transformer block. To support higher level abstractions of structure, we include tracks for secondary structure (SS8) tokens and solvent accessible surface area (SASA) tokens. Keywords describing biological activity, such as binding, enzymatic function, and domain or fold classifications, allow an even higher level semantic description of protein architecture and function. Derived from free-text descriptions in InterPro (36) and Gene Ontology (GO) terms for each residue, these keywords are tokenized (Appendix A.1.8), embedded, and summed at the network input. Residue level annotations provide multi-hot labeling of the functions of individual residues, such as catalytic sites and post-translational modifications (Appendix A.1.8.3).

The largest ESM3 model is trained on 2.78 billion natural proteins collected from sequence and structure databases (2, 37–40). Since a small fraction of structures have been experimentally determined relative to sequences, we leverage predicted structures (4, 5). Sequences are annotated with function keywords using a library of hidden markov models (36). We also generate synthetic sequences with an inverse folding model (described in Appendix A.2.1.3) for all structures, including predicted ones. Overall this increases training data to 3.15 billion protein sequences, 236 million protein structures, and 539 million proteins with function annotations, totaling 771 billion unique tokens. Full details of the training dataset are described in Appendix A.2.1.

We train ESM3 models at three scales: 1.4 billion, 7 billion, and 98 billion parameters. In an initial series of experiments to evaluate representation learning performance in response to architecture hyperparameters, we found a greater response to increasing depth than to width. This informed the choice of relatively deep networks for the final architectures, with the 98 billion parameter model incorporating 216 transformer blocks (Appendix A.1.5).

Scaling ESM3 from 1.4 billion to 98 billion parameters results in substantial improvements in the loss for all tracks on the test set, with the greatest improvements observed in sequence loss (Fig. 1D, Fig. S11). The gap between unconditional and conditional negative log-likelihoods increases with scale. Conditioning on function keywords primarily constrains sequence at high masking rates, so while responsiveness to keyword conditioning is observed at high mask rates, it is less apparent in the averaged negative log-likelihood (Fig. S12). These gains in test loss lead to better representation learning (Table S8 and Fig. S8). In single sequence structure prediction ESM3 98B surpasses ESMFold (0.880 vs 0.861 mean local distance difference test, LDDT, CAMEO test set; Table S9). Generating sequences from the model without prompting (unconditional generation) produces high quality proteins—with a mean predicted LDDT (pLDDT) 0.84 and predicted template modeling score (pTM) 0.52—that are diverse in both sequence (mean pairwise sequence identity 0.155) and structure (mean pairwise TM score 0.48), spanning the distribution of known proteins (Fig. 1E, Fig. S14).

Our results show that scaling with language modeling, enabled by tokenization, efficient architectures, and masked token prediction, yields continued improvements in both representational and generative applications. This approach allows the model to build a shared multimodal representation space that is learned from the data, rather than explicitly hardcoded into its architecture, which given increasing compute and data could learn an increasingly richer and more general feature space. In the following sections we show that this approach achieves high fidelity for the controllable generation of proteins.

### Programmable design with ESM3

We explore the ability of ESM3 to follow complex prompts with different compositions. ESM3 can be prompted with instructions from each of its input tracks: sequence, structure coordinates, secondary structure (SS8), solvent-accessible surface area (SASA), and function keywords. This allows prompts to be specified at multiple levels of abstraction, from atomic level structure to high level keywords describing the function and fold topology.

We evaluate ESM3’s ability to follow prompts in each of the tracks independently (Fig. 2A). A set of prompts are constructed for each of the tracks using a temporally held out test set of natural proteins (Appendix A.3.8). The resulting generations are evaluated using ESMFold for consistency with the prompt and confidence of structure prediction (pTM). We define consistency metrics for each track: constrained site RMSD (cRMSD), the RMSD between the coordinates of the prompt, i.e. the positions of the backbone atoms, and the corresponding coordinates in the generation; SS3 accuracy, the fraction of residues where three-class secondary structure between the prompt and generations match; SASA Spearman *ρ*, the correlation between the SASA prompt and the corresponding region of the generation; and keyword recovery, the fraction of prompt keywords recovered by InterProScan (36). Across all tracks, the 7B parameter ESM3 finds solutions that follow the prompt and have structures which are confidently predicted by ESM-Fold (pTM > 0.8).Some mode switching is observed, including under keyword prompting, where a fraction of the generations have confidently predicted structures that do not recover the keywords.

**Figure 2.**
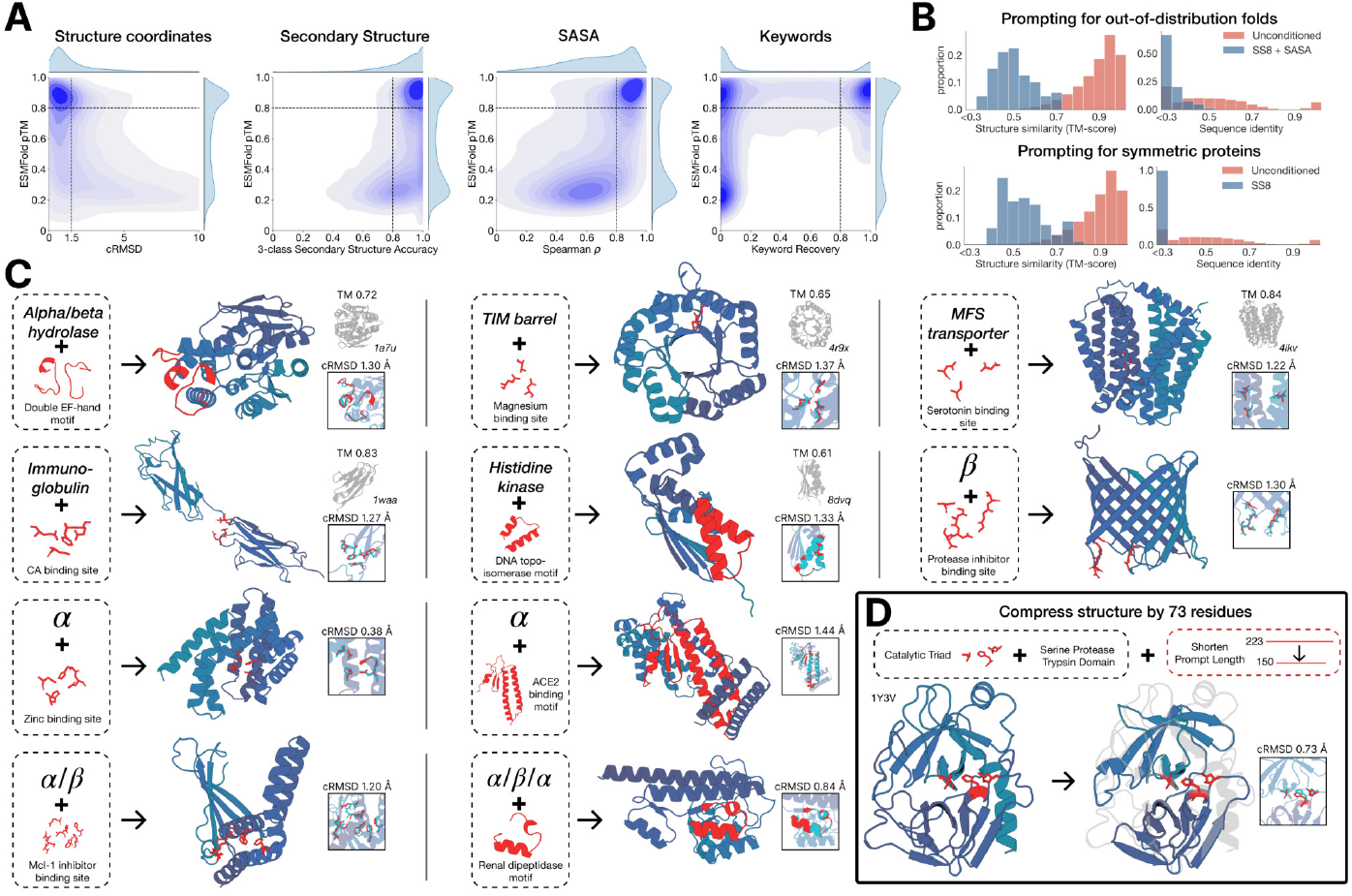
Generative programming with ESM3. (A) ESM3 can follow prompts from each of its input tracks. Density of faithfulness to prompting for each of the tracks is shown. Generations achieve consistency with the prompt (backbone cRMSD, SS3 accuracy, SASA Spearman *ρ*, keyword recovery) and high structure prediction confidence (pTM). (B) ESM3 can be prompted to generate proteins that differ in structure (left) and sequence (right) from the training set and natural proteins. Prompted generations (blue) shift toward a more novel space vs. unconditional generations (red), in response to prompts derived from out-of-distribution natural structures (upper panel) and computationally designed symmetric proteins (lower panel). (C) ESM3 generates creative solutions to a variety of combinations of complex prompts. We show compositions of atomic level motifs with high level instructions specified through keywords or secondary structure prompts. Fidelity to the prompt is shown via similarity to a reference structure (for keyword prompts) and all-atom RMSD (for motif prompts). Solutions differ from the scaffolds where the motif prompt was derived (median TM-score 0.36 ± 0.14), and for many motifs (e.g. serotonin, calcium, protease inhibitor, and Mcl-1 inhibitor binding sites), we could find no significant similarity to other proteins that contain the same motif. (D) An example of especially creative behavior. ESM3 compresses a serine protease by 33% while maintaining the active site structure.

Unconditional generations reflect the distribution of natural proteins. Since we observed ESM3 can faithfully follow prompts, we reasoned that prompting could steer the model to generate proteins that differ from the training set and natural proteins. First we test the ability of the model to follow out-of-distribution prompts. We construct a set of prompts combining SS8 and SASA from held out structures (TM < 0.7 to training set). Under these prompts, while the model continues to generate coherent globular structures (mean pTM 0.85 ± 0.03), the distribution of similarities to the training set (as measured by TM-score and sequence identity) shifts to be more novel (average sequence identity to nearest training set protein < 20% and mean TM-score 0.48 ± 0.09; Fig. 2B top). To test the ability to generalize to structures beyond the distribution of natural proteins, we use secondary structure prompts derived from a dataset of artificial symmetric protein designs distinct from the natural proteins found in the training dataset (Appendix A.3.9). Similarly, ESM3 produces high confidence generations (pTM > 0.8, pLDDT > 0.8) with low sequence and structure similarity to proteins in the training set (sequence identity < 20% and TM-score 0.52 ± 0.10; Fig. 2B bottom), indicating that the model can be used to generate protein sequences and structures highly distinct from those that exist in nature.

ESM3 is able to follow complex prompts, and has the ability to compose prompts from different tracks, and at different levels of abstraction. To evaluate this ability, we prompt ESM3 with motifs that require solving for spatial coordination of individual atoms, including atoms participating in tertiary contacts between residues far apart in the sequence, such as catalytic centers and ligand binding sites. We combine the atomic level motif prompts with high level prompts, either secondary structure prompts or keyword prompts that specify the fold architecture. For each unique combination of atomic level motif and high level prompt, we generate sequences until there is a success (for atomic level prompts, when all-atom RMSD < 1.5Å ; for fold architecture keyword prompts, when TM > 0.6 to a representative structure; for secondary structure prompts, when SS3 accuracy > 80%; and pTM > 0.8, pLDDT > 0.8 for the entire generated protein).

We find that ESM3 is able to solve a wide variety of such tasks (Fig. 2C). It does so without retrieving the motif’s original scaffold (median TM-score of 0.40 ± 0.10 ; Appendix A.3.10). In some cases, the scaffolds are transferred from existing proteins which have similar motifs (for example, the ESM3-designed alpha-helical scaffold for the zinc-binding motif has high similarity to Ni_2+_-binding proteins, PDB: 5DQW, 5DQY; Fig. 2C, row 3 column 1). For many motifs (e.g., binding sites for serotonin, calcium, protease inhibitor, and Mcl-1 inhibitor) Foldseek (41) finds no significant similarity to other proteins that contain the same motif. In these cases we observe that sometimes the motif has been grafted into entirely different folds (e.g. a protease inhibitor binding site motif in a beta-barrel which is most similar to a membrane-bound copper transporter, PDB: 7PGE; Fig. 2C, row 3 column 3). At other times, the scaffold appears to be entirely novel, such as an alpha/beta protein designed to scaffold the Mcl-1 inhibitor binding motif, which has low structural similarity to all known proteins in the PDB, ESMAtlas, and the AlphaFold databases (max. TM-score < 0.5; Fig. 2C, row 4 column 1). Overall, the generated solutions have high designability, i.e. confident recovery of the original structure after inverse folding with ESM-IF1 (42) and refolding with ESMFold (median pTM 0.80 ± 0.08; scTM 0.96 ± 0.04; Appendix A.3.10).

Through experiments with prompt engineering, we have observed especially creative responses to prompts. Here, we highlight an example of protein compression (Fig. 2D). Starting from a natural trypsin (PDB 1Y3V), we prompt with the sequence and coordinates of the catalytic triad as well as functional keywords describing trypsin, but reduce the overall generation length by a third (from 223 to 150 residues). ESM3 maintains the coordination of the active site (all-atom RMSD 0.73Å) and the overall fold with high designability (pTM 0.84, scTM mean 0.97, std 0.006), despite the significant reduction in sequence length and the fold only being specified by the function keyword prompt (Appendix A.3.11).

These examples illustrate ESM3’s ability to find creative solutions to prompts specified in any of its input tracks, individually or in combination. This capability enables a rational approach to protein design, providing control at various levels of abstraction, from high level topology to atomic coordinates, using a generative model to bridge the gap between the prompt and biological complexity.

### Biological alignment

While we have observed meaningful increases in the performance of the base models with scale, larger models could have even greater latent capabilities that we do not observe. The base ESM3 models can be prompted to perform difficult tasks such as tertiary motif scaffolding and composition of prompts, despite the fact that the models have not been explicitly optimized for these objectives. Since the properties we evaluate generative outputs on—such as adherence to the prompt, or the confidence of the scaffold—are only seen by the model indirectly during pretraining, aligning the model directly to the generative task with finetuning could elicit even greater capability differences with larger models.

We study how the base models can be aligned (43, 44) to generate proteins that satisfy challenging prompts. For each model, we construct a dataset of backbone atomic coordinate prompts, consisting of contiguous spans of residues, and tertiary motifs (which also specify the identities of the contacting amino acids). We generate multiple protein sequences for each prompt and fold each of the sequences using ESM3, scoring for consistency with the prompt (backbone cRMSD) and structure prediction confidence (pTM). High quality samples are paired with low quality samples for the same prompt to construct a preference dataset (Appendix A.4). ESM3 is then finetuned with a preference optimization loss (45, 46), which causes the model to put higher likelihood on the high quality samples relative to the low quality samples.

After aligning each of the base models, we evaluate their absolute performance, and the shift in the distribution of generations. We focus on a series of challenging prompts that require coordination of the backbone atoms of residues in tertiary contact. We evaluate the ability to generate high quality scaffolds (pTM > 0.8) that follow the prompt with high resolution (backbone cRMSD < 1.5Å), using ESM-Fold for the evaluation. We prompt each model with amino acid identities and backbone atomic coordinates from a held-out dataset of 46 ligand binding motifs (Appendix A.4.5). For each motif, we create 1024 prompts by permuting the order of the residues, varying their position in the sequence, and varying the length of the sequence. A single protein is generated per prompt. The 1024 generations for each motif are used to construct an unbiased estimator of the fraction of tertiary coordination tasks solved after 128 generations (Pass@128; Appendix A.4.5).

Aligned models solve double the tertiary coordination tasks compared to base models (Fig. 3A). While the base models show differences in the percentage of tasks solved (9.5% for 1.4B, 19.0% for 7B, 26.8% for 98B; Fig. 3A), a much larger capability difference is revealed through alignment (increasing from 9.5% to 18.8%, 19.0% to 37.4%, and 26.8% to 65.5% for the 1.4B, 7B and 98B models, respectively). Preference-tuned models not only solve a greater proportion of tasks, but also find a greater number of solutions per task, as evaluated by the number of distinct structural clusters (TM > 0.8) with backbone cRMSD < 1.5Å and pTM > 0.8 (Fig. 3B). A shift in the distribution of ESMFold pTM and backbone cRMSD for each ligand binding motif is observed (Fig. 3C; Fig. S18). At the 98B scale, the finetuned model produces more distinct successful clusters than the base model on 37 of the 46 tested ligands, while the remaining 9 ligands were not solved by either the base or aligned model, indicating that alignment almost universally improves the faithfulness to the prompt and confidence of the structure prediction for the generated proteins. These results represent state-of-the-art motif scaffolding performance (Table S16). Compared to a supervised finetuning baseline, which only maximizes the likelihood of the positive examples, preference tuning leads to larger improvements at all scales (Appendix A.4.6).

**Figure 3.**
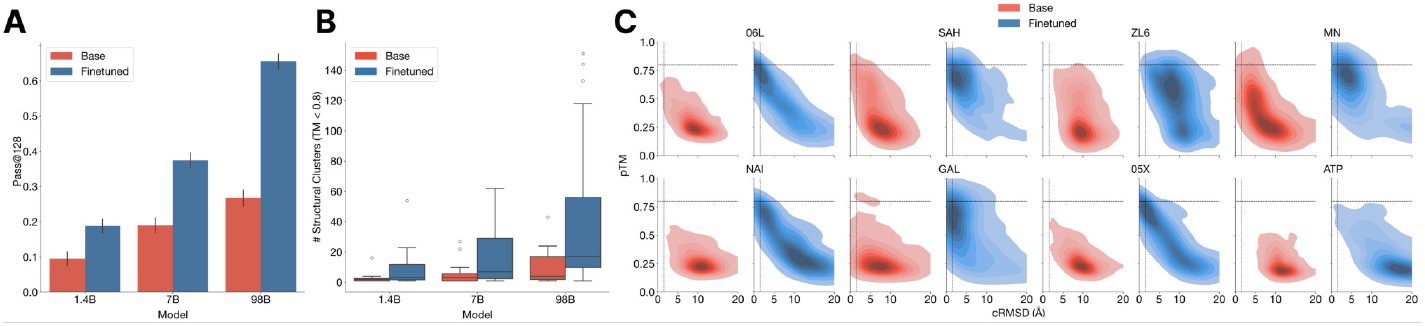
The ability to solve complex tasks increases with scale through alignment. ESM3 is aligned to follow tertiary coordination prompts with a dataset of preference pairs constructed from prompted generations, where positive samples with good scores for desired properties (high pTM, low cRMSD) are paired with negative samples with worse scores. The preference tuning loss encourages the model to put higher likelihood on the positive samples. After training, models are evaluated by prompting with the backbone atomic coordinates of residues in tertiary contact. (A) We show the effect of finetuning on the fraction of tasks solved with 128 generations (Pass@128; 2 s.d. error bars). A large gap opens between the models with scale. The response to alignment shows a latent capability to solve complex tasks in the largest model. (B) Number of distinct solutions (clustered at TM *>* 0.8) generated for each tertiary motif. After finetuning there are often many unique solutions for ligands where there are successes. (C) Densities of prompted generations are shown for the base model (left) and aligned model (right) at the 98B scale for a number of randomly selected ligands. After alignment, the fidelity to the prompt (backbone cRMSD) and quality of generations (pTM) tends to improve substantially.

Our experiments with alignment reveal a considerable difference in capabilities between model scales. The largest aligned model improves substantially relative to the base model before alignment, as well as in comparison to the smaller models after alignment. Through alignment the models learn to generalize from a small number of examples: the distribution of generations shifts to improve the quality of scaffolds, and consistency with prompts, increasing the fraction of tasks solved and the number of distinct solutions.

Alignment requires the models to learn by example. The ability to identify the underlying properties that are illustrated by the finetuning examples, and to generalize those demonstrations to new tasks, implies that there is an internal representation of the properties that the finetuning accesses. This representation space is learned through the process of pretraining, where the model is trained on proteins across evolution, which suggests it reflects and contains the immense variety and complexity of protein biology. Such a representation space is likely to contain features that support generalization of many biological properties. The greater responsiveness of larger models to alignment suggests that their internal representation space better approximates those underlying properties, which is evidence of a deep capability for transfer, via the features learned in pretraining, that improves with scale.

### Generating a new fluorescent protein

We sought to understand if the base pretrained ESM3 model has sufficient biological fidelity to generate functional proteins. We set out to create a functional green fluorescent protein (GFP) with low sequence similarity to existing ones. We chose the functionality of fluorescence because it is difficult to achieve, easy to measure, and one of the most beautiful mechanisms in nature.

Responsible for the fluorescence of jellyfish and the vivid colors of coral (47), proteins in the GFP family are unique in their ability to form a fluorescent chromophore without cofactors or substrates (30). This property allows the GFP sequence to be inserted into the genomes of other organisms to visibly label molecules, cellular structures, or processes, providing a foundational toolkit that has been broadly applied across the biosciences.

The GFP family has been the subject of decades of protein engineering efforts, but still the vast majority of the diversity of functional variants has come from prospecting the natural world. Rational design and mutagenesis have yielded GFP sequences with improved properties—such as higher brightness or stability, or differently colored variants—that incorporated small numbers of mutations (typically 5 to 15, out of the total 238 amino acid coding sequence). In a few cases, leveraging high-throughput experimentation and machine learning, scientists have been able to introduce up to 40-50 mutations (i.e. 80% sequence identity) while retaining fluorescence (48–50).

Generating a new GFP would require materialization of the complex biochemistry and physics that underlie its fluorescence. In all GFPs, an autocatalytic process forms the chromophore from three key amino acids in the core of the protein. The unique structure of GFP, a kinked central alpha helix surrounded by an eleven stranded beta barrel with inward facing coordinating residues, enables this reaction (51). Once formed, the chromophore must not just absorb light but also emit it in order to be fluorescent. Light emission is highly sensitive to the local electronic environment of the chromophore. The fitness landscape of GFP reflects the precise configuration of both the active site and the surrounding tertiary interactions required to achieve its function, since a few random mutations are sufficient to reduce fluorescence to zero (48, 52).

In an effort to generate new GFP sequences, we directly prompt the base pretrained 7B parameter ESM3 to generate a 229 residue protein conditioned on the positions Thr62, Thr65, Tyr66, Gly67, Arg96, Glu222, which are critical residues for forming and catalyzing the chromophore reaction (Fig. 4A). We additionally condition on the structure of residues 58 through 71 from the experimental structure in 1QY3, which are known to be structurally important for the energetic favorability of chromophore formation (53). Specifically, sequence tokens, structure tokens, and atomic coordinates of the backbone are provided at the input, and generation begins from a nearly completely masked array of tokens corresponding to 229 residues, except for the token positions used for conditioning.

**Figure 4.**
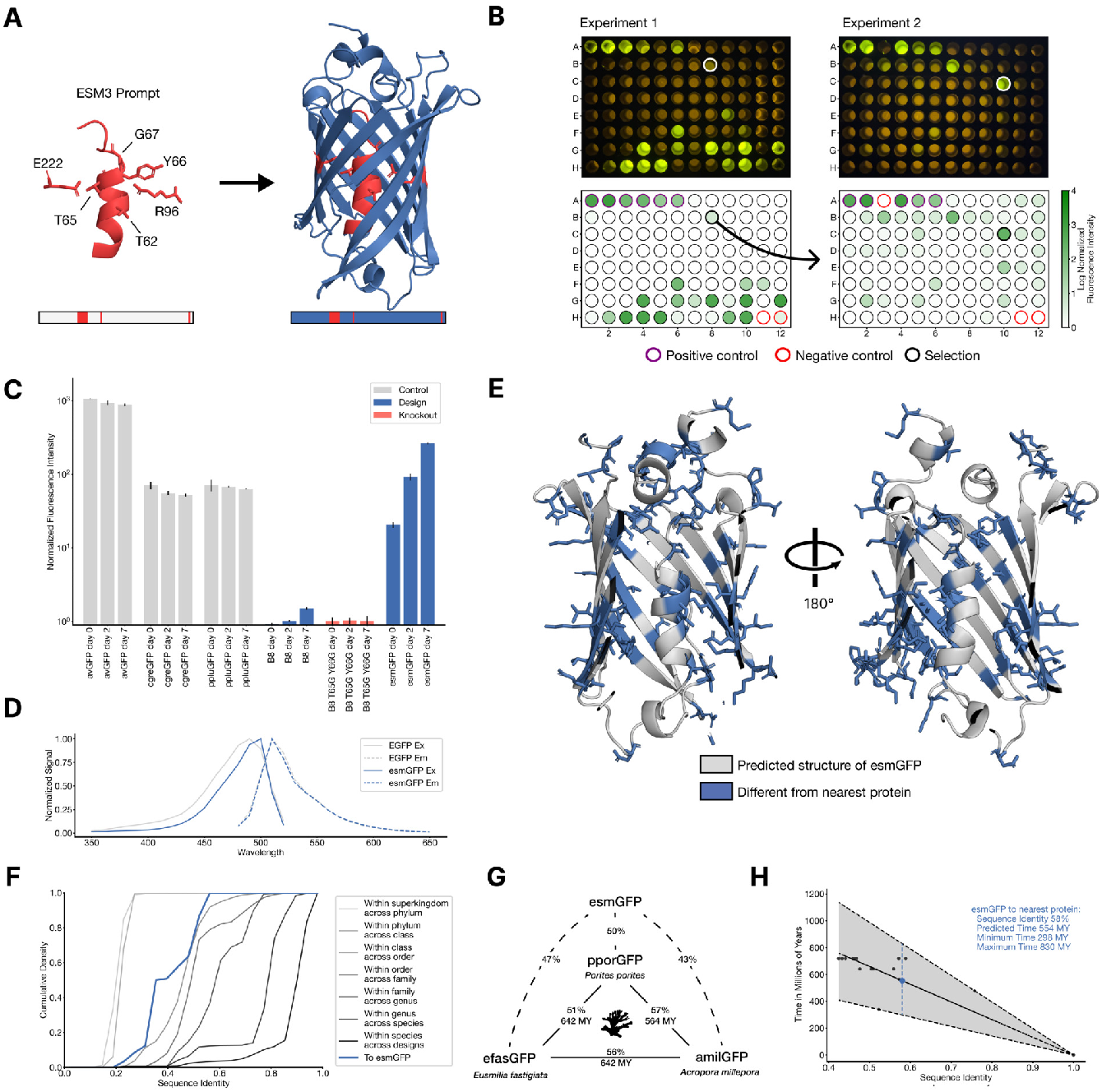
Generating a new fluorescent protein with a chain of thought. (A) We prompt ESM3 with the sequence and structure of residues required for forming and catalyzing the chromophore reaction, as well as the structure of part of the central alpha helix from a natural GFP (left). Through a chain of thought, ESM3 generates design candidates (right). (B) ESM3 found a bright GFP distant from other known GFPs in two experiments. We measured fluorescence in *E. coli* lysate. Top row, photograph of plates. Bottom row, plate reader fluorescence quantification. Positive controls of known GFPs are marked with purple circles, negative controls with no GFP sequence or no *E. coli* are marked with red circles. In the first experiment (left) we expressed designs with a range of sequence identities. A notable design with low sequence identity (57%) to known fluorescent proteins appears in the well labeled B8 (highlighted in a black circle bottom, white circle top). We continue the chain of thought from the protein in B8 for the second experiment (right). A bright design appears in the well labeled C10 (58% sequence identity to known fluorescent proteins, black circle bottom, white circle top) which we designate esmGFP. (C) esmGFP exhibits fluorescence intensity similar to common GFPs. Normalized fluorescence is shown for a subset of proteins in experiment 2. (D) Excitation and emission spectra for esmGFP overlaid on the spectra of EGFP. (E) Two cutout views of the central alpha helix and the inside of the beta barrel of a predicted structure of esmGFP. The 96 mutations esmGFP has relative to its nearest neighbor, tagRFP, are shown in blue. (F) Cumulative density of sequence identity between fluorescent proteins across taxa. esmGFP has the level of similarity to all other FPs that is typically found when comparing sequences across orders, but within the same class. (G) Evolutionary distance by time in millions of years (MY) and sequence identities for three example anthozoan GFPs and esmGFP. (H) Estimator of evolutionary distance by time (MY) from GFP sequence identity. We estimate esmGFP is over 500 million years of natural evolution removed from the closest known protein.

We generate designs using a chain-of-thought procedure as follows. The model first generates structure tokens, effectively creating a protein backbone. Backbones that have sufficiently good atomic coordination of the active site but differentiated overall structure from the 1QY3 backbone pass through a filter to the next step of the chain. We add the generated structure to the original prompt to generate a sequence conditioned on the new prompt. We then perform an iterative joint optimization, alternating between optimizing the sequence and the structure. We reject chains-of-thought that lose atomic coordination of the active site (Appendix A.5.1). We draw a computational pool of 10s of thousands of candidate GFP designs from the intermediate and final points in the iterative joint optimization stage of the generation protocol. We bucket the designs by sequence similarity to known fluorescent proteins and filter and rank designs using a variety of metrics (details in Appendix A.5.1.5).

We performed a first experiment with 88 designs on a 96 well plate, evaluating the top generations in each sequence similarity bucket. Each generated protein was synthesized, expressed in *E. coli*, and measured for fluorescence activity at an excitation wavelength of 485 nm (Fig. 4B left). We measured brightness similar to positive controls from a number of designs that have higher sequence identity with naturally occurring GFPs. We also identify a design in well B8 (highlighted in a black circle) with only 36% sequence identity to the 1QY3 sequence and 57% sequence identity to the nearest existing fluorescent protein, tagRFP. This design was 50x less bright than natural GFPs and its chromophore matured over the course of a week, instead of in under a day, but it presents a signal of function in a new portion of sequence space that to our knowledge has not been found in nature or through protein engineering.

We continue the chain of thought starting from the sequence of the design in well B8 to generate a protein with improved brightness, using the same iterative joint optimization and ranking procedure as above. We create a second 96 well plate of designs, and using the same plate reader assay we find that a few designs in this cohort have a brightness in the range of GFPs found in nature. The best design, located in well C10 of the second plate (Fig. 4B right), we designate esmGFP.

We find esmGFP exhibits brightness in the distribution of natural GFPs. We evaluated the fluorescence intensity at 0, 2, and 7 days of chromophore maturation, and plot these measurements for esmGFP, a replicate of B8, a chromophore knockout of B8, along with three natural GFPs avGFP, cgreGFP, ppluGFP (Fig. 4C). esmGFP takes longer to mature than the known GFPs that we measured, but achieves a comparable brightness after two days. To validate that fluorescence was mediated by the intended Thr65 and Tyr66, we show that B8 and esmGFP variants where these residues were mutated to glycine lost fluorescence activity (Fig. S22).

Analysis of the excitation and emission spectra of esmGFP reveals that its peak excitation occurs at 496 nm, which is shifted 7 nm relative to the 489 nm peak for EGFP, while both proteins emit at a peak of 512nm (Fig. 4D). The shapes of the spectra indicated a narrower full-width-half-maximum (FWHM) for the excitation spectrum of es-mGFP (39mm for esmGFP vs 56 nm for EGFP), whereas the FWHM of their emission spectra were highly comparable (35nm and 39 nm, respectively). Overall esmGFP exhibits spectral properties consistent with known GFPs.

We next sought to understand how esmGFP compares to known proteins. A BLAST (54) search against the non-redundant protein sequences database and an MMseqs (55) search of ESM3’s training set report the same top hit— tagRFP, which was also the nearest neighbor to B8—with 58% sequence identity, representing 96 mutations throughout the sequence. tagRFP is a designed variant, and the closest wildtype sequence to esmGFP from the natural world is eqFP578, a red fluorescent protein, which differs from esmGFP by 107 sequence positions (53% identity). Sequence differences between esmGFP and tagRFP occur throughout the structure (Fig. 4E) with 22 mutations occurring in the protein’s interior, which is known to be highly sensitive to mutations due to chromophore proximity and a high density of interactions (56).

Examination of a sequence alignment of 648 natural and designed GFP-like fluorescent proteins revealed that esmGFP has the level of similarity to all other FPs that is typically found when comparing sequences across taxonomic orders, but within the same taxonomic class (Fig. 4F). For example, the difference of esmGFP to other FPs is similar to the level of difference between FPs belonging to the orders of Scleractinia (stony corals) and Actiniaria (sea anemones) both of which belong to the larger class Anthozoa of marine invertebrates (Fig. 4G). The closest FPs to esmGFP come from the Anthozoa class (corals and anemones; average sequence identity 51.4%), but esmGFP also shares some sequence identity with FPs from the Hydrozoa (jellyfish) where avGFP was discovered (average sequence identity 33.4%; Fig. S23).

We can draw insight from evolutionary biology on the amount of time it would take for a protein with similar sequence identity to arise through natural evolution. In Fig. 4G we show esmGFP alongside three anthozoan GFPs. We use a time-calibrated phylogenetic analysis of the anthozoans (57) that estimated the millions of years ago (MYA) to last common ancestors to estimate evolutionary time between each pair of these species. Using a larger dataset of six anthozoan GFPs and species for which we have accurate MYA to last common ancestors and GFP sequence identities, we construct a simple estimator that correlates sequence identity between FPs to MY of evolutionary time between the species (Fig. 4H) to calibrate against natural evolution. Based on this analysis we estimate esmGFP represents an equivalent of over 500 million years of evolution from the closest protein that has been found in nature.

## Discussion

We have found that language models can reach a design space of proteins that is distant from the space explored by natural evolution, and generate functional proteins that would take evolution hundreds of millions of years to discover. Protein language models do not explicitly work within the physical constraints of evolution, but instead can implicitly construct a model of the multitude of potential paths evolution could have followed.

Proteins can be seen as existing within an organized space where each protein is neighbored by every other that is one mutational event away (58). The structure of evolution appears as a network within this space, connecting all proteins by the paths that evolution can take between them. The paths that evolution can follow are the ones by which each protein transforms into the next without the collective loss of function of the system it is a part of.

It is in this space that a language model sees proteins. It sees the data of proteins as filling this space, densely in some regions, and sparsely in others, revealing the parts that are accessible to evolution. Since the next token is generated by evolution, it follows that to solve the training task of predicting the next token, a language model must predict how evolution can move through the space of possible proteins.

Simulations are computational representations of reality. In that sense, a language model which can predict possible outcomes of evolution can be said to be a simulator of it. ESM3 is an emergent simulator that has been learned from solving a token prediction task on data generated by evolution. It has been theorized that neural networks discover the underlying structure of the data they are trained to predict (59, 60). In this way, solving the token prediction task would require the model to learn the deep structure that determines which steps evolution can take, i.e. the fundamental biology of proteins.

In ESM3’s generation of a new fluorescent protein, it is the first chain of thought to B8 that is the most intri(guin g. At 96 mutations to B8’s closest neighbor there are 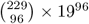 possible proteins, out of which only a vanishingly small fraction can have function, since fluorescence falls off sharply even after just a few random mutations. The existence of C10 and other bright designs in the neighborhood of B8 confirms that in the first chain of thought to B8, ESM3 found a new part of the space of proteins that, although unexplored by nature, is dense with fluorescent proteins.

## ACKNOWLEDGEMENTS

We thank Eric Schreiter, Karel Svoboda, and Srinivas Turaga for feedback on the properties of esmGFP. We thank Ian Holmes for feedback on the evolutionary analysis of esmGFP. We thank Marko Iskander, Vishvajit Kher, and the Andromeda cluster team for support on compute infrastructure. We thank April Pawluk for assistance with manuscript preparation. We also thank the experts who provided feed-back on our approach to responsible development, and the experts who participated in the review of the risks and benefits of releasing ESM3-open.

## CONTRIBUTIONS

**Data:** H.A., Z.L., R.R., A.R., T.S., N.T., R.V.

**Pre-training:** H.A., S.C., J.D., T.H., Z.L., D.O., R.R., A.R., T.S., I.S., R.V., M.W.

**Post-training:** H.A., S.C., A.D., J.G., T.H., D.O., R.R., A.R., M.W.

**Evaluation and Analysis:** R.B., J.D., A.D., T.H., Y.A.K., C.K., Z.L., R.S.M., A.R., N.J.S.

**Open Model & Responsible Development:** J.G., I.S., N.J.S., T.S., R.S.M., Z.L., R.R., A.R., N.T.

**API & Deployment:** J.G., C.M., R.S.M., Z.L., T.S.

**GFP Computational:** S.C., T.H., N.J.S., A.R., R.V.

**GFP Experimental Validation:** L.J.B., M.N., P.D.H., Y.A.K., N.J.S., N.T., V.Q.T.

**Manuscript:** S.C., T.H., R.R., A.R., N.J.S. overall manuscript. All authors contributed to the sections for which they are credited.

**Supplement:** H.A., R.B., L.J.B., S.C., J.D., A.D., T.H., C.K., Z.L., R.S.M., D.O., R.R., A.R., N.J.S., T.S., I.S., N.T., V.Q.T., R.V., M.W.

**Overall Scientific Direction:** A.R.

## MODEL AND DATA AVAILABILITY

Weights and code for ESM3-open are provided for academic research use at https://github.com/evolutionaryscale/esm. They have also been permanently archived (61). The ESM3-open model was reviewed by a committee of technical experts who found that the benefits of releasing the model greatly outweighed any potential risks. ESM3 models will be available via API with a free access tier for academic research. The sequence of esmGFP (along with the other GFPs generated for this work) is committed to the public domain. Plasmids for esmGFP-C10 and esmGFP-B8 will be made available.

## Appendices

### A. Materials and Methods

#### A.1 ARCHITECTURE

##### A.1.1 Notation

In the following, we use *L* to denote the sequence length, *d* for the embedding dimension, {*a*..*b*} to denote the inclusive set of integers from *a* to *b*, and [*a, b*] an interval of real numbers. *SE*(3) is the special Euclidean group, which we use to denote frames (Appendix A.1.6.1).

##### A.1.2 Overview

ESM3 is all-to-all generative model that both conditions on and generates a variety of different tracks. As input, ESM3 is conditioned on various tracks as described in Appendix A.1.5.1, and as output, ESM3 generates predictions detailed in Appendix A.1.5.2.

The generative pipeline is as follows.

###### Tokenization

First, raw inputs are tokenized as described in Appendix A.1.3. Structural inputs are tokenized via a VQ-VAE (Appendix A.1.7). Function keywords are tokenized by quantizing the TF-IDF transform of functional keywords with locality sensitive hashing (LSH), detailed in Appendix A.1.8.

###### Transformer Trunk

A standard Transformer (62, 63) architecture processes the post-tokenized inputs. Geometric Attention (Algorithm 6 and Fig. S2) directly processes structural coordinates as input. Model outputs are logits over token space, and can be sampled to obtain outputs described in Appendix A.1.5.2. The overall architecture is diagrammed in Fig. S1.

###### Decoder

Most tracks can be naively decoded into tokens detailed in Appendix A.1.3. Structure tokens must be decoded with a model—we use a 700M parameter transformer model to do this, trained post-hoc (Appendix A.1.7.2). The decoder uses sequence tokens and structure tokens to directly predict coordinates, pTM, and pLDDT (64). Function tokens are decoded using a small 3-layer transformer, trained post-hoc to invert the LSH quantization procedure (Appendix A.1.8.2.1).

##### A.1.3 Tokenization

All tracks are represented as a sequence of tokens, with tokens specified in each amino acid position. During tokenization, special beginning-of-sequence (BOS) and end-of-sequence (EOS) tokens are prepended and appended.

###### Sequence

Protein sequences are tokenized as the 20 canonical amino acids, plus BOS, EOS, mask, pad, unknown.

We keep four non-standard amino acids as in Lin et al. (5), B - Asparagine, U - selenocysteine, Z - glutamic acid, and O - ornithine. This totals to 29 tokens.

###### Structure

Structure tokenization is described in Appendix A.1.7.1. ESM3 uses a codebook size of 4096 with 4 special tokens - EOS, BOS, mask, and pad.

###### Secondary Structure

Secondary structure is taken to be the canonical 8-class tokens (65), with unknown and mask, for a total of 10 tokens. The mask token is forced to be the 0-vector during embedding.

###### SASA

The continuous values representing SASA are tokenized by discretization into a fixed set of 16 bins. SASA bin boundaries were chosen by computing SASA on 100 random structures and ensuring an equal number of residues belong in each bin. Unknown and mask are used for a total of 18 tokens. The mask token is forced to be the 0-vector during embedding.

###### Function annotations

We tokenize function annotations as bags of keywords, described in Appendix A.1.8. Keywords are quantized using LSH into 8 tokens per residue, each of which can be one of 255 tokens. There are three special tokens, empty set, no-annotation, and mask. Again, the mask token is forced to be the 0-vector during embedding.

###### Residue annotations

InterPro annotations are tokenized as a multi-hot feature vector (1478 dimensions) over possible InterPro labels (36). Input annotations are limited to a maximum of 16. When annotations are not present, we enforce that the 0-vector is added.

##### A.1.4 ESM3 Inputs and Forward Pass

As mentioned above, ESM3 can take several tracks, all of which are optionally disabled via masking. In the following, we concisely denote the inputs to ESM3 as

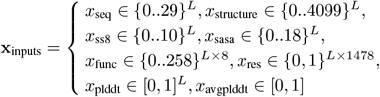

We now present the high level algorithm for a forward pass of ESM3:

**Figure S1.**
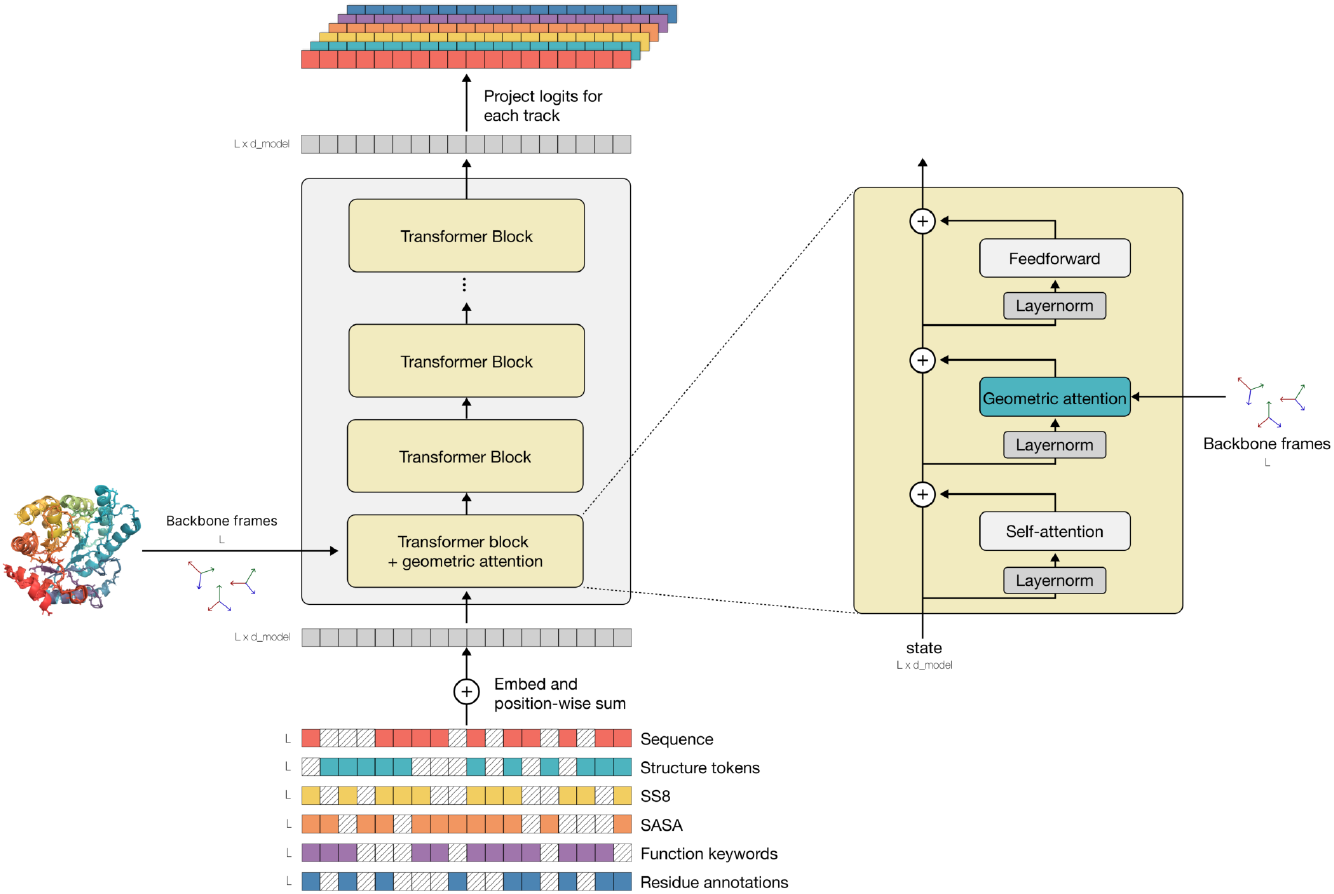
The ESM3 architecture. ESM3 is a masked language model that reasons over protein sequence, structure, and function, each of which are represented as token tracks at the input and output. Tokens are embedded and summed at the input to a transformer stack. The first block (expanded on the right) contains an additional geometric attention layer for processing atomic coordinate inputs. During training, random masks are sampled and applied to each track. Masked token positions are predicted at the output.

###### Algorithm 1

`esm3_forward`

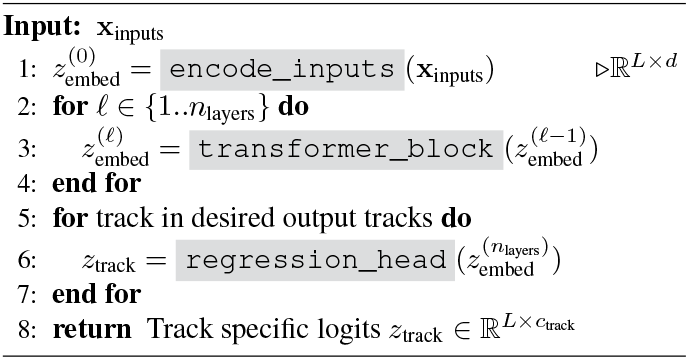

In the next few sections, we detail each component.

##### A.1.5 Transformer

Our network is based on the transformer architecture (62), incorporating several subsequent improvements: We use Pre-LN instead of Post-LN (63), rotary embeddings (66) instead of absolute positional embeddings, and we replace ReLU non-linearity with SwiGLU (67). The hidden dimension is set to approximately ^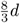^, rounded to the nearest multiple of 256 for training efficiency. No biases are used in linear layers or layer norms, as suggested by PaLM (68). We have observed through the literature and in internal experiments that these architecture changes improve the stability and performance of models.

A core architectural modification we make is the insertion of the Geometric Attention sub-layer in the first block of the network only (Algorithm 2, line 3).

###### Algorithm 2

`transformer_block`

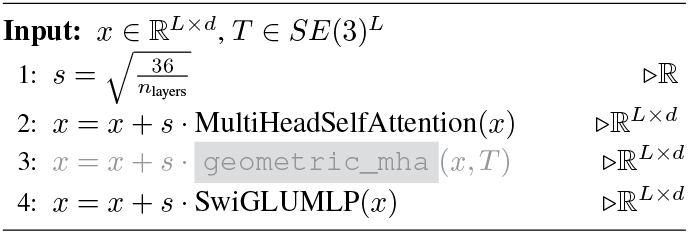

ESM3-small (1.4B) is a 48-layer network, while ESM3-medium (7B) has 96 layers, and ESM3-large (98B) has 216 layers. We experimented with different width-to-depth ratios and observed higher returns for depth than width. Prior work also demonstrates that modalities like ours benefit more from deeper networks (69, 70). Detailed network specifications can be found in Table S1.

###### A.1.5.1 Embedding

There are 7 unique input tracks to ESM3: (a) sequence (amino acid tokens), (b) structure coordinates, (c) structure tokens, (d) 8-class secondary structure labels (SS8), (e) quantized solvent-accessible surface area (SASA) values, (f) function keyword tokens and (g) residue (InterPro) annotation binary features.

There are two additional tracks used during pretraining only: (h) per-residue confidence (pLDDT) and (i) averaged confidence (pLDDT). At inference time, these values are fixed, and these tracks are equivalent to adding a constant vector *z*_plddt_.

Structure coordinates are parsed through the Geometric Attention and are not embedded.

For keyword-based function tokens, each of the eight integers per residue is converted to a “sub-embedding” (Algorithm 3 line 5), then concatenated to form the per-residue embedding (Algorithm 3 line 6). For InterPro residue annotations, the inputs are multi-hot. To create an embedding vector, we sum the embeddings for each of the “on” features (equivalent to the matrix-multiply on Algorithm 3 line 7).

The largest model, 98B, has an additional taxonomy track detailed in Appendix A.1.9.2, only enabled in the final 30K steps of pretraining.

The embeddings are all summed as input to the first layer in the network architecture.

###### Algorithm 3

`encode_inputs`

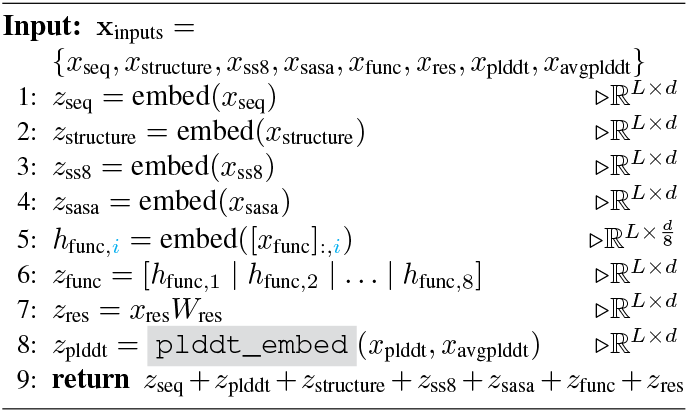

###### A.1.5.2 Logits

We use a `regression_head` to take in *d* dimensional last layer hidden features and produce *c*_track_-dimensional logits for each of the tracks, where *c*_track_ corresponds to the size of the vocabulary per track. Note that for the keyword function tokens, we produce *c*_func_ × 8 logits, and softmax over each of the 8 independently when calculating the loss.

**Table S1.** Parameter details for different model configurations.

| Params | $n_{\text{layers}}$ | $d_{\text{model}}$ | $d_{\text{head}}$ | Context length | Learning rate | Warmup steps | Batch size in tokens | Num steps | Total tokens | FLOPs |
| --- | --- | --- | --- | --- | --- | --- | --- | --- | --- | --- |
| 1.4B | 48 | 1536 | 64 | 2048 | 4.0e-4 | 5K | 1,572,864 | 50K | $\sim 80\text{B}$ | $6.72 \times 10^{20}$ |
| 1.4B | 48 | 1536 | 64 | 2048 | 4.0e-4 | 5K | 1,572,864 | 200K | $\sim 320\text{B}$ | $2.7 \times 10^{21}$ |
| 7.7B | 96 | 2560 | 128 | 2048 | 2.5e-4 | 5K | 3,932,160 | 140K | $\sim 550\text{B}$ | $2.47 \times 10^{22}$ |
| 98.5B | 216 | 6144 | 128 | 2048 | 1.0e-4 | 20K | 4,194,304 | 430K | $\sim 1.8\text{T}$ | $1.07 \times 10^{24}$ |

###### Algorithm 4

`regression_head`

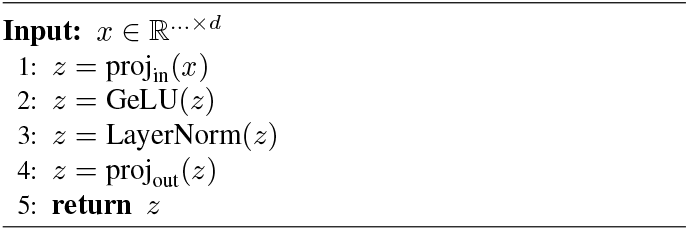

Except for structure coordinates, we output predictions for each of the tracks detailed in Appendix A.1.5.1: (a) sequence, (b) structure tokens, (c) SS8, (d) quantized SASA, (e) function keyword tokens and (f) residue (InterPro) annotation binary features.

Except for the multi-hot residue annotations, all other tracks are predicted as a categorical distribution over possible tokens.

##### A.1.6 Geometric Attention

As mentioned in Appendix A.1.5.1, ESM3 processes structural information in two independent ways:

###### Geometric Attention

Described in Algorithm 6, this leverages fine-grained 3D information via conditioning on atomic coordinates of backbone atoms. Coordinates are only used as model inputs.

###### Structure Tokens

Described in Appendix A.1.7, structure tokens enable faster learning due to rich local neighborhood semantics being compressed into tokens. Structure tokens are generally used as model outputs.

Geometric attention enables high-throughput encoding of protein structures. Protein backbone structure can be represented by the relative distance and orientation of frames defined by each residue’s backbone coordinates. Reasoning over the relative orientation of frames is important to capture the local backbone geometry when only partial structure is provided. Geometric attention is an *SE*(3) invariant all-to-all attention mechanism which reasons over the relative distances and orientations of all defined frames in the input (Fig. S2). Because this attention operation can be realized using the same computational primitives as attention, it is readily scalable.

We first provide an overview of frames, and then describe how geometric attention uses them:

###### A.1.6.1 Frames

Frames are representations that encapsulate the positions and orientations of each residue in the protein. We use a formulation similar to Ingraham et al. (71). Each frame *T* ∈ *SE*(3) consists of a rotation matrix R ∈ *SO*(3) and a translation vector t ∈ ℝ^3^.

###### Definition

A frame *T*_*i*_ for residue *i* is defined as:

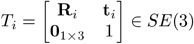

where R_*i*_ ∈ *SO*(3) and t_*i*_ ∈ ℝ^3^.

**Rotation Matrix:** The rotation matrix R_*i*_ for residue *i* is composed of three 3-dimensional vectors 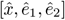:

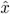 and *ê*_1_ are orthogonal unit vectors on the *N* − *C*_*α*_ − *C* plane.

*ê*_2_ is a unit vector perpendicular to both 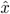 and *ê*_1_.

This matrix rotates vectors to a local coordinate system where the *N* −*C*_*α*_ −*C* plane for the corresponding residue spans the *xy* plane.

**Translation Vector:** The translation vector t_*i*_ specifies the position of the residue’s *C*_*α*_.

**Transformation:** To transform a point p ∈ ℝ^3^ from the local frame of residue *i* to the global coordinate system, the following equation is used:

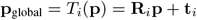

**Inverse Transformation:** To transform a point p_global_ ∈ ℝ^3^ from the global coordinate system back to the local frame of residue *i*, the following equation is used:

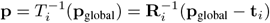

**Figure S2.**
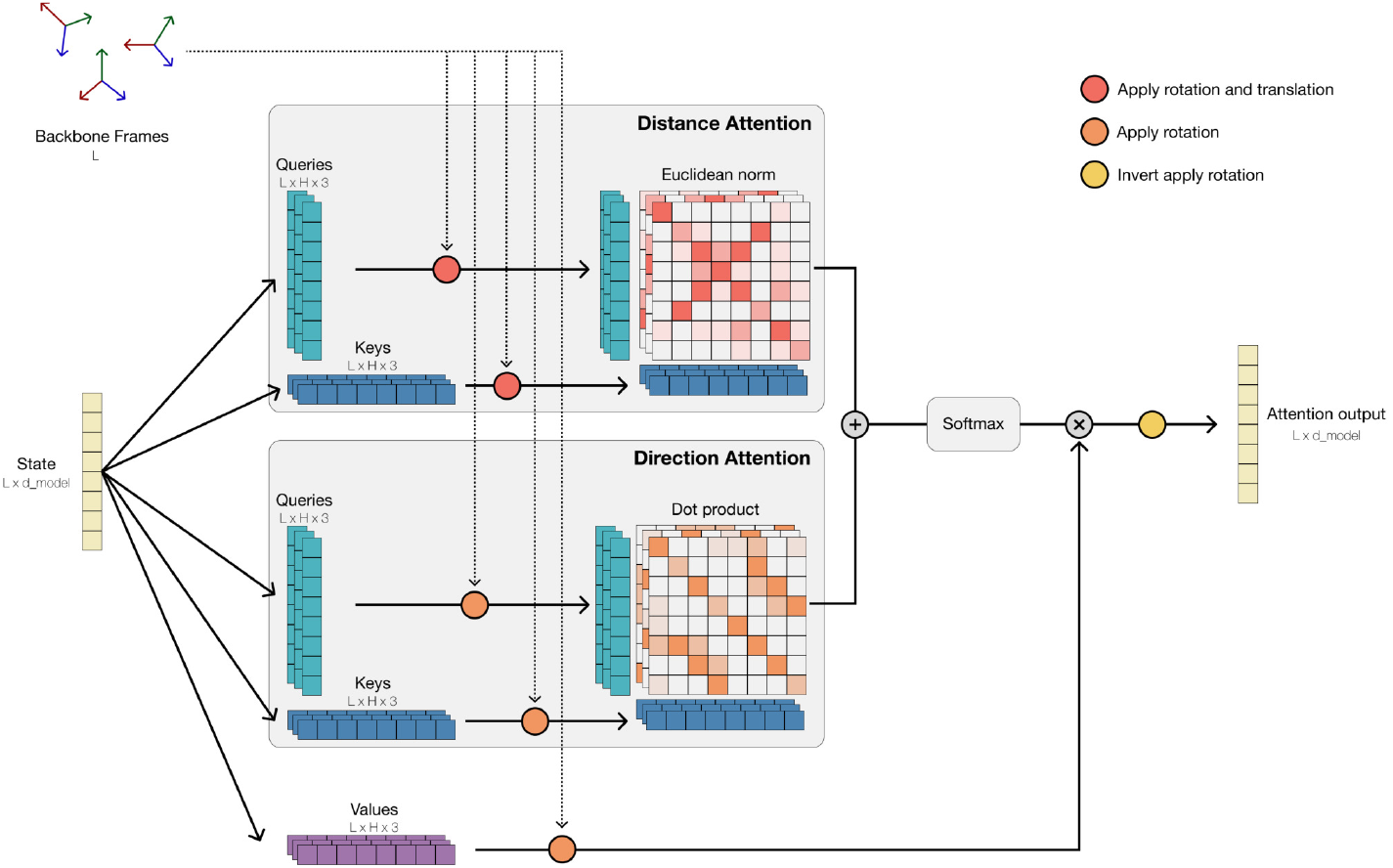
Geometric attention. Geometric attention is an SE(3) invariant all-to-all attention mechanism where the attention score matrix is a weighted sum of two terms: (1) the pairwise distances between queries and keys rotated and translated by their respective backbone frames, and (2) the pairwise dot products between queries and keys rotated by their respective backbone frames. This attention mechanism encodes structural information with throughput comparable to the standard attention operation in transformers.

To create frames, all we require is a translation vector 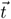, and two vectors 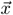 and 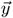 defining the local *xy* plane *after* conversion to global coordinates, from which the frame *T* can be calculated with the standard Gram-Schmidt algorithm:

###### Algorithm 5

`gram_schmidt`

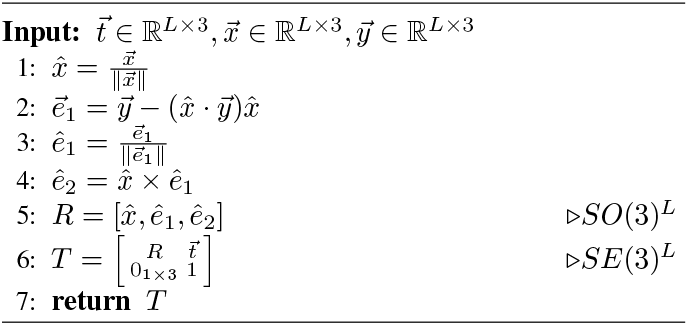

We construct frames such that the *C*_*α*_ is at the origin of the frame 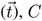 on the negative x-axis 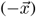, and *N* is on the *xy*-plane.

###### A.1.6.2 Geometric Self-Attention

Algorithm 6 details the Geometric Self-Attention layer. It can be efficiently implemented using similar ideas as FlashAttention (72). It is used twice in our system: in the VQ-VAE encoder for structure tokens (Appendix A.1.7.1), and in the first layer of ESM3.

Unlike regular self-attention, which only operates on per-residue embeddings, Geometric Attention incorporates the per-residue frames *T* to integrate geometric information in a rotation and translation invariant way. The process of forming the attention matrix *A* is as follows:

1. **QKV Projections:** Two sets of keys and queries (*Q*_*r*_, *K*_*r*_) and (*Q*_*d*_, *K*_*d*_), along with *V*, all with shapes ∈ ℝ^*L×h×*3^ are linearly projected from layer input *X. L* is the sequence length, *h* is the number of heads.
2. **Convert QKV to global frame:** Each of the queries, keys and values are initially assumed to be in the local frame of their corresponding residue.
  a. **Convert to Global Rotational Frame:** We convert each of the vectors in *Q*_*r*_, *K*_*r*_, *V* from their local frame (where the *xy* plane is the *N* −*C*_*α*_ −*C* plane for each residue) to a *global* rotational frame (where the *xy* plane is aligned for all residues) by applying R_*i*_ (Algorithm 6, lines 3, 4).
  b. **Convert to Global Distance Frame:** We convert each of the vectors in *Q*_*d*_, *K*_*d*_ from their local frame to a *global* frame by applying *T*_*i*_ (Algorithm 6, lines 5, 6).
3. **Direction Attention:** The pairwise, per-head *h* rotational similarity *R* between keys *i* and queries *j* is calculated using the dot product 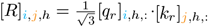_,:_. This is equivalent to the cosine distance between projected points.
4. **Distance Attention:** The pairwise, per-head *h* distance similarity *D* between keys *i* and queries *j* is computed using the *L*_2_ norm of the difference 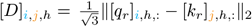.
5. **Scale Factor:** *R* and *D* are scaled per-head with learned scalars 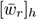 and 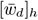, respectively, where 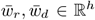. We use the softplus function to transform weights into [0, ∞)^*h*^. This scaling allows certain heads to specialize in attending via distance or direction attention.

###### Algorithm 6

`geometric_mha`

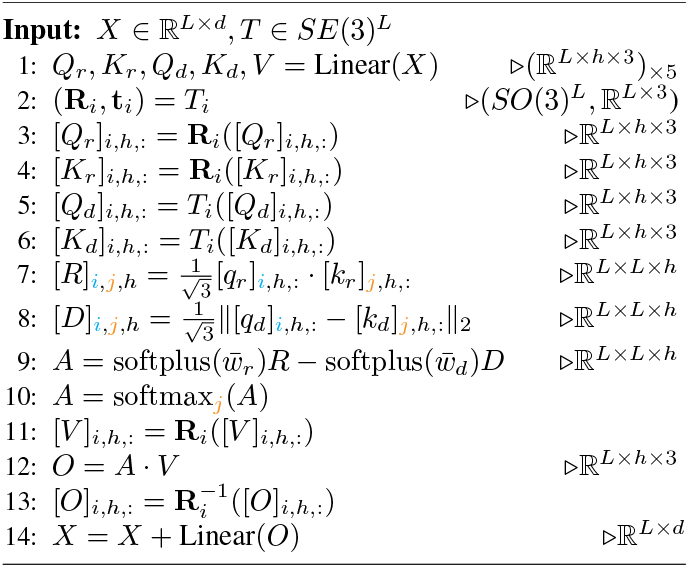

In the Geometric Self-Attention layer of ESM3, partially or fully masked coordinates can be input. Undefined coordinates are handled by masking keys and zeroing the attention output wherever coordinates are missing.

##### A.1.7 Structure Tokenizer

Each residue is associated with one of 4,096 structure tokens (+4 special tokens), designed to provide a rich, learned representation of its local neighborhood. Prior works have show structure tokens can be used for search (41), to improve representation learning (73), and for generation (16), but we trained our own tokenizer to optimize for structure generation. The tokens are generated with a VQ-VAE encoder, with a corresponding decoder to enable decoding of generated tokens back to 3D coordinates.

###### A.1.7.1 Encoder

The VQ-VAE encoder *f*_enc_ consists of two geometric attention blocks (Transformer blocks, but self-attention replaced with `geometric_mha`) with an embedding width of 1024 and 128 geometric heads per geometric attention layer.

The VQ-VAE encoder reasons over the backbone frames and the relative sequence position of residues in the local structure. Relative sequence positions are encoded through a learned positional embedding. Sequence positions are determined relative to the query residue (i.e., if the query residue has residue index 56, then the residue in index 58 has a +2 sequence position). Relative sequence positions are clamped to +/-32 before encoding, meaning long-range contacts share sequence positional embeddings. Relative sequence positional embeddings define the initial encoder state *N*, and has shape *L*× 16× *d* (Algorithm 7, line 4). Note that this means the input to the VQ-VAE encoder is purely structural: no sequence (amino acid), function or other information is used here. Furthermore, each neighborhood is processed completely independently; for each residue, the encoder only uses the information of its 16 nearest neighbors.

Geometric attention blocks operate similar to Transformer blocks in that they transform a state according to an attention operation (`geometric_mha`) and feedforward network (SwiGLU MLP). As such, the output has the same shape as the input. In this case, this means that the encoder outputs 16 latents per residue. However, we want to learn a single token, i.e., a single latent per residue, hence we take the embedding corresponding to the query residue position *N*_:,0,:_.

The process of generating structure tokens (Algorithm 7) from the full 3D coordinates of the protein then is as follows:

1. **Local Neighborhood:** For each residue, obtain the indices *N*_idx_ ∈ {0..*L*−1}^*L×*16^ of the 16 nearest residues (as measured by *C*_*α*_ distance). The first of the 16 neighbors is always the residue itself. We also obtain the frames for each residue in a local neighborhood with *T*_knn_.
2. **Embed Neighbors:** Embed the relative distance in sequence space for each neighbor, ∆*i* = clamp(*N*_idx_ − *i*, −32, 32) to form *N* ∈ R^*L*×16×*d*^.
3. **Encode:** Pass *N* through a shallow encoder *f*_enc_ consisting of 2 Transformer blocks, with regular multi-head self-attention swapped with `geometric_mha`. The attention is unmasked, all-to-all over the entire neighborhood.
4. **Quantize:** Extract the first element *N*_:,0,:_ from the neighborhood, which corresponds to the residue itself. Project it linearly, and quantize by replacing with the nearest vector in a codebook. This yields the structure token per residue.

###### Algorithm 7

`structure_encode`

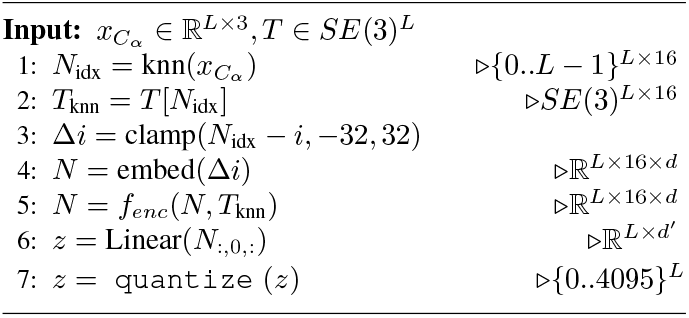

###### 1.7.1.1 Codebook Learning

`quantize` transforms the *L* latents into *L* discrete tokens. Since the VQ-VAE was initially proposed (74), numerous approaches and tricks have been developed to address issues with poor codebook utilization and unstable training. We chose to learn the codebook as an exponential moving average of encoder outputs (74–76). To improve codebook utilization, unused codes are re-initialized to encoder outputs.

###### 1.7.1.1 Parallel Encoding

To improve training and inference efficiency, we encode all local structure graphs within a protein in parallel. In practice, this means that given a batch of *B* proteins with average sequence length *L*, then the inputs to the structure encoder will have shape *BL* × 16 × *d*.

###### A.1.7.2 Decoder

While the encoder independently processes all local structures in parallel, the decoder *f*_dec_ attends over the entire set of *L* tokens to reconstruct the full structure. It is composed using a stack of bidirectional Transformer blocks with regular self-attention.

As discussed in Appendix A.1.7.3, the VQ-VAE is trained in two stages. In the first stage, a smaller decoder trunk consisting of 8 Transformer blocks with width 1024, rotary positional embeddings, and MLPs is trained to only predict backbone coordinates. In the second stage, the decoder weights are re-initialized and the network size is expanded to 30 layers, each with an embedding dimension of 1280 (∼600M parameters) to predict all atom coordinates.

The exact steps to convert structure tokens back to 3D allatom coordinates using the decoder is provided in Algorithm 8 and detailed as follows,

1. **Transformer:** We embed the structure tokens and pass them through a stack of Transformer blocks *f*_*dec*_ (regular self-attention + MLP sublayers, no geometric attention).
2. **Projection Head:** We use a projection head to regress 3 3-D vectors per residue: a translation vector 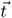, and 2 vectors 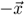 and 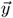 that define the *N* −*C*_*α*_ −*C* plane per residue *after* it has been rotated into position. This head also predicts the unnormalized sine and cosine components of up to 7 side chain torsion angles.
3. **Calculate *T* :** We use `gram_schmidt` to convert 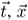, and 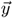 into frames *T* ∈ *SE*(3)^*L*^.
4. **Calculate *T***_**local**_: We normalize the sine and cosine components and convert them to frames *T*_local_ ∈ *SE*(3)^*L*×7^ corresponding to rotations around the previous element on the side chain.
5. **Compose Frames:** We compose each element of *T*_local_ with its predecessors on a tree rooted at *T* to form *T*_global_ ∈ *SE*(3)^*L*×14^, corresponding to the transformations needed for each heavy atom per residue in atom14 representation.
6. **Apply Frames:** We then apply the frame to the 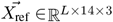 coordinates in a reference frame, to rotate and transform each residue into their final positions.

###### Algorithm 8

`structure_decode`

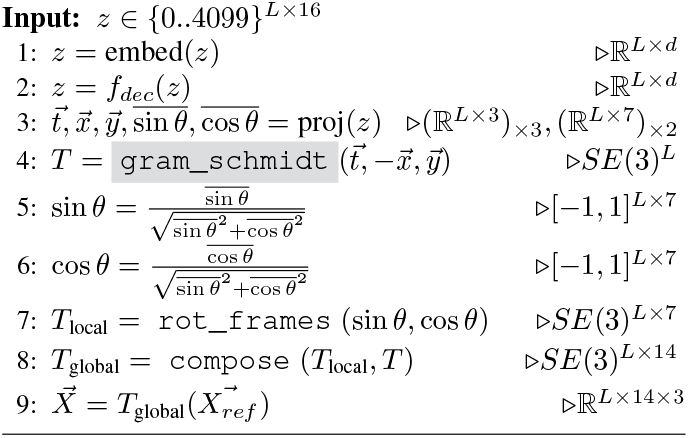

###### A.1.7.3 Training

When using a VQ-VAE to learn discrete representations which maximize reconstruction quality, it is common to train in the autoencoder in two stages (77). In the first stage, the encoder and codebook is learned with a relatively small and efficient decoder. In the second stage, the encoder and codebook are frozen and a larger or otherwise more computationally expensive decoder is trained to maximize reconstruction quality. We follow this two-stage training approach for the structure tokenizer. The VQ-VAE is fully trained before ESM3 pretraining, with the encoder and decoder kept frozen during ESM3 training.

###### A.1.7.3.1 Stage 1

The VQ-VAE is trained for 90k steps on a dataset of single chain proteins from the PDB, AFDB, and ESMAtlas. We use the AdamW optimizer (Loshchilov et al. 2017) with learning rate annealed from 4e-4 according to a cosine decay schedule. Proteins are cropped to a maximum sequence length of 512. Five losses are used to supervise this stage of training. The geometric distance and geometric direction losses are responsible for supervising reconstruction of high quality backbone structures.

Additionally, binned distance and direction classification losses are used to bootstrap structure prediction but are ultimately immaterial to reconstruction. We have found that these structure prediction losses formulated as classification tasks improve convergence early in training. To produce these pairwise logits, we use a `pairwise_proj_head`, that takes x ∈ℝ^*L*×*d*^ and returns logits 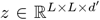. It works as follows:

###### Algorithm 9

`pairwise_proj_head`

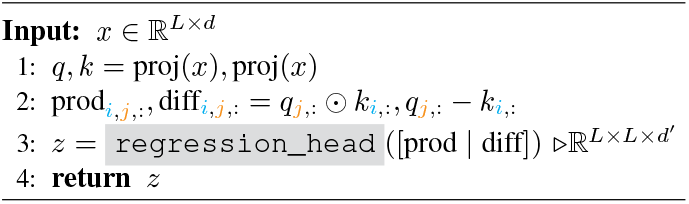

Finally, an inverse folding token prediction loss (i.e., a crossentropy loss between predicted sequence and ground truth sequence) is an auxiliary loss used to encourage the learned representations to contain information pertinent to sequence-related tasks.

The five losses are covered in detailed as follows:

1. **Backbone Distance Loss:** Compute the pairwise *L*_2_ distance matrix for the predicted and true coordinates of the 3 backbone atoms (*N, C*_*α*_, *C*). Let *D*_pred_, *D* ∈ ℝ^3*L*×3*L*^. Compute (*D*_pred_ −*D*)^2^, clamp the maximum error to (5 Å)^2^, and take the mean.

#### Algorithm 10

`backbone_distance_loss`

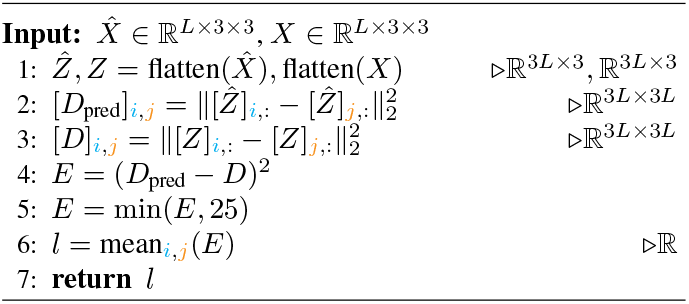
2. **Backbone Direction Loss:** Compute six vectors for both predicted and ground truth coordinates for each residue:
  a. *N* → *C*_*α*_
  b. *C*_*α*_ → *C*
  c. *C* → *N*_next_
  d. 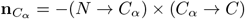
  e. n_*N*_ = (*C*_prev_ → *N*) × (*N* → *C*_*α*_)
  f. n_*C*_ = (*C*_*α*_ → *C*) × (*C* → *N*_next_)
  Compute the pairwise dot product, forming *D*_pred_, *D* ∈ ℝ^6*L*×6*L*^. Compute (*D*_pred_ *D*)^2^, clamp the maximum error to 20, and take the mean. In algorithm form (with `compute_vectors` computing the six vectors described above):

#### Algorithm 11

`backbone_direction_loss`

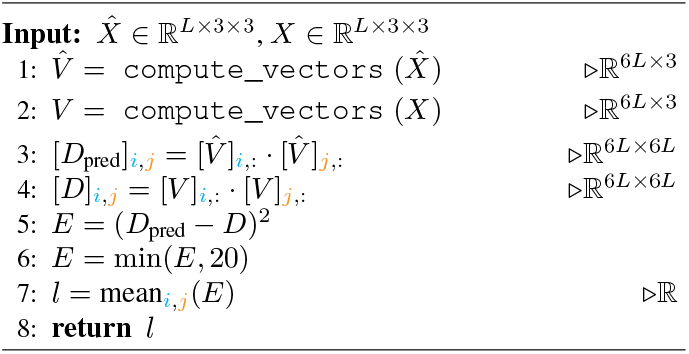
3. **Binned Direction Classification Loss:** This loss captures a coarser similarity between ground truth and predicted orientations to stabilize early training. It uses the last layer representations of the decoder, not the predicted coordinates. The process is as follows:
  a. **Unit vectors:** Compute three vectors per residue from ground truth coordinates: *C*_*α*_ → *C, C*_*α*_ → *N*, and 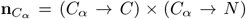, and normalize them to unit length.
  b. **Dot Products:** Compute pairwise dot products between each pair of vectors for all residues, forming *D* ∈ [−1, 1]^*L×L*×6^. Bin the dot products into 16 evenly spaced bins in [−1, 1], forming classification labels *y* ∈ {0..15}^*L*×*L*^.
  c. **Pairwise Logits:** Pass the final layer representations of the decoder *h* ∈ ℝ ^*L*×*d*^ through a `pairwise_proj_head` to obtain logits *z* ∈ ℝ ^*L*×*L*×6×16^.
  d. **Cross Entropy:** Calculate cross-entropy loss using the labels *y* from the ground truth structure and the logits *z*, and average over all *L* ×*L* ×6 values.
4. **Binned Distance Classification Loss:** Similar to the Binned Direction Classification Loss, this loss bins the true distances between residues (specifically, their *C*_*β*_) to get ground truth targets and computes a crossentropy between these targets and pairwise logits. In detail:
  a. **Calculate** *C*_*β*_: Given the ground truth *N, C*_*α*_, and *C* coordinates, we compute the location of *C*_*β*_:
    i. Obtain the three vectors *N* → *C*_*α*_, *C*_*α*_ → *C*, and n = (*N* → *C*_*α*_) × (*C*_*α*_ → *C*).
    ii. Define the following scalars:

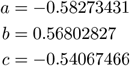
    iii. Compute the location of *C*_*β*_ using the formula:

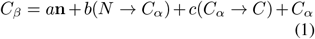
  b. **Pairwise** *C*_*β*_ **distances:** Compute an *L* × *L* pairwise distance matrix of the *C*_*β*_, and bin them into one of 64 bins, with lower bounds [0, 2.3125^2^, (2.3125 + 0.3075)^2^, …, 21.6875^2^], forming the labels *y* ∈ {0..63}^*L*×*L*^.
  c. **Pairwise logits:** Pass the final layer representations of the decoder *h* ∈ ℝ ^*L*×*d*^ through a `pairwise_proj_head` to obtain the logits *z* ∈ ℝ ^*L*×*L*×64^.
  d. **Cross Entropy:** Calculate the cross-entropy using the labels *y* computed from the ground truth structure and the logits *z*, then average over all *L* × *L* values.
5. **Inverse Folding Loss:** Pass final layer representations of the decoder through a regression head to produce logits *z*. Using ground truth residues as labels *y*, compute cross-entropy for the classification task of predicting residues from final layer representations.

###### A.1.7.3.2 Stage 2

In the second stage of VQ-VAE training, the encoder and codebook are frozen and a new, deeper decoder is trained. This second stage of training has multiple purposes. First, a larger decoder improves reconstruction quality. Second, augmented structure tokens from ESM3 are added to enable learning pAE and pLDDT heads. Third, we add sequence conditioning and train with all-atom geometric losses to be able to decode all-atom protein structures. Fourth, we extend the context length of the decoder to be able to decode multimers and larger single chain proteins.

Training data for stage 2 consists of predicted structures in AFDB and ESMAtlas, as well as single chains, multimers, and antibody-antigen complexes from the PDB. Sequence conditioning was added to the decoder via learned embeddings which are summed with structure token embeddings at the input to the decoder stack.

The structure token decoder was trained in three stages: 2A, 2B, 2C detailed in Table S2. The purpose of stage 2A is to efficiently learn decoding of all-atom structures. Enhanced training efficiency is achieved by keeping a short context length and omitting the pAE and pLDDT losses, which are both memory-consuming and can be in competition with strong reconstruction quality. In stage 2B, we add the pAE and pLDDT losses. These structure confidence heads cannot be calibrated unless structure tokens are augmented such that ESM3-predicted structure tokens are within the training distribution. To this end, for stages 2B and 2C we replace ground truth structure tokens with ESM3-predicted structure tokens 50% of the time. In stage 2C, we extend context length to 2048 and upsample experimental structures relative to predicted structures.

1. **All-atom Distance Loss:** We generalize the Backbone Distance Loss to all atoms by computing a pairwise *L*_2_ distance matrix for all 14 atoms in the atom14 representation of each residue. This results in *D*_pred_, *D* ∈ ℝ^14*L*×14*L*^. The rest of the computation follows as before: (*D*_pred_ − *D*)^2^, clamping to (5 Å)^2^, and taking the mean, while masking invalid pairs (where any atom14 representations are “empty”).
2. **All-atom Direction Loss:** We extend the Backbone Direction Loss to all heavy atoms:
  a. Identify all covalent bonds between heavy atoms within each residue.
  b. For each atom with at least two covalent bonds, compute a normal (z-axis) vector to the plane spanned by its first two covalent bonds. This results in a set of z-axis and bond vectors.
  c. Compute all-to-all pairwise dot products between z-axis and bond vectors, forming *D*_pred_, *D* ∈ ℝ^*n*×*n*^. Compute (*D*_pred_ − *D*)^2^, clamp the max to 20, and take the mean.
3. **pLDDT Head:** Uses a Regression Head with 50 output classes (each capturing 0.02 units from 0 to 100). Predicted structures are compared to ground truth to calculate per-residue pLDDT values, which are supervised with cross-entropy loss.
4. **pAE Head:** Use a Pairwise Projection Head to produce 64 logits per residue pair ∈ ℝ ^*L*×*L*×*d*^, converting to probabilities *p* via softmax. Each probability corresponds to a bin representing 0.5 Å of positional error, with centers [0.25, 0.75, …, 31.25, 31.75].

**Computing Loss:**

a. Compute the pairwise distances between residues in both the predicted and ground truth structures, resulting in distance matrices *D*_pred_ and *D* ∈ ℝ ^*L*×*L*^.
b. Calculate the differences (*D*_pred_ − *D*).
c. Bin these differences into 64 bins, generating classification targets for the logits.
d. Compute the loss using cross-entropy between these targets and the logits.

**Computing pAE:** Multiply probabilities by bin centers and sum to obtain the expected positional error per residue pair, with values ∈ [0.25, 31.75].

**Computing pTM:** Additionally, the pairwise logits are used to compute the pTM (Predicted Template Modeling) score, as follows:

a. Compute *f*_*d*_ for sequence length *L* as:

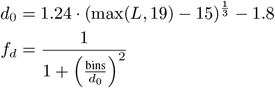
b. Compute pTM using previously computed probabilities *p*:

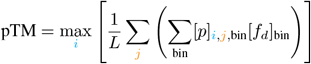

###### A.1.7.4 Evaluation

We evaluate the reconstruction quality of the structure tokenizer after stage 1 and stage 2 of training using a set of CAMEO, CASP14, and CASP15 proteins taken after the training cutoff date (Fig. S3). Both decoders consistently reach backbone RMSD < 1Å, LDDT-CA > 0.98. The retraining of the structure token decoder results in substantial improvements in reconstruction quality across all test sets.

**Table S2.**
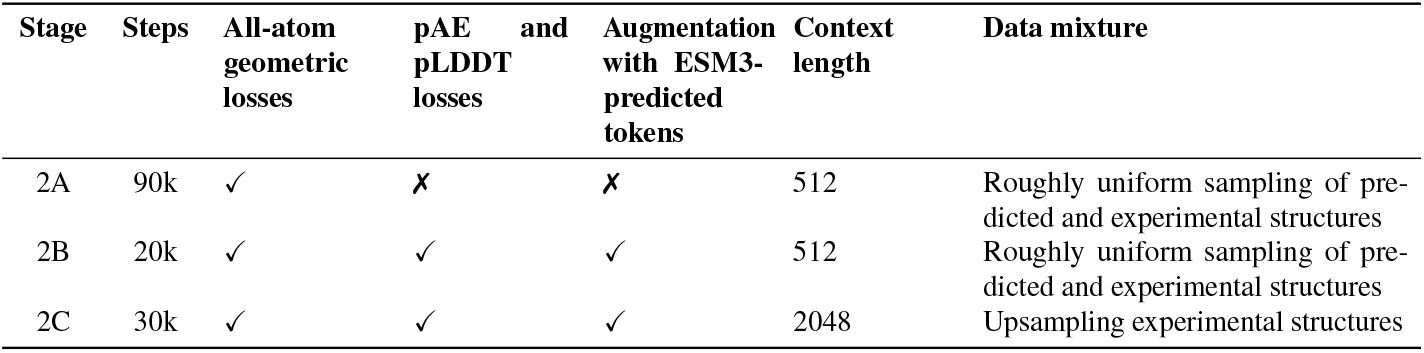
Training details for stage 2 training of an all-atom structure token decoder.

The stage 2 decoder, trained with an all-atom reconstruction loss and a sequence input, achieves strong all-atom reconstruction as well (Fig. S3C). We also visualize a random sample of backbone reconstructions on the CAMEO test set (Fig. S4A). Looking at the proteins with worse reconstruction quality, we find that long regions with few tertiary contacts, disordered regions, and unresolved coordinates can lead to inaccurate global orientation of structural elements, while local structure reconstruction remains largely error-free (Fig. S4B). This behavior can be explained by the fact that the tokenizer relies on tertiary contacts to resolve the global orientation of a residue.

We also investigate the vocabulary learned by the structure tokenizer by visualizing the local neighborhoods which map to the same learned structure token. We find that many structure tokens encode semantically coherent sets of local neighborhoods (Fig. S5A). However, a number of tokens appear to represent multiple local neighborhoods (Fig. S5B). While the meaning of a single token may be ambiguous, the high-fidelity reconstruction quality from the decoder suggests that it is able to disambiguate given surrounding context in the full set of structure tokens.

Fig. S6 indicates that pLDDT and pTM have good predictive power. We assess the calibration of the structure confidence heads on the CAMEO test set using structure tokens predicted by ESM3 7B. Most predictions for pLDDT lie along the diagonal, though there is a small bias towards more confident predictions. As pTM is a pessimistic estimator of the TM-score, we find that pTM is biased downwards. Anecdotally, we also find that pLDDT can be poorly calibrated for some generated sequences, particularly in alpha helical regions where it can be an overestimate.

##### A.1.8 Function Tokenization

ESM3 processes annotations of functional characteristics of proteins through two tracks: function tokens, and residue annotations. Both support input conditioning and output heads for generation. Appendix A.1.5.1 outlines how tokens are processed into the network. We further describe the creation of the tokens in this section.

###### A.1.8.1 Function Tokens

Function tokens are a dense semantic representation of functional characteristics of proteins derived from free-text descriptions of the InterPro and Gene Ontology (GO) terms at each residue. At training time, function tokens are produced from each protein’s InterPro annotations by a multi-step process illustrated in Fig. S7. At a high level:

1. For each residue, we gather free-text for each InterPro annotation via annotation term names, associated GO terms per annotation (via InterPro2GO mapping), and all ancestor GO terms. All residues with the same InterPro domain are assigned the same set of GO terms. We parse the free-text into counts from a vocabulary of 68,103 keywords. The vocabulary is composed of unigram and bigrams extracted from the free-text of all valid InterPro annotations (and their associated GO/ancestor GO terms) in our training datasets.
2. The keywords are converted to a sparse TF-IDF vector per InterPro annotation. During training, we also produce a corrupted version by dropping keywords at the protein level (i.e. the same keywords have their counts set to 0 across all residues) at a 15% probability per keyword.
3. To create a vector per residue from the per annotation vectors, we max pool the TF-IDF vectors for the annotations per residue. During training, we apply further corruption by dropping annotations at the protein level (i.e. the same annotations are removed from the max pool across all residues) at a 15% probability per annotation.
4. We then quantize each residue’s vector (a highly sparse vector with float entries) into a discrete representation suitable for input to the language model as tokens by applying a fixed series of 8 locality sensitive hashes (LSH), each with 8 hyperplanes.

**Figure S3.**
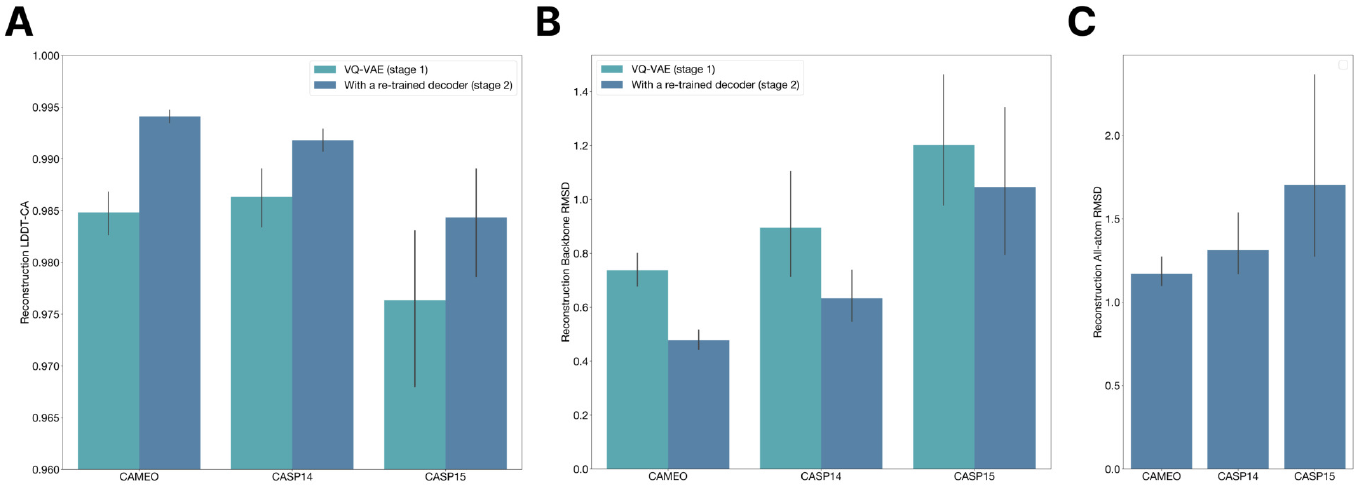
Structure tokenizer reconstruction quality. Reconstruction quality of the structure tokenizer after stage 1 and stage 2 of VQ-VAE decoder training evaluated on temporally held out CAMEO, CASP14, and CASP15. (A) Reconstruction LDDT-CA. (B) Reconstruction backbone RMSD. (C) All-atom reconstruction RMSD from the stage 2 decoder which additionally receives sequence input. Error bars represent 95% confidence intervals.

The result is a sequence of 8 tokens each ranging in value from 0 to 255 per-residue. We reserve a special token `<none>` to represent positions with an empty set of InterPro annotations. For proteins that lack any functional annotations, the tokens are filled with the `<pad>` token which has an embedding value fixed to all zeros. At test time, we can produce per residue vectors using the process described, or directly creating a TF-IDF vector with keywords.

During pretraining we use the corrupted versions of the function tokens at input, predicting the un-corrupted version function tokens at positions which have been masked. 90% of the time, the entire input is replaced with `<mask>`. The other 10% of the time, we replace all 8 tokens of selected residue with a `<mask>`, with the per-residue selection probability sampled from a cosine masking schedule per protein. The model has an output head which predicts each of the 8 function tokens in positions with `<mask>` as input, and is trained with a categorical cross entropy loss.

Function tokenization offers several key advantages as compared to simpler approaches (e.g., using a dedicated InterPro tag vocabulary). Encoding functional annotations with a generic functional keyword vocabulary supports flexible prompting of the model with combinations of keywords.

Function tokenization can also be viewed through the lens of data compression. This choice of representation reduces the input/output space from all possible InterPro combinations which would naively be represented by 35k bits, to a reduced space of 8 tokens x 8 bits / token = 64 bits. This also affords significant memory saving during pretraining by eliminating the need to perform multi-class multi-label binary classification.

###### A.1.8.2 Function Prediction

ESM3 is trained to predict all 8 function tokens, each spanning 256 possible values. To extract interpretable predictions of protein function from ESM3, we decode the predicted function tokens into function keywords using a seperately trained function token decoder.

###### A.1.8.2.1 Function Token Decoder

We train a 3-layer transformer model to learn the inverse map of the function tokenization process. The model takes as input the 8 function tokens representing the locality sensitive hash of function keywords. It outputs for each residue the binary-classification predictions predicting the presence of each function keyword, as well as predicting InterPro annotations from which the keywords originate. We find that unpacking the 8-bit LSH tokens into single-bit tokens improves training dynamics of the function token decoder. We train the function token decoder offline using combinations of InterPro tags from the UniRef annotated proteins. Since the function token vocabulary is fixed, the decoder is applied identically across different ESM3 model sizes.

###### A.1.8.2.2 Evaluation

To evaluate ESM3’s performance in predicting protein function, we compute Average Precision, a standard measure of information retrieval, using a validation set of proteins from UniRef and their associated InterProScan function annotations. We present results in Fig. S8.

**Figure S4.**
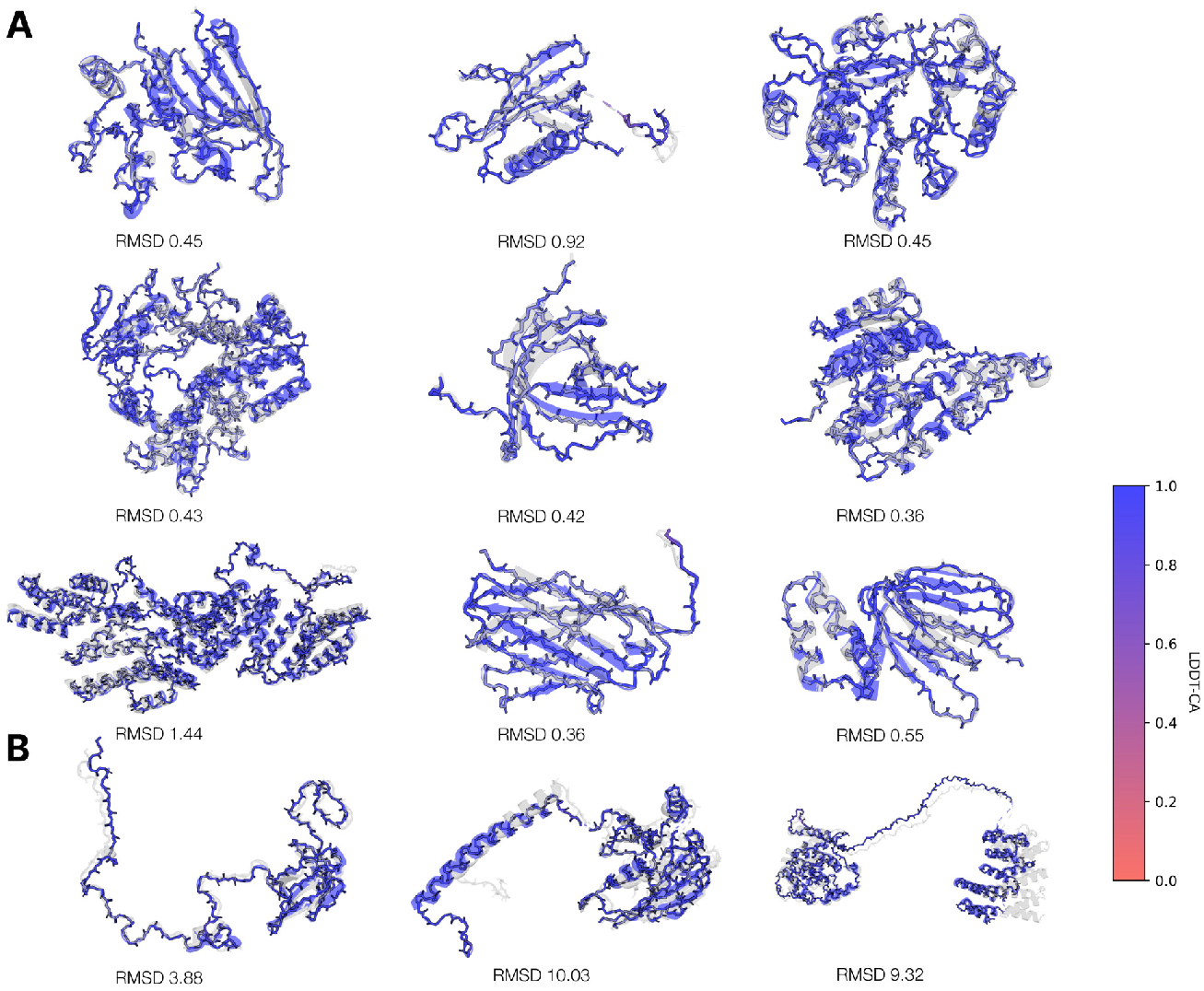
Visualization of structure tokenizer backbone reconstructions. (A) A random sample of reconstructions from the structure tokenizer on the test set. The vast majority of structures have near perfect backbone reconstruction. (B) A selection of the worst reconstructions in the test set. Long stretches of disordered regions, a lack of tertiary contacts, and unresolved coordinates can lead to inaccurate global orientation of structural elements, while local structure reconstruction remains largely error-free.

**Figure S5.**
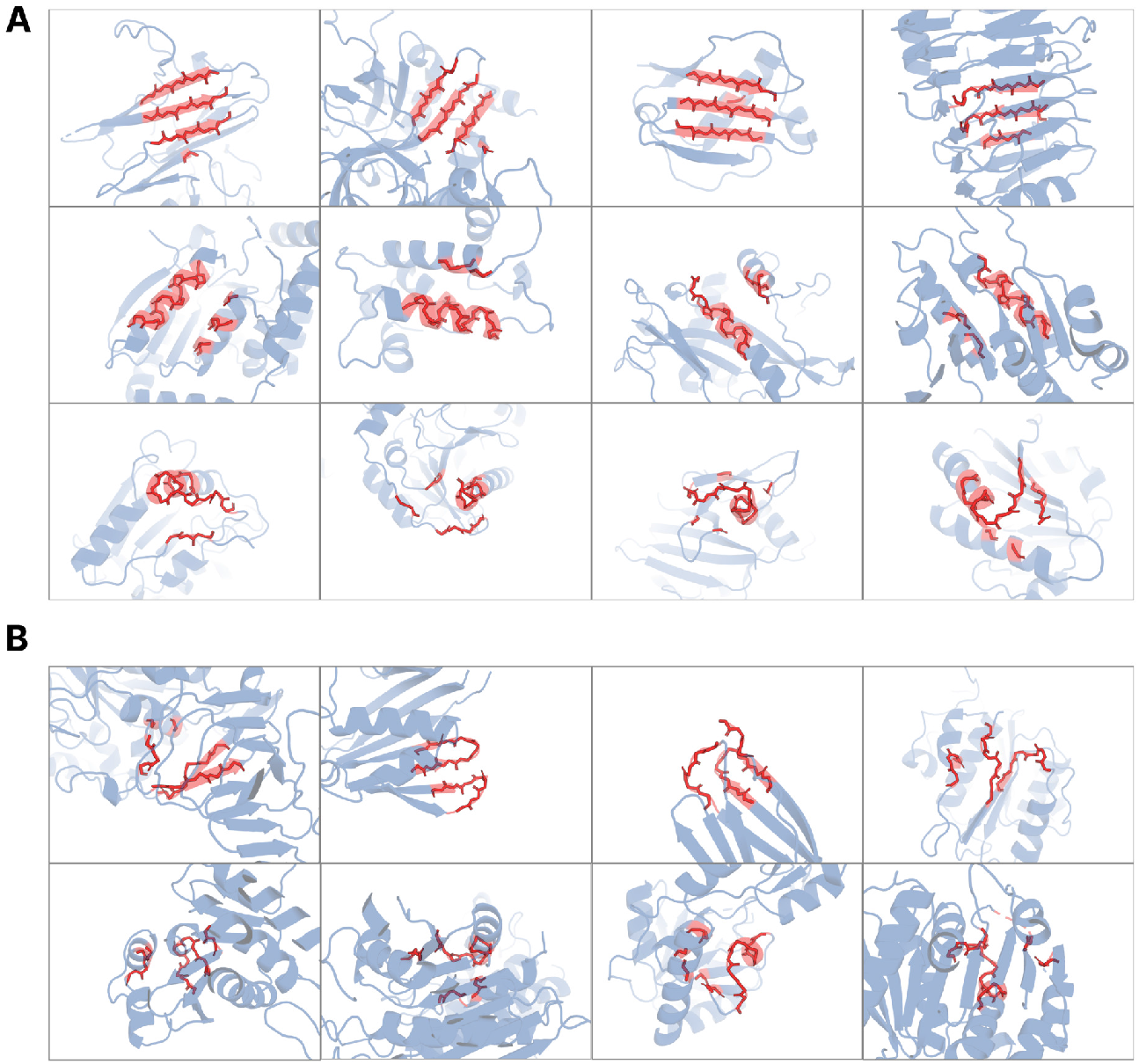
Visualization of local neighborhoods which map to the same learned structure token. The VQ-VAE encoder reasons over local structure neighborhoods (highlighted in red) which include the query residue and the 15 nearest neighbors in structure space. (**A**) Rows correspond to token indices 585, 59, and 3692 for top, middle, and bottom, respectively. Columns show different local structures mapping to the same token. (**B**) Some tokens represent multiple types of local neighborhoods. All local neighborhoods in B map to the same codebook index 3276. While the meaning of a single token may be ambiguous, the decoder is able to disambiguate given surrounding context in the full sequence of structure tokens.

**Figure S6.**
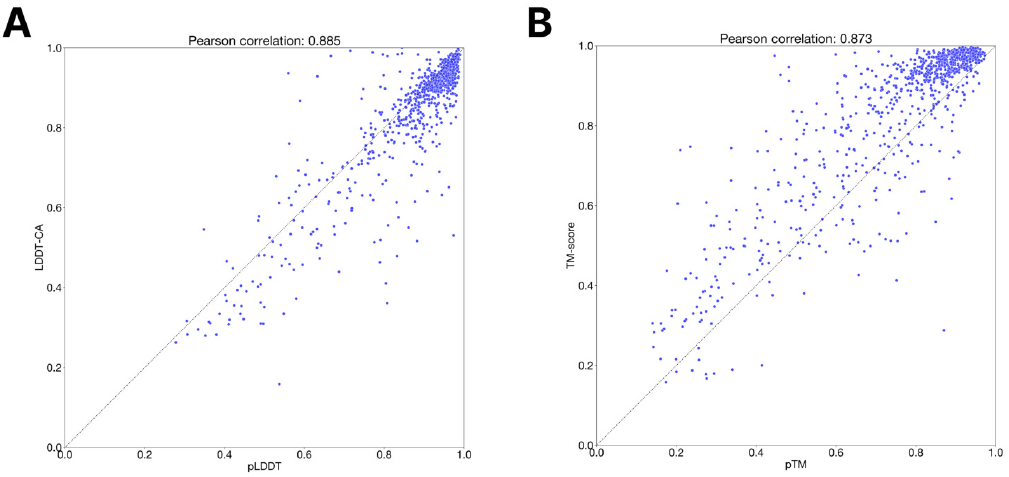
pTM and pLDDT calibration. Calibration of the structure token decoder pLDDT and pTM (using ESM3 7B as the structure token prediction model) on the CAMEO test set.

**Figure S7.**
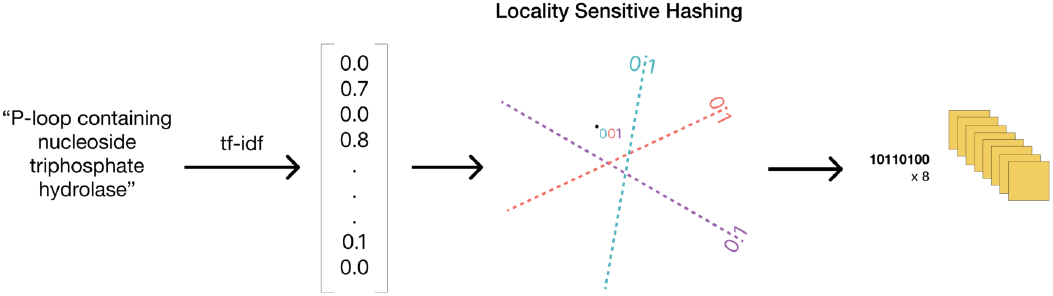
Schematic of function tokenization. The set of InterPro and GO descriptions of protein function are vectorized by a TF-IDF model and then hashed by a locality sensitive hash to produce 8 tokens each containing 8 bits.

**Figure S8.**
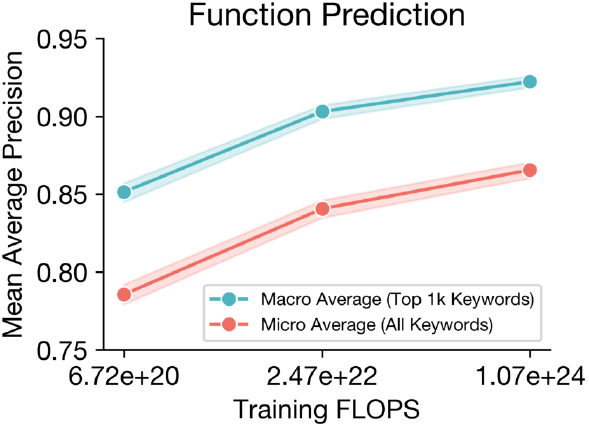
Function prediction benchmarking results. Mean Average Precision (mAP) for function keyword prediction. Predictions and labels are compared on a per-position basis to evaluate the model’s ability to localize site-specific functional attributes by keywords such as “active site”. We report mAP for the full keyword set (red) with a “micro” average because many keywords have few or no labels in the validation set. To report a “macro” average mAP we compute mAP for each of the top 1,000 most prevalent keywords in our evaluation set (discarding uninformative keywords such as “the”) and report a uniform average (blue). 95% confidence intervals are shown by shading.

###### A.1.8.3. Residue Annotations Track

Residue annotations label a protein’s sites of functional residues with a vocabulary of 1474 multi-hot labels emitted by InterProScan. To gather this data, we run InterProScan with databases (SFLD, CDD, PIR) on all cluster members in our UniRef and Mgnify datasets (seq-id 90 clustered). We take all unique residue annotation descriptions that occur in more than 1k proteins across all of UniRef90 and MGnify90, and deduplicate labels by punctuation and case insensitivity. We join these annotations into our UniRef, MGnify, AFDB, and ESMAtlas datasets for training.

As introduced in Appendix A.1.5.1, ESM3 has a track dedicated to processing residue annotations that supports input conditioning, and an output head for prediction and generation. The residue annotation labels for a protein are tokenized into a sequence of token-sets in length equal to the protein. At each position there is an unordered set of tokens representing the residue annotations present at that position. The tokens are input to ESM3 first through an embedding lookup followed by a sum over embeddings. The permutation invariance of the sum retains that the labels are represented to an unordered set as a model. The per-position embedding sums are then added onto the per-position sequence embedding before input into the first transformer block. Positions with no residue annotations are represented by a `<pad>` token which has an embedding fixed to zeros.

The residue annotations track has an output head which outputs a set of binary classification logits predicting for each position the presence or absence of each residue annotation in the vocabulary. We apply a masking procedure to partially/fully mask residue annotation labels, and train the output head with a binary cross-entropy loss function to reconstruct the full residue annotation. In pretraining, with 90% probability all residue annotations are masked, and otherwise we independently sample positions to mask with a square root schedule.

##### A.1.9. Other Tracks

###### A.1.9.1. Confidence Tracks

As mentioned in Appendix A.1.5.1, ESM3 has two additional tracks that are only used during pretraining, and only used as input (we do not have output heads predicting these values). The first is a per-residue pLDDT: for ground truth PDB structures, these values are all 1; for AlphaFold-DB/ESMFold structures, we use the provided pLDDT. We also provide an averaged pLDDT across all the residues when structure is provided (1 otherwise), with the average calculated before any tokens are masked.

This information allows the model to distinguish between gold-standard structures and computationally predicted ones; at inference time, we set these to 1 throughout, with the goal of producing structures better than the computational predictions used to pretrain the model. The embedding itself is straightforward, with the pLDDT values first having a radial basis function, followed by a Linear layer applied to them:

###### Algorithm 12

`rbf`

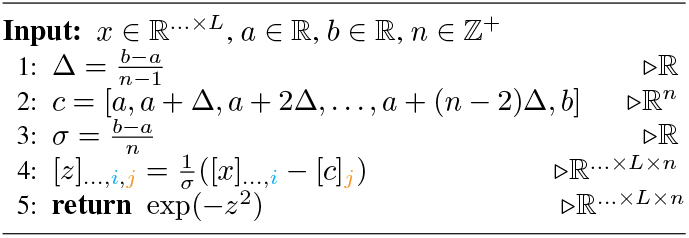

###### Algorithm 13

`plddt_embed`

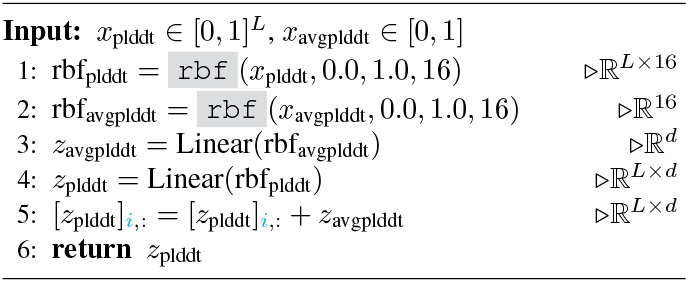

###### A.1.9.2. Taxonomy Track

The final 30,000 steps in the pretraining of the 98B variant of ESM3 includes a track for processing the taxonomic and species classification of the organism from which the protein sequence originates. For each protein, the taxonomic and species classifications are concatenated to create a full taxonomic lineage. The list of terms is then tokenized using a vocabulary comprised of the top 40,000 taxonomic terms in the UniRef training dataset. At input, learned embeddings (dimension 768) for each term in the lineage are summed and projected by a learned linear projection to a single embedding of *d*_*model*_. This single embedding is then repeated across the length of the sequence and summed with the embeddings of all other tracks. The linear projection is zero-initialized at the start of this stage of training to preserve model behavior, enabling continuation of pretraining with no degradation in performance.

In pretraining we apply random corruption to the taxonomic lineages and train ESM3 to reconstruct the full lineage by predicting dropped terms with a shallow MLP head trained on the final layer’s representations. To corrupt the taxonomic lineage, we either drop all terms (probability 25%) or drop a set of the most specific terms of the lineage of size chosen uniformly at random from 1 to the length of the lineage (probability 25%). We also independently drop any taxonomic term with probability 10%. The output head outputs binary classification logits over the full set of 40,000 taxonomic lineage terms, and is trained to predict the missing terms via a binary-cross entropy loss.

###### A.1.10. ESM3 Inference

Since ESM3 is a bidirectional transformer capable of representing arbitrary factorizations of the joint probability in any order or combination of tracks, we have significant flexibility during inference. We can generate sequence, structure, or function conditioned on any combination of other tracks. We also have a choice of how much compute to apply during generation.

The usual inference strategy is to fix a prompt (which may be a combination of any of the tracks, either fully or partially specified) and choose a track for generation (which may have been partially specified). When predicting the tokens for the generation track, a number of strategies are possible. Here we discuss two notable strategies. Argmax decoding, which predicts all tokens in the generation track in a single forward pass of the model; this computation is *O*(*L*^2^) in the length of the protein and is extremely efficient. Iterative decoding, on the other hand, samples tokens one position at a time, conditioning subsequent predictions on those already sampled. The runtime for iterative decoding is *O*(*L*^3^) in the length of the protein.

Additionally, the number of decoding steps can be chosen at runtime. Argmax decoding is equivalent to decoding in one step, while iterative decoding is equivalent to decoding in *L* steps. It is possible to select any number of decoding steps between these two extremes to find an optimal tradeoff between computation and accuracy for a particular use case. See Appendix A.3.4 for a case study on structure prediction, in which the generation track is the structure tokens track.

When using iterative decoding, ESM3 further allows flexibility in choosing the next position to decode. We choose the position based off of the logits output of ESM3, and for the results of this paper utilize two strategies: entropy decoding, which chooses the position with the lowest entropy after softmax, or max probability decoding, which chooses the position with the maximum probability. To generate *k* tokens in one pass, we rank by either entropy or max logit and take the top *k* positions.

In the following algorithm, assume a single forward pass of ESM3 is a function *f* of a prompt *x*, and that we can access the logits of a specific token track through a subscript; e.g. sequence logits would be *f*_sequence_(*x*) ∈ ℝ^*L×*32^. Furthermore, denote *π* (·; *z*) as the probability distribution induced by the logits *z*, including an implicit softmax, and *T* ∈ ℝ^*L*^ a temperature schedule.

###### Algorithm 14

`generate` from track

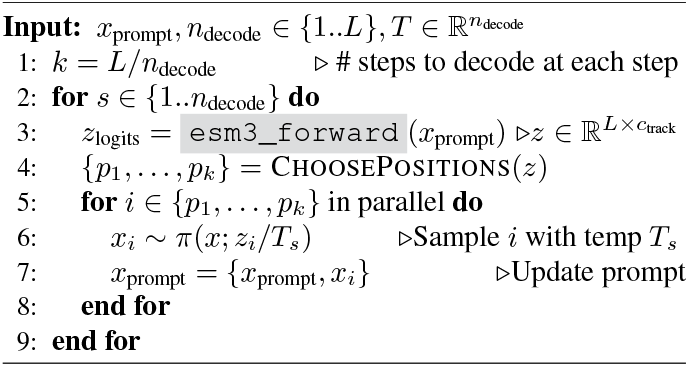

#### A.2. TRAINING ESM3

##### A.2.1. Pretraining Data

###### A.2.1.1. Sequence Databases

UniRef release 2023 02 is downloaded and parsed from the official UniRef website (78). MGnify90 version 2023 02 is downloaded and parsed from MGnify (38). All non-restricted studies available in JGI on July 31st, 2023 are downloaded and concatenated into the JGI dataset (79). OAS, which includes over a billion antibody sequences from 80 studies, is downloaded and clustered at 95% sequence identity (39).

###### A.2.1.2. Clustering

In all cases, data is clustered with mmseqs2 (80), with flags `--kmer-per-seq 100 --cluster-mode 2 --cov-mode 1 -c 0.8 --min-seq-id <seqid>`.

During training we re-weight the data distribution using clustering. We first sample a random cluster at the outer level, and then sample a random member of the cluster. To deduplicate cluster members, we perform a first step of clustering at a high level of sequence identity (e.g. 90%), taking one representative per cluster. We then cluster these deduplicated sequences at a lower level of sequence identity (e.g. 70%). We limit the number of samples within a cluster to 20.

###### A.2.1.3. Inverse Folding

As data augmention we train a 200M parameter inverse folding model and use it to create additional training examples.

The inverse folding model uses geometric attention for structure conditioning and an output projection head for the sequence logits, similar to ESM3. Unlike ESM3, the transformer stack alternates between blocks with geometric attention and standard attention. The model is trained on the sequence and structure pairs in PDB, AlphaFold-DB, and ESMAtlas, with the single training task of (and loss computed on) predicting sequence at the output given structure at the input. Model architecture and training methodology is otherwise substantially similar to ESM3.

This model is used to generate additional sequences corresponding to each structure in the training data for ESM3 (5 sequences per structure for ESMAtlas and AlphaFold-DB, 64 sequences per structure for the PDB). When training ESM3, with 50% probability the original sequence and structure pair is presented to the model as a training example. The other 50% of the time one of the inverse folded sequences is paired with the structure as the training example seen by ESM3.

###### A.2.1.4. Functional Labels

Functional labels are obtained from InterPro (36) and InterProScan (81), both version 95.0. All annotations for UniProtKB were downloaded from the InterPro website via the ‘protein2ipr.dat.gz’ file. InterProScan was applied to the entirety of MGnify90 with flags `--goterms –iprlookup --pathways --disable-precalc`. The resultant values are taken as ground truth functional labels for model training.

###### A.2.1.5. Structural Data

We use all PDB chains, clustered by unique PDB ID and entity ID within the PDB structure. We filter to all structures deposited before May 1, 2020, determined by X-ray crystallography, and better than 9A° resolution (40).

AlphaFoldDB is downloaded as the v4 version specified on their website (4). We notice that structures with high pLDDT are disproportionately alpha helices. Therefore, we ensure globularity by measuring the number of long range (*>*12 sequence distance) contacts in the chain. If this value is *<* 0.5L with an L length protein, we omit it from our training set. We also filter out all proteins *<* 0.7 pLDDT.

ESMAtlas is downloaded as version v0 and v2023 02. Similarly we use a *<* 0.7 pLDDT filter. We use a 0.7 pTM cutoff as well to enforce globularity. High pTM structures tends to be more compact.

Structural data also includes any functional labels that exist for the corresponding sequence.

###### A.2.1.6. Solvent Accessible Surface Area and Secondary Structure

For solvent accessibility surface area, we use the Shrake-Rupley rolling probe algorithm as implemented in biotite (82). This generates a set of real numbers, or a nan value when structural coordinates are not provided. Similarly, SS8 labels are generated using the mkdssp tool (83) and taken as ground truth labels. SS8 and SASA tracks were generated for the AlphaFoldDB and ESMAtlas datasets.

###### A.2.1.7. Purging of Validation Sequences

We keep track of validation set performance on a set of held out sequences from each training set: UniRef, MGnify, and JGI. In order to properly hold out a sufficiently diverse set of validation proteins, we first sample 25000 proteins from each set. Then we use mmseqs easy-search to filter out proteins from this set with a 70% sequence identity threshold. We choose the set of proteins from our training set to be the “query” set, and the set of validation proteins as our “target” set for mmseqs. We use the flags `--alignment-mode 3 -c 0.8 {cov-mode 0 --max-seqs 300 --max-accept 3 --start-sens 2 -s 7 --sens-steps 3`. This query is designed such that early stopping in mmseqs will not affect if we find a hit in the “query” training set.

Train purges are run to generate a list of blacklisted UniRef, MGnify, and JGI IDs, which are removed from the training set.

###### A.2.1.8. Token counts

The dataset counts in Table S3 are computed after limiting the large clusters to 20. The number of tokens are computed by multiplying the number of sequences with the average length of the dataset.

In order to compute the approximate number of sequences and tokens seen during training, we first compute the number of times the dataset is repeated at the cluster level. Given the number of repeats, we know the expected number of unique samples seen when sampling with replacement is 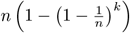 with a cluster of size *n* and *k* items selected. Computing this on the size of each cluster and number of dataset repeats results in the approximate number of tokens presented in Table S4. Our largest model is trained on all of this data, while our smaller models use a portion of it depending on the model’s token budget.

##### A.2.2. Pretraining Tasks

###### A.2.2.1. Noise Schedule

In the masked generative framework, corruption is applied to each input to the model. To enable generation, the amount of noise applied to an input is sampled from a distribution with probability mass on all values between 0 and 1.

We select various noise schedules for different tracks with several goals in mind. First, ESM3 should see all combinations of tracks as input and output, enabling it to generate and predict based on arbitrary inputs. Second, ESM3 should maintain a balance of strong representation learning and high quality generations. Third, the type of inputs provided should be representative of what users would like to prompt the model with.

In initial experimentation, we found that a fixed 15% noise schedule led to poor generation results, while a linear noise schedule where probability of each mask rate was constant led to good generation but poor representation learning results. We find a good trade-off between representation learning and generation by sampling the noise schedule from a mixture distribution. 80% of the time, the mask rate is sampled from a *β*(3, 9) distribution with mean mask rate 25%. 20% of the time, the mask rate is sampled from a uniform distribution, resulting in an average overall mask rate of 30%.

The noise schedules applied to each input are listed in Table S6. For the structure coordinate track, we also modify the masking to be applied as span dropping 50% of the time. This ensures that the model sees contiguous regions of masked and provided coordinates, which better mimics the types of inputs users may provide.

###### A.2.2.2. Track Dropout

Along with applying noise to each track, we want to ensure ESM3 is able to perform well when some tracks are not provided at all (e.g. to perform structure prediction when no structure is provided as input). We enable this by wholly dropping out some tracks with varying probabilities, listed in Table S6.

###### A.2.2.3. Loss Weights

Losses for each track are combined with weights listed in Table S6. Losses are also normalized on each rank by their relative number of loss contributing tokens, which helps to reduce gradient noise for tracks which are more sparsely present in pretraining data (e.g., residue annotations).

###### A.2.2.4. Structure Noise

We apply gaussian noise with standard deviation 0.1 to all coordinates the model takes as input.

###### A.2.2.5. Tertiary Coordination Sampling

An interesting use case of generative protein models involves conditioning on key structural information, such as an active site, and generating the sequence and structure of a protein that contains this information. It is possible to define an tertiary coordination task as 3 residues which are mutually in contact in structure space (*Cα* − *Cα* distance *<* 6A°), but are distant in sequence space (≥ 10 positions apart) (26). Training on this conditioning may enable the model to better perform the type of tertiary coordination required for active site sampling.

While this task will be sampled with some probability under the standard noise schedules, we also manually sample the task with 5% probability whenever a structure is available. If the task is sampled and a valid tertiary coordination triplet is found, the structure coordinates for that triplet are shown to the model. For each residue in the triplet, the adjacent residues are also independently shown with 50% probability, which leads to a total size of between 3 and 9 residues. All other structure coordinates are masked. Normal masking is applied to the other tracks.

###### A.2.2.5. Tertiary Interface Sampling

Predicting and generating binding interfaces is another important task for generative protein models. To help with this capability, we add computational data augmentation that simulates the binding interface task.

We define a tertiary interface as one involving a long range contact (*Cα* − *Cα* distance *<* 8A°, ≥ 24 sequence positions). When this task is sampled (5% probability whenever a structure is present) and a long range contact is found, then the chain is split into two chains, each containing one side of the contact interface. Suppose the contacting positions are given by the indices *i, j*. Then the first chain will contain residues between [randint(1, *i* − 3), randint(*i* + 3, *j* − 15)], while the second chain will contain residues between [randint(*i* + 15, *j* − 3), randint(*j* + 15, *L*)]. This ensures there is always a residue gap between the two pseudochains. A chainbreak token “—” is inserted to represent the residue gap.

**Table S3.**
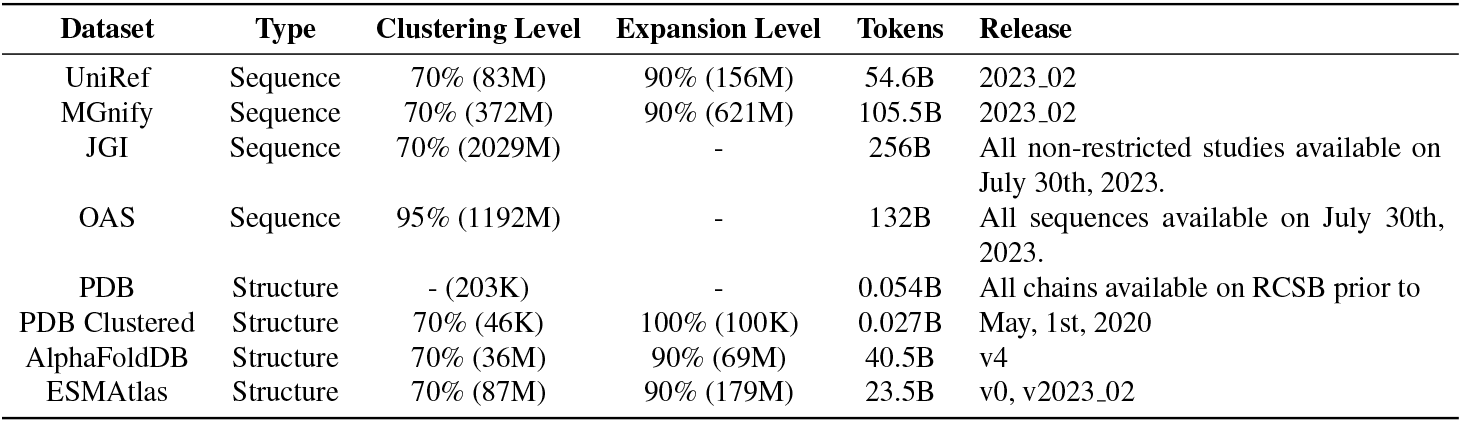
Pretraining dataset statistics. Includes number of tokens, release, and clustering level. Numbers are derived after dataset filtering.

**Table S4.** Pretraining unique token statistics. Broken down by token type and dataset type.

| Dataset Name | Unique Samples (M) | Unique Tokens (M) |
| --- | --- | --- |
| UniRef | 133 | 40,177 |
| MGnify | 406 | 65,780 |
| JGI | 2,039 | 265,070 |
| OAS | 203 | 22,363 |
| PDB | 0.2 | 55 |
| AFDB | 68 | 20,510 |
| ESMAAtlas | 168 | 38,674 |
| AFDB inverse folded | 111 | 33,300 |
| ESMAAtlas inverse folded | 251 | 57,730 |
| Sequence | 3,143 | 484,441 |
| Structure | 236 | 177,710 |
| Annotation | 539 | 105,957 |
| Total unique training tokens |  | 768,109 |

**Table S5.** Data augmentation and conditioning information applied to each dataset.

| Dataset | Inverse Folding | Function Labels | SASA | Secondary Structure |
| --- | --- | --- | --- | --- |
| UniRef | ✓ | ✓ | - | - |
| MGnify | ✓ | ✓ | - | - |
| JGI | ✗ | ✗ | - | - |
| OAS | ✗ | ✗ | - | - |
| PDB | ✗ | ✗ | ✗ | ✗ |
| AlphaFoldDB | ✓ | ✓ | ✓ | ✓ |
| ESMAAtlas | ✓ | ✓ | ✓ | ✓ |

**Table S6.** Noise schedules, dropout probabilities, and loss weights.

| Track | Noise Schedule | Dropout Prob | Loss Weight |
| --- | --- | --- | --- |
| Sequence | betalinear30 | 0 | 1.0 |
| Structure Tokens | cosine | 0.25 | 0.5 |
| Structure Coordinates | cubic | 0.5 | N/A |
| Secondary Structure (8-class) | square root | 0.9 | 0.01 |
| SASA | square root | 0.9 | 0.01 |
| Function Tokens | square root | 0.9 | 0.01 |
| Residue Annotations | square root | 0.9 | 0.01 |

**Figure S9.**
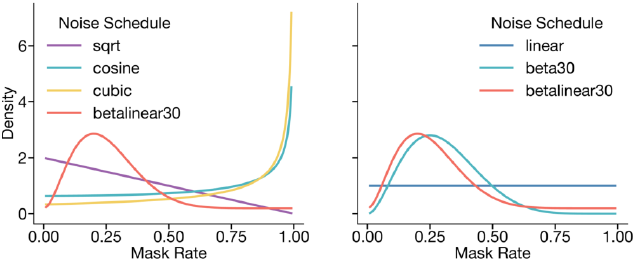
Visualization of noise schedules used. Left shows the probability density function of all noise schedules used. Right shows the betalinear30 distribution (which is drawn from *β*(3, 9) with 80% probability and a linear distribution with 20% probability) against a beta30 distribution (defined by *β*(3, 7)) and a linear distribution.

###### A.2.2.7. Residue Gap Augmentation

To encourage the model to learn to represent residue gaps using the chainbreak token, we introduce a task which randomly splits a single chain into multiple subchains.

First, a number of chains to sample is sampled from a geometric distribution with probability 0.9, up to a maximum of 9 possible chains. If the number of chains sampled is 1, no additional transformations are applied. A minimum separation of 10 residues between chains is defined. Sequence lengths of the chains along with gaps are sampled from a dirichlet distribution to maintain identically distributed sequence lengths for each chain. This transformation is applied to all samples.

###### A.2.2.8. Geometric Attention Masking

In the case that multiple chains are provided to the model from either the interface sampling or residue gap augmentation tasks, we mask the geometric attention layer to prevent the model from attending to cross-chain coordinates. This simulates tasks where the structure of individual chains is known, but the interface is unknown.

##### A.2.3. Training Details

###### A.2.3.1. Hyperparameters

We train all models using AdamW optimizer (84), with the following hyperparameters: *β*_1_ = 0.9, *β*_2_ = 0.95. We use a weight decay of 0.01 and gradient clipping of 1.0. We employ 5K to 20K warmup steps until reaching the maximum learning rate, and utilize a cosine decay scheduler to decay LR to 10% of the maximum learning rate by the end of training.

###### A.2.3.2. Infrastructure

Our training codebase uses Pytorch. We use Pytorch’s FSDP (85) implementation for data parallelism. We also use custom components from the TransformerEngine (86) library.

We have made several optimizations to increase the training speed of our models. For multi-head attention uses, we use the memory efficient implementation from the xformers library (87). We also save activations that are expensive to compute during training when necessary. We employ mixed precision training, utilizing FP8, BF16, and FP32 as needed based on accuracy requirements and kernel availability throughout our network.

###### A.2.3.3. Stability

Scaling ESM3 to 98 billion parameters with its novel architecture, multi-modal inputs, and low precision computation requirements poses significant training stability challenges. Our model is significantly deeper than its NLP counterparts, and literature has shown that deeper networks are harder to train due to attention collapse (88).

We observed training instability early in the architectural innovation phase, which we addressed through several changes. We apply layer normalization to the query and key vectors within the attention mechanism (89). We observe longer warm up helps (90). Another source of instability is the masking rate in pretraining tasks. We found that a very high masking rate is more likely to cause training divergences than a lower one, especially early in the training. Choosing a masking schedule biased towards lower mask rates improved both performance and training stability. Interestingly, the introduction of conditioning from other modalities also improves training stability, perhaps suggesting that stability is related to the degree of underspecification of a task.

An incorrectly set learning rate is another source of instability. To ensure the right balance between learning effectiveness and stability, we optimized the learning rate on smaller models and scaled it according to best practices as outlined in (91, 92). We find empirically that the initialization has a small effect on model stability, and the majority of stabilization can be gained from simply scaling the learning rate at the appropriate rate. By applying the rules in both width-*μ*P and depth-*μ*P, we can simply scale the learning rate inversely proportional to the square root of the number of parameters, and find this results in stable training.

Following these modifications, we successfully trained our 98-billion-parameter model without any issues related to training instability.

###### A.2.3.4. Staged training

We stage training to alter dataset composition, train on longer contexts that would be too expensive for the entire pretraining, or introduce features such as the taxonomy track (A.1.9.2).

#### A.3. MODEL EVALUATIONS

ESM3 is both a generative model and a representation learning model that can be adapted for predictive tasks. In this section, we present benchmarking results for both capabilities.

##### A.3.1. Models

ESM3 models are trained at three scales—1.4B, 7B, and 98B parameters—on approximately 75B, 560B, and 1.8T training tokens, respectively.

The ESM3 1.4B model, trained on 75B tokens and noted for its small size and speed, allows rapid iteration both during training and at inference. Optimal model size and number of training tokens are studied by extrapolating from a series of smaller runs, given a training compute budget, model architecture, and dataset characteristics (22, 24). After determining compute optimality for training, a variety of factors such as release frequency, amount of inference, ease of use, and usage patterns are also taken into account to determine the ideal number of tokens on which to train the model. To enable efficient inference for the benefit of the research community, we have trained two additional versions of ESM3 1.4B, named 1.4B Overtrained and 1.4B Open, which are trained on 300B tokens, far beyond their compute optimality for training.

##### A.3.2. Evaluation Data

In the following benchmarks for this section, unless otherwise noted, models are evaluated on a test set of 902 proteins whose structures are temporarily held out from the ESM3 training set. The proteins were sourced from the Continuous Automated Model EvaluatiOn (CAMEO) targets released from May 1, 2020 through Aug 1, 2023 (93). A summary of the datasets used in each model evaluation is provided in Table S7.

For contact and structure prediction evaluations, we also evaluate on the CASP14 (71 proteins) and CASP15 (70 proteins) structure prediction benchmarks (94, 95). The CASP14 and CASP15 sets are obtained directly from the organizers and include both free modeling and template-based modeling single domain targets. Both CASP sets are temporally held out from the PDB training set, which has a cutoff date of May 1, 2020.

##### A.3.3. Representation Learning

The contact prediction model is a multilayer perceptron (MLP) head that operates independently over the representations of each amino acid pair, outputting the probability of contact between them. We use LoRA (96) for finetuning, which is a common alternative to full weight finetuning that uses much less memory while attaining strong performance. LoRA is applied to the base model for finetuning, and the MLP along with the LoRA weights are trained end-to-end using the cross-entropy loss with respect to the ground truth contact prediction map. For the ground truth, all residues at least 6 positions apart in the sequence and within an 8A° C*α*-C*α* distance are labeled as a contact. All models are trained with LoRA rank 4, batch size 64 and a learning rate of 1e-3 for 10k steps on a mix of sequence and structure data from PDB, AlphaFold-DB, ESMAtlas, and OAS Predicted Structures. Data are sampled in a ratio of 1:3:3:0.03 from these datasets.

**Table S7.** Datasets for model evaluations.

| Reference | Task | Dataset | Holdout Method |
| --- | --- | --- | --- |
| Fig. 1D, Table S10, Fig. S11 | Conditional and unconditional NLL | CAMEO | Temporal |
| Fig. 2A | Generative prompt following | CAMEO | Temporal |
| Fig. S3, Fig. S4 | Structure tokenizer reconstruction | CAMEO, CASP14, CASP15 | Temporal |
| Fig. S6 | pTM, pLDDT calibration | CAMEO | Temporal |
| Table S8 | Contact prediction | CAMEO, CASP14, CASP15 | Temporal |
| Table S9, Fig. S10 | Structure prediction | CAMEO, CASP14, CASP15 | Temporal |
| Fig. S8 | Function prediction | UniRef | 70% sequence holdout |

Table S8 shows the performance on each structural test set through the metric of precision at L (P@L), which evaluates the precision of the top-L most confident predictions, where L is the length of the protein. The smallest ESM3 model, with 1.4B parameters, achieves a P@L of 0.76 ± 0.02 on the CAMEO test set, which is higher than the 3B parameter ESM2 model (0.75 ± 0.02). Furthermore, it trains on an order of magnitude less compute during pretraining (6.72 × 10^20^ FLOPs vs. 1.8 × 10^22^ FLOPs), demonstrating the benefits of multimodal pretraining.

##### A.3.4. Structure Prediction

ESM3 can directly predict protein structures without additional finetuning by first predicting structure tokens, then decoding these tokens into coordinates. When predicting structure tokens, we follow the strategy outlined in Appendix A.1.10 and test both single-pass argmax decoding and full iterative decoding.

Iterative decoding and single-pass decoding have equivalent performance for structure prediction. We hypothesize that this is because ESM3 has a latent representation of structure, so whether structure tokens are decoded in a single pass or iteratively does not meaningfully change the sampling trajectory. Furthermore, ESM3 was trained with dropout of the structure tokens track, which mimics single-pass decoding (see Appendix A.2.2).

Structure prediction scaling curves as a function of training compute, are provided in Fig. S10.

##### A.3.5. Conditional Likelihood

The conditional likelihood of an output given a prompt serves as a proxy for the generative capabilities of a model. Fig. S11 and Table S10 evaluate the performance of ESM3 as a conditional generative model, using its negative log-likelihood (NLL) on the test set. For each track—sequence, structure, function, SASA, and secondary structure—NLL is evaluated both unconditionally and conditioned on each of the other tracks.

**Figure S10.**
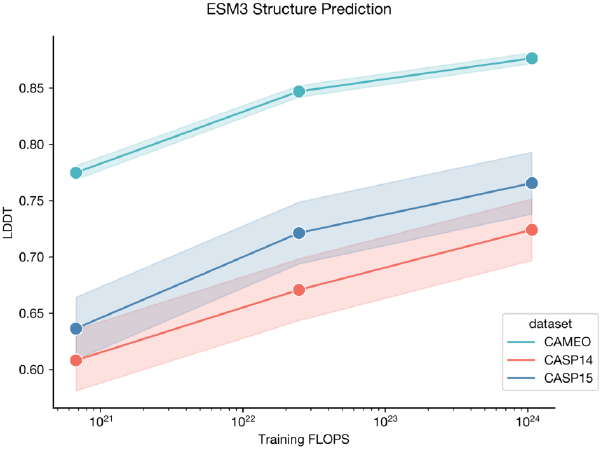
Scaling curves for structure prediction. Error bars are ±1 standard error.

Unlike, for example, an autoregressive model, ESM3 is a generative model over masking patterns, so is trained to predict tokens given any masking pattern. The NLL of a sample under ESM3 is given by 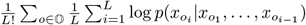, where *O* is the set of all decoding orders with normalization constant 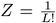. This computation is intractable (as the set of all decoding orders is exponential in length of a protein), but can be approximated by sampling a single decoding order *o* for each *x* in our dataset. At each step, teacher forcing is used to replace the masked token with the ground truth token and report the mean NLL over the output tokens.

We observe a number of interesting trends in these results. As expected, the unconditional NLL (Fig. S11, black lines) is always higher than conditional, and conditioning on full 3D structure reduces the loss on secondary structure prediction to nearly zero (1.4B: 0.24, 7B: 0.19, 98B: 0.16). Conditioning on sequence results in a lower structure prediction loss than conditioning on secondary structure (98B; sequence: 3.13, secondary structure: 3.37). We observe a clear log-linear relationship between pretraining FLOPs and NLL for the sequence track, under each form of conditioning.

**Table S8.** Precision@L results. Measured on CASP14, CASP15 and CAMEO for the ESM3 model family. Intervals represent bootstrapped 95% confidence intervals.

| Model | CASP14 | CASP15 | CAMEO |
| --- | --- | --- | --- |
| ESM2 3B | 0.57 (0.49 - 0.64) | 0.57 (0.48 - 0.65) | 0.75 (0.73 - 0.77) |
| ESM3 1.4B | 0.56 (0.48 - 0.64) | 0.59 (0.50 - 0.66) | 0.76 (0.74 - 0.78) |
| ESM3 7B | 0.62 (0.54 - 0.70) | 0.64 (0.56 - 0.73) | 0.82 (0.80 - 0.84) |
| ESM3 98B | 0.66 (0.57 - 0.74) | 0.66 (0.57 - 0.75) | 0.85 (0.83 - 0.86) |

**Table S9.** Protein structure prediction results. We benchmark ESMFold, ESM3 models, and Alphafold2. Left side: ESM3 iterative inference of structure tokens conditioned on sequence. Because iterative inference is *O*(*L*^3^) in length of a protein sequence, it is comparable to the runtime complexity of ESMFold and AF2. Right panel: Single pass argmax prediction of structure tokens given sequence. Iterative decoding does not significantly improve structure prediction performance, and *O*(*L*^2^) runtime complexity is sufficient for structure prediction. Both the Open and Overtrained models are trained up to 200k steps. The 1.4B model is used for scaling comparisons, and is trained to 50k steps.

| Model | Iterative / $O(L^3)$ | | | Single pass / $O(L^2)$ | | |
| --- | --- | --- | --- | --- | --- | --- |
|  | CAMEO | CASP14 | CASP15 | CAMEO | CASP14 | CASP15 |
| 1.4B Open | 0.801 | 0.624 | 0.691 | 0.806 | 0.639 | 0.695 |
| 1.4B Overtrained | 0.816 | 0.623 | 0.704 | 0.821 | 0.637 | 0.707 |
| 1.4B | 0.771 | 0.596 | 0.654 | 0.777 | 0.613 | 0.654 |
| 7B | 0.845 | 0.661 | 0.735 | 0.848 | 0.670 | 0.738 |
| 98B | 0.880 | 0.724 | 0.787 | 0.879 | 0.724 | 0.782 |
| ESMFold | 0.861 | 0.722 | 0.745 |  |  |  |
| AlphaFold2 | 0.903 | 0.857 | 0.833 |  |  |  |

To better understand the effect of function conditioning, we plot the sequence token NLL gap under function conditioning across the range of mask rates (Fig. S12). We observe that the NLL gap is larger at higher mask rates (e.g., 0.20 at a 90% mask rate) but is negligible at lower mask rates. These results suggest that at low mask rates, there is sufficient information in the sequence for the model to infer the remaining tokens without needing additional information from the function track. We further analyze the function-conditioned NLL gap by InterPro tags, finding that tags for more common domains show larger NLL gaps. For instance, IPR015426 has a mean NLL gap of 0.59, comparable to the gap between sequence and structure conditioned sequence NLL in the 7B model (0.61).

##### A.3.6. Unconditional Generation

To assess the model’s unconditional generation capability, we sampled 16 proteins for each length *L* sampled uniformly every 4 values between 64 and 1024 (i.e. {64, 68, 72, …, 1024}) using ESM3 98B with a constant temperature of 0.7. Structures for each sequence were predicted using ESM3 7B, and the resulting structures have high confidence under ESM3, even for sequences over 1000 amino acids in length, as shown in Fig. S14A. The distribution of pTM and pLDDT are shown in Fig. S14B. ESM3 generates more high-quality structures than ESM2, which was trained using a simple MLM objective over sequence only with a fixed mask rate.

We use pTM for evaluating structure predictions from ESM3 in addition to pLDDT. This is because pLDDT can be miscalibrated for generated structures and can overestimate the confidence of a prediction. pLDDT is biased towards local structural confidence, which can result in pathologies such as very long alpha helices with high pLDDT at all positions. pTM is a more global measure of structural confidence, and is more robust to these pathologies. Fig. S13 shows that pTM and pLDDT correlation drops for generated sequences (Pearson r: natural = 0.85, generation = 0.79), and a clear pattern of high pLDDT (*>* 0.8) but low pTM (*<* 0.6) emerges.

To assess whether ESM3 is biased towards particular secondary structures, we use DSSP to predict the three-class secondary structure of the high-confidence (pTM *>* 0.8, mean pLDDT *>* 0.8) generations and measure the percentage of residues that form alpha helices and beta sheets. When compared to a background distribution computed over the PDB, we find that ESM3 closely matches the secondary structure distribution of known proteins (Fig. S14D), unlike other methods which preferentially generate helical structures (15, 26, 28). Finally, to confirm that the structures predicted with high confidence by ESM3 are designable, we inverse folded and re-folded each using ESM3 7B. The majority of generations successfully re-folded with TM-score of greater than 0.8 to the hallucinated structures, demonstrating that ESM3 has high self-consistency for its own high-confidence designs (Fig. S14C).

**Table S10.** Negative log-likelihood of each track conditioned on other tracks. Each row is a model size, generating a particular modality. Each column is the conditioning. The diagonal, highlighted with italics, are the unconditional NLL of each track. We observe that indeed adding conditioning improves NLL in all cases.

|  | Model | Conditioning |  |  |  |  |
| --- | --- | --- | --- | --- | --- | --- |
|  |  | Sequence | Structure | Function | SASA | Secondary Structure |
| Sequence | 1.4B | *2.31* | 1.71 | 2.28 | 1.81 | 2.02 |
|  | 7B | 2.04 | 1.43 | 2.00 | 1.47 | 1.74 |
|  | 98B | 1.84 | 1.21 | 1.76 | 1.21 | 1.50 |
| Structure | 1.4B | 4.09 | *4.98* | 4.93 | 4.39 | 4.42 |
|  | 7B | 3.42 | 4.20 | 4.18 | 3.62 | 3.71 |
|  | 98B | 3.13 | 3.85 | 3.80 | 3.24 | 3.37 |
| Function | 1.4B | 1.81 | 1.98 | *4.52* | 2.29 | 2.24 |
|  | 7B | 1.22 | 1.47 | 3.75 | 1.67 | 1.70 |
|  | 98B | 0.93 | 1.20 | 3.63 | 1.41 | 1.40 |
| SASA | 1.4B | 1.78 | 1.81 | 2.42 | *2.48* | 2.10 |
|  | 7B | 1.57 | 1.66 | 2.26 | 2.31 | 1.92 |
|  | 98B | 1.46 | 1.56 | 2.15 | 2.23 | 1.82 |
| Secondary Structure | 1.4B | 0.42 | 0.24 | 0.70 | 0.50 | *0.83* |
|  | 7B | 0.31 | 0.19 | 0.57 | 0.31 | 0.60 |
|  | 98B | 0.26 | 0.16 | 0.50 | 0.25 | 0.54 |

**Figure S11.**
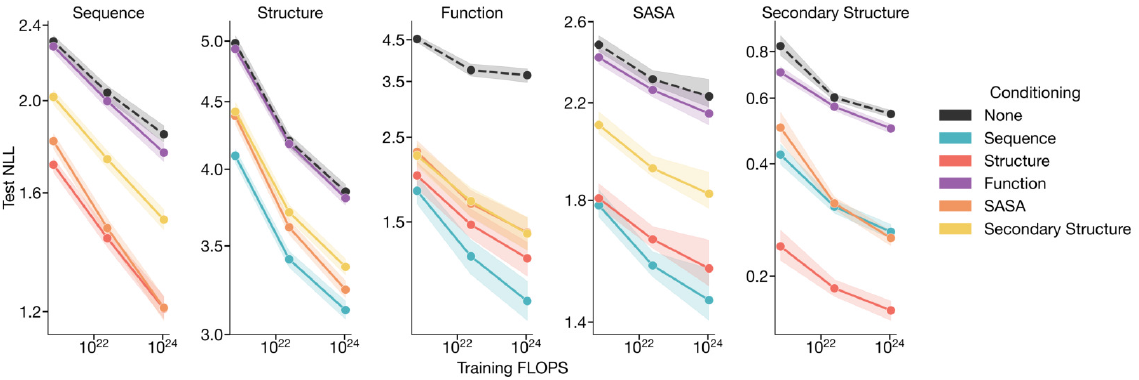
Conditional and unconditional scaling behavior for each track. Loss is shown on the test set (Appendix A.3.2). Shaded regions denote 95% confidence intervals.

**Figure S12.**
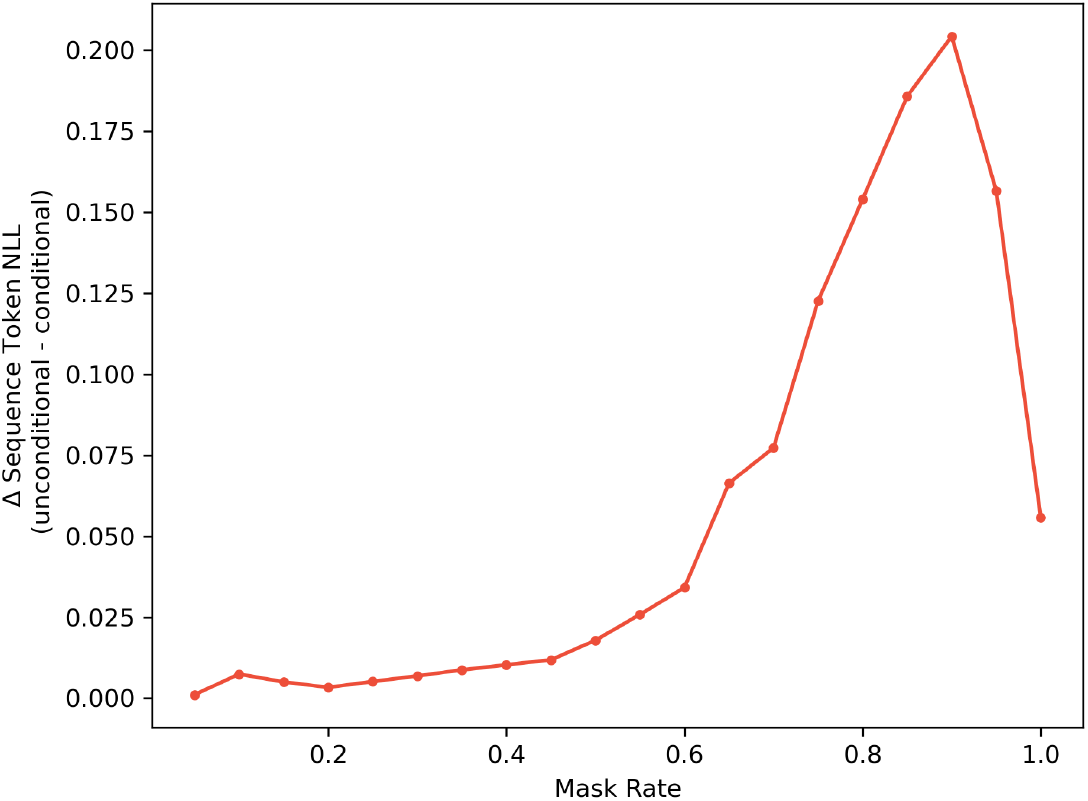
Function-conditioned sequence negative log-likelihood over mask rates. Loss is computed on the test set (Appendix A.3.2) using the ESM3 base 7B model. The sequence NLL loss is computed only in sequence positions constrained by function conditioning. The gap in sequence token NLL with and without function conditioning is plotted over a range of sequence token mask rates. Function conditioning drives down the loss at higher mask rates.

**Figure S13.**
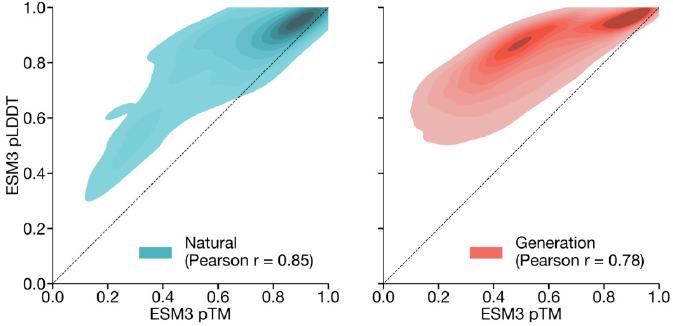
Distribution of pTM and pLDDT. Measured on natural (left) and generated (right) sequences under ESM3 7B structure prediction. Generated sequences show a clearly lower correlation (Pearson r 0.79 vs. 0.85) as well as a mode of sequences with high pLDDT but low pTM. Natural sequences are from the test set (Appendix A.3.2), generations are unconditional generations from ESM3 98B.

To visualize the distribution of unconditional generations, we generate a larger dataset by sampling 100 protein lengths randomly from the PDB and generating 1,024 sequences for each, again using ESM3 98B with a constant temperature of 0.7. We compute sequence embeddings by extracting the final layer outputs produced by running ESM3 7B with sequence inputs only. Protein-level embeddings are computed by averaging over all positions in the sequence to produce a 2560-dim embedding. We then project these embeddings into two dimensions using a UMAP projection (97) fit on a background distribution of 50,000 randomly sampled sequences from UniProt with minimum distance 0.1 and number of neighbors 25. Examples are selected by computing structural clusters with Foldseek-cluster (using default parameters) and sampling the example with highest ESM3 pTM from each cluster. A subset of these cluster representatives are shown in Fig. 1E. Sequence similarity to the training set was computed using mmseqs2 (80) with the following parameters: `--cov-mode 2 -c 0.8 -s 6.0`. Proteins generated unconditionally are similar—but not identical—to proteins found in the training set (Fig. S16) and have high coverage of the training set (Fig. 1E), demonstrating that the model has properly fit the training distribution and does not exhibit mode collapse. We observe a cluster of generations with very high sequence identity to the training set; these correspond to antibody sequences, with the framework regions accounting for the high sequence identity.

To explore alternative ways of generating proteins, we assess the quality of proteins generated by a chain-of-thought (CoT) procedure in which ESM3 7B generates the secondary structure (SS8 tokens), then the 3-D backbone coordinates (structure tokens), followed by the amino acid sequence (sequence tokens) (Fig. S15). We compare the quality of amino acid sequences generated from this CoT procedure with the above method of unconditionally directly generating amino acid sequences. We find that the CoT procedure generates sequences that have higher confidence ESM3-predicted structures than the directly-generated sequences as measured by pTM and mean pLDDT (Fig. S15A). Compared to high-confidence (pTM *>* 0.8, mean pLDDT *>* 0.8) directly-generated sequences, the high-confidence subset of CoT-generated sequences are also more designable: the CoT-generated sequences have predicted structures whose inverse folded, then re-refolded structures have higher TM-score to the originally predicted structure (Fig. S15C). The CoT-generated sequences show a small bias towards higher alpha and beta proportion compared to those generated directly (Fig. S15D).

##### A.3.7. Prompt Construction

Prompts for ESM3 specify inputs for each track in every position along the sequence length of the protein, which is determined by the positions of BOS and EOS tokens. ESM3 can scaffold motifs or hotspots with unknown relative positions by shuffling their positions in the prompt and generating multiple samples.

A summary of prompt lengths and constraint position variation for each design task is provided in S14.

##### A.3.8. Prompt-following Evaluations

To evaluate ESM’s ability to follow prompts, we use a set of held-out proteins as described in Appendix A.3.2. The test set is further filtered to remove proteins with length greater than 1024, which removes 7 proteins from the test set. To construct prompts for the structure coordinate, secondary structure, and SASA tracks, we sample a random span of length 15% of the original protein length. The model is then shown the corresponding track for the randomly sampled span, and is tasked with generating the sequence for the entire protein. For example, for the structure track, for a protein of length 100, we may sample a random span of 15 residues from residue 20-35. The model would then have to generate a protein sequence of length 100 conditioned on structure coordinate conditioning from residues 20-35 derived from the original test protein. This same procedure is applied for the secondary structure and SASA tracks. For the function track, we form the prompt by tokenizing the keywords from the InterProScan annotations associated with each sequence. The ESM3 7B model is used for all generations with a temperature of 0.7 and *L* decoding steps (where *L* is the length of the sequence). The model generates 64 sequences per prompt, which we use to compute pass@64.

To evaluate the generations, we use ESMFold to fold the sequences generated by ESM3. For the structure coordinate, secondary structure, and SASA tracks, alignment metrics (backbone cRMSD, 3-class secondary structure accuracy, and SASA Spearman *ρ*) can be calculated using the relevant span in the ESMFold-predicted structure and the original template protein. Continuing the previous example for the structure track, we would compute the RMSD between residues 20-35 in the ESMFold structure predicted of the ESM3-generated sequence and residues 20-35 of the original test protein. For the function annotation track, we run InterProScan (36) on each generated sequence and extract function keywords from the emitted annotations. We report function keyword recovery at the protein level, computing the proportion of all function keywords in the prompt which appear anywhere in the function keywords from the Inter-ProScan annotations of the generation.

**Figure S14.**
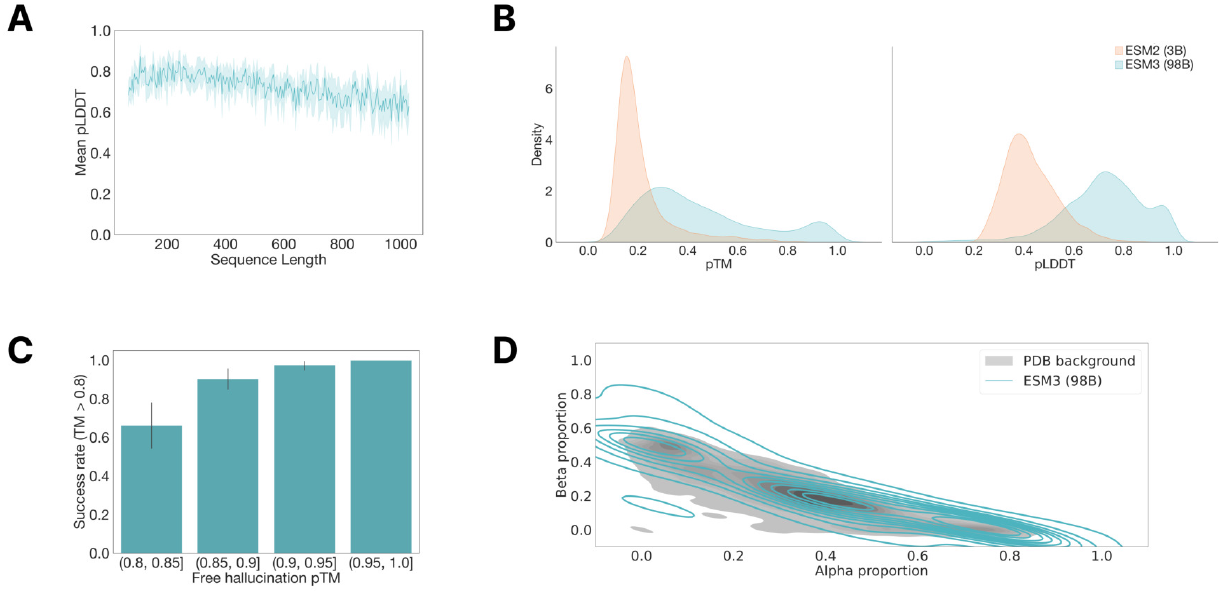
Unconditional generation of high-quality and diverse proteins using ESM3. (A) Distribution of ESM3 pLDDT (mean and 95% confidence intervals) for each sequence length in the unconditional generation dataset. (B) pLDDT and pTM of unconditional generations from ESM3 compared to sequences designed using the 3B-parameter ESM2 model. (C) Round-trip success rate of high-confidence generations using ESM3. Predicted structures were inverse folded to predict a new sequence and then re-folded to produce a new structure. Success was measured by a TM-score of greater than 0.8 between the original and refolded designs. (D) Secondary structure composition of unconditional generations relative to the distribution of proteins in the PDB, which is shown in gray.

**Figure S15.**
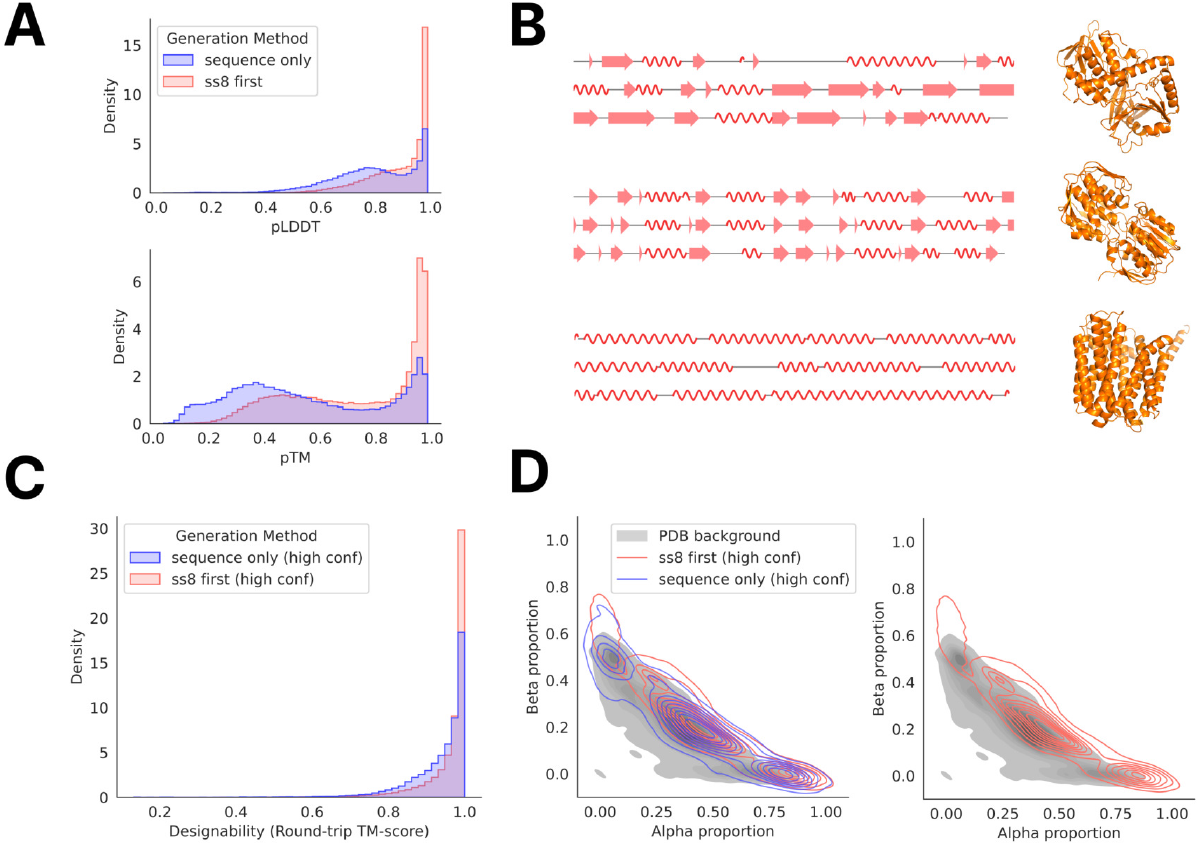
Generation of sequences using chain of thought. SS8 tokens are generated first, followed by structure tokens, then amino acid sequence with the ESM3 7B model. (A) Distribution of mean pLDDT and pTM of sequences generated by chain of thought (“ss8 first”) compared to directly generating the sequence (“sequence only”). (B) Sample generations of SS8 tokens and the predicted structure of its corresponding CoT sequence. (C) TM-score between predicted structures of high-confidence (pTM *>* 0.8, mean pLDDT *>* 0.8) generated sequences and their corresponding inverse folded, then re-folded structures. (D) Comparison

##### A.3.9. Steerable Design

To test the ability of ESM3 to generalize beyond its training distribution under prompting, we evaluate two prompting scenarios. First, we identify proteins which were deposited in the PDB after our training cutoff (December 2020) and choose eight with TM *<* 0.7 to any structure in our training dataset (PDB IDs: 2JVN chain A, 2KAF chain A, 2L8K chain A, 2MJM chain A, 7ZUO chain A, 8EXF chain B). Using DSSP, we compute the residue-level SS8 and SASA for each of these proteins to prompt ESM3, masking all other tracks. We show in Fig. S16A that the generated proteins are diverse, globular, and closely follow the SS8 and SASA prompts while having no close sequence or structure neighbors in the training set. Interestingly, the original protein sequences are not folded with high confidence or accuracy by ESMFold (mean pTM 0.44, mean TM-score to reference 0.33), suggesting that these are challenging proteins to fold. The ESM3-generated sequences have a similar confidence (mean pTM 0.45) but much higher accuracy (mean TM-score 0.64).

Second, we classify the residue-level secondary structure for a set of eight symmetric protein backbones using DSSP. These proteins were previously designed using ESMFold (5, 98) and have varying secondary structure (alpha and beta) and varying symmetries (5-fold and 8-fold). Again, ESM3 is able to design these proteins successfully with high confidence (pTM *>* 0.8, pLDDT *>* 0.8) and low sequence similarity to the training set Fig. S16B. The structural similarity is moderate for these designs due to the high structural conservation of the protomer units in each design. All designs are generated using a constant temperature of 0.7 with L/2 decoding steps, where L is the protein length. We sample 256 sequences for each prompt and filter generations by pTM (*>* 0.8), pLDDT (*>* 0.8), and accuracy in satisfying the SS8 prompts (SS8 accuracy *>* 0.8). Final examples were selected from these filtered designs by visual inspection. Sequence similarity to the training set was computed using the same procedure as the unconditional generations, and structure similarity was computed using Foldseek (41) in TM-score mode (`alignment-type 1`) with sensitivity `-s 7.5`.

##### A.3.10. Composing Prompts

ESM3 is able to compose multimodal prompts across its input tracks—sequence, structure, SS8, SASA, and function keywords—to generate proteins with novel characteristics. To demonstrate this, we augment the standard functional motif scaffolding task (i.e., partial structure and sequence prompts) with additional conditioning to specify the type of scaffold for ESM3 to design. The functional sites comprise a combination of ligand binding sites coordinated by residues remote in sequence and those defined by short local motifs. For each motif, the coordinates and amino acid identities of all residues from the reference PDB structures are input to the model, with random shuffling and augmentation of the gaps between each active site. See Appendix A.4.5 for a description of this augmentation procedure and the specifications of the ligand-binding sites chosen. In addition to these sites, we also create a set of 12 partial sequence and structure prompts derived from conserved functional motifs (Table S11). These motifs are defined using a combination of the benchmark dataset in Watson et al. (26) and conserved sequence patterns from the Prosite database (99).

The scaffold conditioning is defined using either SS8 tokens (to specify secondary structure composition) or function keywords defined by InterPro accession numbers (to specify a particular fold). For each combination of functional site and scaffold prompt, we sample between 256 and 2048 times to generate proteins with diverse and novel characteristics. All designs were generated with the 7B-parameter model, a constant temperature of 0.7, and *L/*2 decoding steps for a protein of length *L*.

###### Secondary structure prompting

We generated proteins under four main classes of secondary structure composition: mostly alpha helices, mostly beta sheets, and mixed alpha-beta proteins (split into alpha/beta, alpha/beta/alpha, and beta/alpha/beta topologies). For each generation, we prompt the model with a random set of SS8 spans up to a total length *L*, with mask tokens in between. For example, an all-alpha SS8 prompt for a protein of length *L*=20 might look like `__HHHH__HHHHH___HH` and a beta-alpha-beta prompt might look like `__EEE__HHHHH___EE_`, where H is a residue within an alpha helix and E is a residue in a beta strand. We then combine this with the augmented partial structure and sequence tracks given by a functional site motif. To increase the diversity of the scaffolds and maximize the probability of generating physically realizable prompt combinations, we generate between 256 and 1024 designs for each combination of SS8 and functional site motif. For each generation, we uniformly sample a random length *L* between 150 and 400. Then, we produce a set of secondary structure spans with length 5-20 residues, each separated by a gap of 3-10 residues, such that the total length adds up to *L*. Finally, to avoid incompatibility between the partial structure and secondary structure constraints, we also mask the SS8 tokens at positions where structure is specified by the functional site prompt. Secondary structure–prompted designs were assessed by running DSSP on the designed sequence and measuring the fraction of prompted residues which were assigned the correct secondary structure. Success was determined by a pTM *>* 0.8, all-atom cRMSD *<* 1.5 for the functional site, and SS8 accuracy *>* 0.8.

**Figure S16.**
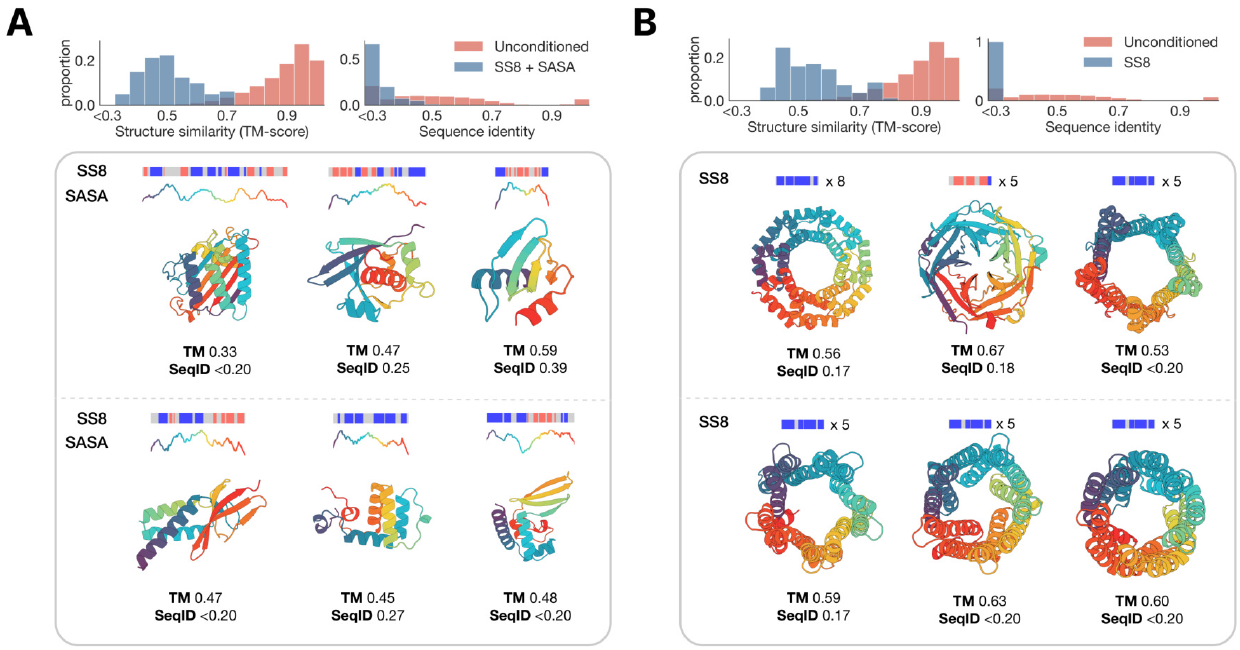
Prompting ESM3 to generalize beyond its training distribution. (A) Proteins designed using SS8 and SASA prompts derived from recent structures in the PDB with low structural similarity to the training set. Prompts along the protein length are visualized above each generation; secondary structure is shown using three-class (alpha = blue, beta = orange, coil = gray) and SASA is shown as a line plot colored by residue index to match the cartoon below. (B) Symmetric proteins designed using SS8 prompting. Histograms show the similarity to the nearest training set protein by structure (TM-score) and sequence (sequence identity) compared to unconditional generation.

**Table S11.**
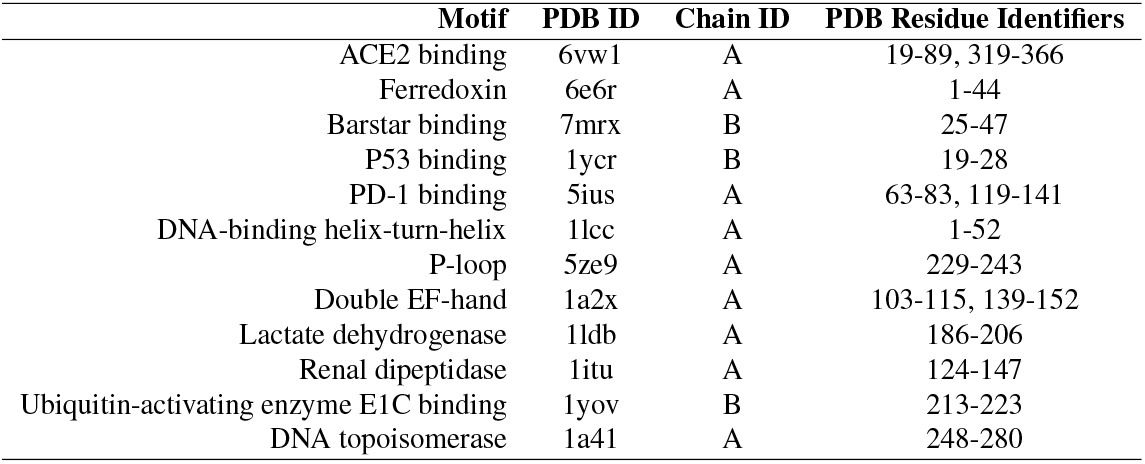
Functional motif definitions for conserved regions.

###### Keyword prompting

To prompt the model to generate proteins with a specific fold, we extracted the set of InterPro tags associated with a set of proteins from the test set for which ESM3 achieved keyword recovery of greater than 80% (Fig. 2A). These tags were then converted into keywords and used to prompt the model in combination with the partial sequence and structure constraints. The list of prompts and function tags is given in Table S12. Keyword-prompted designs were assessed using a self-consistency evaluation, i.e. whether the model successfully predicts any of the prompted InterPro accessions for the designed sequence. Success was determined by a pTM *>* 0.8, all-atom cRMSD *<* 2.0, and number of InterPro accessions recovered *>* 0.

We assess novelty of each motif-scaffold combinations by measuring the TM-score between the generated scaffold and the chain from which the motif is derived (Table S13). This confirms that the model is not retrieving the original motif scaffold, particularly for secondary structure–prompted scaffolds where we do not provide any explicit instructions to produce diverse designs. For the motifs derived from ligand binding residues (magnesium, serotonin, calcium, zinc, protease inhibitor 017, and Mcl-1 inhibitor YLT), we additionally use Foldseek to search the PDB for any other proteins which share that motif (as defined by BioLiP (100)), as a more stringent evaluation of novelty. For all but zinc-binding and magnesium-binding motifs, Foldseek finds no significant hits at an E-value threshold of 1.0. The hits discovered for zinc and magnesium have only modest TM-score (0.76 and 0.64), demonstrating that the model still finds novel scaffolding solutions for these ligands. To assess whether the generated scaffolds are likely to be designable, we measure a self-consistency TM-score (scTM) under orthogonal computational models by inverse-folding the designed structure with ESM-IF1 (42) (using a temperature of 0.5) and re-folding with ESMFold (5). We report the best scTM over 8 inverse folding designs in Table S13.

##### A.3.11. Multimodal Editing Examples

First, we describe the procedure for generating the protein compression example shown in Fig. 2D. A series of prompts of length 150 were constructed. The sequence and structure of the catalytic triad of trypsin (PDB 1Y3V) (H57, D102, S195) were placed in the prompt using the following procedure: three random residue numbers between 20 and 130 were sampled such that the minimum pairwise difference in position between each of the residues was no less than 20. Then, H57 from the template trypsin was placed at the lowest sampled number, D102 at the second lowest, and S195 at the largest number, thus respecting the left-to-right ordering of the catalytic triad in the template trypsin. 128 prompts were generated by this procedure. Each of these prompts was combined with a function key-word prompt derived from the template protein, specifically InterPro (36) tags IPR001254 (serine proteases, trypsin domain) and IPR009003 (peptidase S1, PA clan), to arrive at a final set of 128 prompts. The base ESM 7B model was then prompted to generate the sequence of the remaining 147 residues of the protein conditioned on the randomly placed catalytic triad sequence and structure coordinates and function keywords. *L* = 150 decoding steps were used with a temperature of 0.7, with 32 generations per prompt. Generations were then filtered by all-atom RMSD at the prompt positions, ESM3 pTM, and InterPro Scan keyword outputs, with the generation shown in Fig. 2D selected finally by visual inspection.

Generation quality was measured using ESMFold (5) pTM of the generated sequence, in addition to self-consistency. For self-consistency, we inverse fold the ESM3-predicted structure of the generation with ESM-IF1 (42) 8 times and re-fold with ESMFold, reporting the mean and std of the TM-scores between the 8 ESMFold-predicted structures and the ESM3-predicted structure. To perform a blast search of the sequence, we use a standard Protein Blast search (54). We set the max target sequences parameter to 5000 and sort results by sequence length and sequence identity, selecting the first sequence that is a serine protease. This yields the reference WP 260327207 which is 164 residues long and shares 33% sequence identity with the generation.

We showcase two further examples of protein editing. First, ESM3 is prompted to bury an exposed helix in a protein with an alternating alpha-beta sandwich fold. The prompt is constructed as follows: the prompt is of the same length as the template protein (PDB 1LBS). We identify a buried helix (mean SASA 0.32 A° 2) between residues 106-116 of the template protein. Structure coordinates from this region are placed in the prompt at the same residue indices, to prompt ESM3 to generate the same helix. This is composed with a SASA prompt of 40.0 for each of the 11 helix residues, prompting ESM3 to place this helix on the surface of the protein. Finally, we prompt with the secondary structure of 5 central beta strands surrounding the buried helix, residues 33-36, 62-65, 99-103, 125-130, and 179-182. ESM3 7B is then used to generate 512 protein sequences conditioned on this prompt using 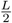 decoding steps and a temperature of 0.7. Designs are filtered by ESM3 pTM and adherence to the SASA prompt. The final generation is chosen by visual inspection. The generation is evaluated as described above (ESMFold pTM 0.71, scTM mean 0.82, std 0.045). Examining the generation, ESM3 is able to satisfy the input constraints: the generated protein maintains the structure of the helix (backbone cRMSD 0.18 A°) and the alternating alpha-beta fold (both the generation and the template have 7 strands alternating with helices), while exposing the helix motif to the surface (mean SASA 28.35 A° 2). Furthermore, the generation is structurally distinct: a Foldseek search (41) of AlphaFold-DB, ESMAtlas, and PDB in TM-align mode reveals no hit with TM-score greater than 0.76.

**Table S12.** InterPro tags extracted from CAMEO test set proteins for prompting with fold specification.

| Scaffold | Reference | InterPro tags | Total Prompt Length |
| --- | --- | --- | --- |
| Beta propeller | 8siuA | IPR001680 (1-350)<br>IPR036322 (1-350)<br>IPR015943 (1-350) | 353 |
| TIM barrel | 7rpnA | IPR000652 (0-248)<br>IPR020861 (164-175)<br>IPR035990 (0-249)<br>IPR013785 (0-251)<br>IPR000652 (2-249)<br>IPR022896 (1-249) | 252 |
| MFS transporter | 4ikvA | IPR011701 (1-380)<br>IPR020846 (1-380)<br>IPR036259 (1-380) | 380 |
| Immunoglobulin | 7sbdH | IPR036179 (0-116; 124-199)<br>IPR013783 (0-206)<br>IPR003597 (124-202)<br>IPR007110 (0-115; 121-207)<br>IPR003599 (6-115)<br>IPR013106 (11-114) | 209 |
| Histidine kinase | 8dvqA | IPR003594 (47-156)<br>IPR003594 (47-158)<br>IPR004358 (118-137)<br>IPR004358 (141-155)<br>IPR004358 (101-112)<br>IPR005467 (0-158)<br>IPR036890 (4-159)<br>IPR036890 (3-156) | 166 |
| Alpha/beta hydrolase | 7yiiA | IPR029058 (0-274)<br>IPR000073 (26-265) | 276 |

**Table S13.**
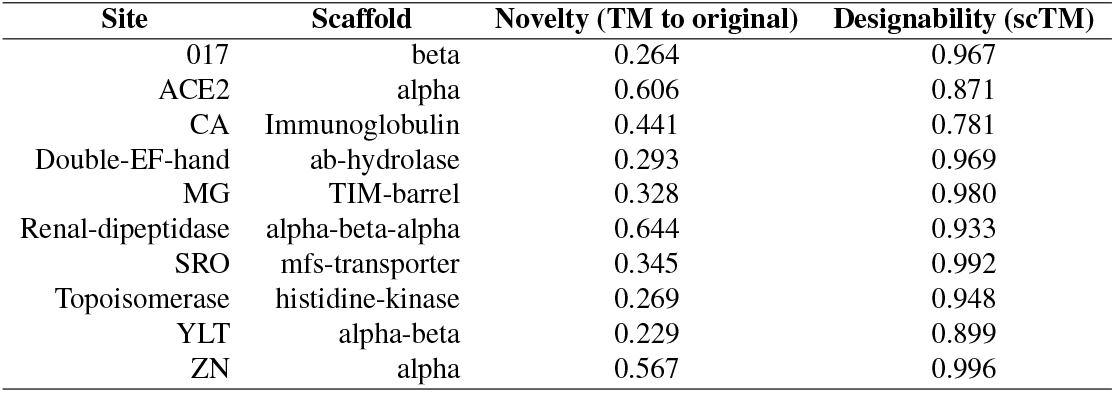
Novelty and designability metrics. Metrics for motif scaffolds shown in Fig. 2C. Novelty is measured by computing the TM-score to the original scaffold from which the motif is derived. Designability is measured by self-consistency TM-score over eight samples by inverse folding with ESM-IF1 and refolding with ESMFold. All designs are distinct from their original scaffolds while retaining high designability.

| Site | Scaffold | Novelty (TM to original) | Designability (scTM) |
| --- | --- | --- | --- |
| 017 | beta | 0.264 | 0.967 |
| ACE2 | alpha | 0.606 | 0.871 |
| CA | Immunoglobulin | 0.441 | 0.781 |
| Double-EF-hand | ab-hydrolase | 0.293 | 0.969 |
| MG | TIM-barrel | 0.328 | 0.980 |
| Renal-dipeptidase | alpha-beta-alpha | 0.644 | 0.933 |
| SRO | mfs-transporter | 0.345 | 0.992 |
| Topoisomerase | histidine-kinase | 0.269 | 0.948 |
| YLT | alpha-beta | 0.229 | 0.899 |
| ZN | alpha | 0.567 | 0.996 |

We also use ESM3 to generate an idealized TIM Barrel with 11-fold symmetry. This generation is undertaken in two steps. First, we derive a secondary structure and function keyword prompt from a reference TIM Barrel (PDB 5EKY). The secondary structure of the reference protein is computed using DSSP and then idealized to construct a prompt for ESM3. To construct the secondary structure prompt, the length of each helix and strand is fixed at 7 residues. Each helix and strand region is then separated by 3 mask tokens, with a mask token appended to the N and C termini of the prompt as well. This yields a secondary structure prompt of total length 159, which is combined with a function keyword prompt derived from the reference protein: keywords are derived from IPR013785 (aldolase-type TIM barrel) and IPR000887 (KDPG/KHG aldolase). ESM3 7B is then used to generate 256 samples with *L* decoding steps and a temperature of 0.7. The design shown is chosen by filtering by ESM3 pTM and visual inspection. In the second step, the secondary structure prompt from the first step is expanded to contain 11 helix-strand subunits, for a total prompt length of 225 residues (4 mask tokens are now appended to the N and C termini, rather than just 1). ESM3 7B is then used to generate 256 samples with *L* decoding steps and a temperature of 0.7, with generations filtered by ESM3 pTM and visual inspection. The generation is evaluated as described above (ESMFold pTM 0.69, scTM mean 0.97, std 0.011). The generation is structurally distinct: a Foldseek search (41) of AlphaFold-DB, ESMAtlas, and PDB in TM-align mode reveals no hit with TM-score greater than .61.

#### A.4. Alignment

##### A.4.1. Algorithm

Since the introduction of RLHF (44) there have been a number of algorithms developed to tune large models trained via unsupervised learning to better follow instructions and generally align their generations to user preferences (45, 46, 101, 102). We use IRPO (Iterative Reasoning Preference Optimization) due to its simplicity in implementation and good performance. The IRPO loss combines supervised finetuning with contrastive learning from preference pairs. IRPO operates on a dataset 𝒟 ∼ (*y*_*w*_, *y*_*l*_, *x*) consisting of prompt *x* and a pair of completions *y*_*w*_ (preferred) and *y*_*l*_ (not preferred). It also operates on two separate models: the reference model *π*_ref_ and the current model *π*_*θ*_. The reference model *π*_ref_ is the fixed base model of the same scale, and the current model *π*_*θ*_ is the model being optimized.

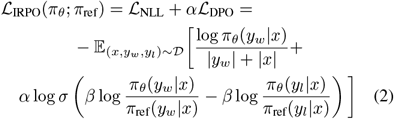

The IRPO loss contains two terms. The ℒ _NLL_ term maximizes the log likelihood of the preferred example normalized by the length of the sequence, providing signal to reinforce the good generations from the model. The ℒ _DPO_ term is the contrastive preference tuning term, which increases the difference in log likelihoods between the preferred and not preferred examples while staying close to the reference model (45). The use of the reference model serves as a regularizer to prevent overfitting to the preference dataset, which can often be small. There are two hyperparameters, *α* and *β. α* weights the relative importance of the supervised with the preference loss and the *β* parameter controls how close we stay to the reference model: the higher the beta, the closer we stay. We minimize this loss with respect to the current model parameters *θ*.

ESM3 is a multi-modal model so the prompt can be any combination of the input tracks of (partial) sequence, structure, and function and the generation y can be any of the output tracks. In our experiments we always generate the amino-acid sequence so this will be our running example from now on. Since an amino-acid sequence *y* can be generated from prompt *x* in many multi-step ways computing the full likelihood *π*(*y* | *x*) would involve integrating over all possible multi-step decoding paths. Since this is intractable, we use a surrogate that mirrors pretraining, shown in Eq. (3) and described below.

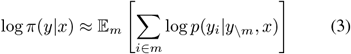

To approximate the likelihood of a generation *y* from prompt *x*, we mask *y* with a mask sampled from a linear noise schedule, prompt ESM3 with {*y*_*\m*_, *x}*, and compute the cross-entropy of ESM3 logits with the masked positions of *y*. During training, the same mask is used to compute the likelihoods for the reference policy vs current policy, as well as for the preferred sample vs non preferred sample.

##### A.4.2. Preference Tuning Intuition

Rearranging the DPO term of the loss function gives some insight into how it finetunes the model for the preference pairs.

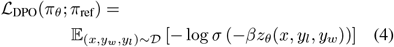

**Figure S17.**
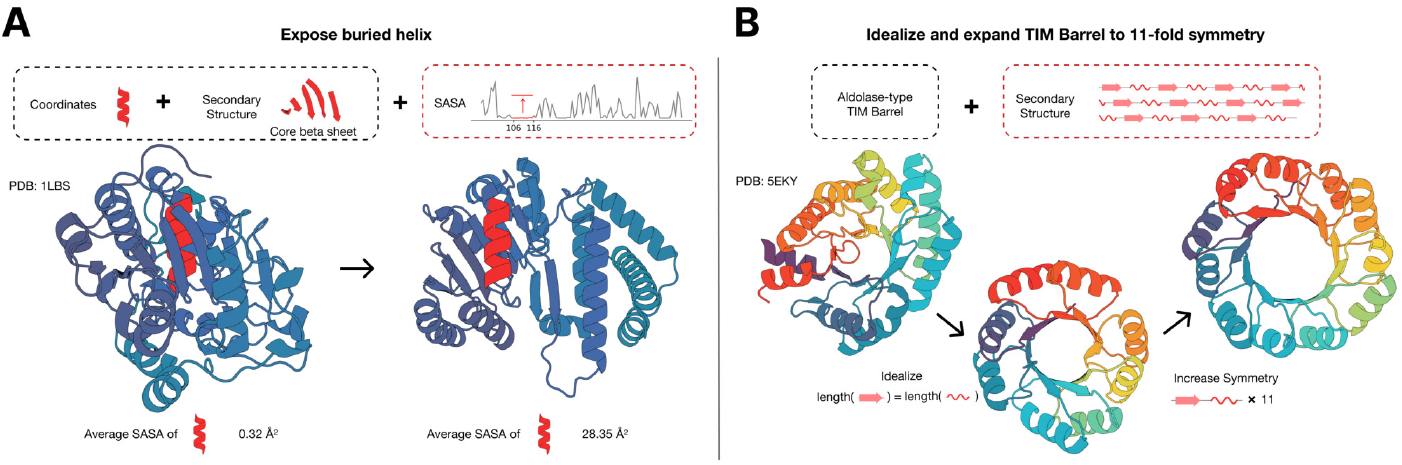
Multimodal protein editing with ESM3. (A) ESM3 exposes a buried helix in an protein while maintaining the alternating alpha-beta sandwich fold of the protein. (B) ESM3 is used in a two-step iterative edit, where first secondary structure prompting and function prompting are used to idealize a reference TIM barrel. Secondary structure prompting is then used to increase the number of subunits in the TIM barrel from 8 to 11.

**Table S14.** Prompt lengths and constraint variation for ESM3 design tasks.

| Task | Prompt Length Variation | Constraint Position Variation |
| --- | --- | --- |
| Fig. 2A (Prompt following for structure coordinates, SS8, SASA, keywords) | No; same as template | No; same as template |
| Fig. 2B (Prompting for out-of-distribution folds) | No; same as template | No; same as template |
| Fig. 2B (Prompting for symmetric proteins) | No; same as template | No; same as template |
| Fig. 2C (Secondary structure + active site prompting) | Yes; sampled uniformly in [150, 400] | Yes; SS spans sampled randomly, active site positions sampled randomly |
| Fig. 2C (Keyword + active site prompting) | No; same as template | Yes; active site positions sampled randomly, keyword positions fixed |
| Fig. 2D (Compress structure) | Yes; reduced to 150 residues | Yes; active site positions sampled randomly, keyword positions fixed |
| Fig. 4A (GFP Design) | No; same as template | No; chromophore positions fixed |
| Fig. S17 (Expose buried helix) | No; same as template | No; helix and core sheet positions held fixed |
| Fig. S17 (Idealize + expand TIM barrel) | Yes; increased to achieve 11-fold symmetry | Yes; SS8 prompt repeated 11x to achieve 11-fold symmetry |

Where

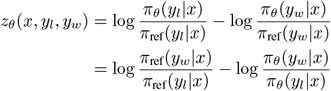

The function *f*(*z*) = −log *σ*(−*βz*) = log(1 + exp(*βz*)) is the softplus function, and is an approximation of the hinge function; in other words *f*(*z*) = *βz* when *z >>* 0 and *f*(*z*) = 0 when *z* ≪ 0. Because of this property, there are two cases. In the case where

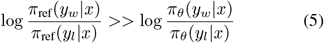

*f*(*z*) is in the linear regime, so the loss function is simply maximizing the likelihood ratio 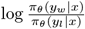. In the case where

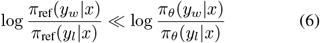

the loss has saturated. This ensures that we do not deviate too far from the reference model.

These dynamics also hold true in the case of ESM3 finetuning. Although we use a surrogate instead of the true likelihood, the loss will increase the surrogate of the preferred pair over the non preferred pair until the current model deviates too much from the reference model.

##### A.4.3. Evaluation Metrics

Possibly the most important part of preference tuning is how the generations are separated into preference pairs. The desired objectives for a generation are quality and correctness. Quality refers to the viability of the sequence to be a stable protein. Correctness refers to the extent to which it follows the given prompt; also called prompt consistency. This section only deals with structure coordinate prompts, so prompt consistency can be measured via constrained site RMSD (cRMSD), which is the RMSD between the backbone coordinates of the prompt and the corresponding coordinates in the predicted structure of the generated sequence. Sequence quality can be measured via predicted-TM (pTM) of a structure predictor on the generated sequence.

As with any metric, especially one which is really a surrogate such as a structure predictor, there is a risk of over optimizing: the model keeps improving the specific metric e.g. in our case pTM but the actual property of interest, the viability of the sequence to be a stable protein, stops correlating with the metric (103). Using orthogonal models to rank our training dataset vs to perform evaluation helps mitigate this.

To create the training datasets, generations are evaluated according to backbone cRMSD and pTM of ESM3 7B to maintain a consistent structure predictor across all datasets. After the preference tuning phase, the generations from the tuned models are evaluated with ESMFold backbone cRMSD and pTM as an orthogonal model. Training on ESM3-scored preference pairs while evaluating using ESM-Fold provides a more objective evaluation.

##### A.4.4. Training Dataset

All ESM3 model scales are trained with the IRPO loss (Eq. (2)) on their respective preconstructed training datasets consisting of structure coordinate prompts and generations. The datasets have 16 generations each for 30,000 prompts from the respective ESM3 model. Preference selection is determined via a threshold of metrics. A sample is considered “good” if it has ESM3 7B pTM *>* 0.8 and backbone cRMSD to its structure prompt *<* 1.5A°.

Each “good” sample is paired with a “bad” sample to create a preference pair. We found that enforcing a gap between metrics of paired generations improves results, so to qualify as a “bad” sample the generation must have a delta pTM = pTMgood − pTMbad *>*= 0.2 and delta backbone cRMSD = cRMSD_good_ − cRMSD_bad_ *<* − 2A°. Each prompt can have multiple preference pairs, and prompts with no valid preference pair are discarded.

The structure prompts are composed of a variety of proteins adapted from our pretraining pipeline. 50% of the prompts are synthetic active sites, while the other 50% are structure coordinates randomly masked with a noise schedule. All of the structure prompts are derived from PDB structures with a temporal cutoff of before May 1st, 2020.

The synthetic active sites are derived by finding sequences from PDB with coordinating residues. For these structures, the amino acid identities are included in the prompt.

The remaining structure track prompts are masked according to a cosine noise schedule. 50% of the noise scheduled prompts are masked in completely random positions, and the other 50% are masked according to an autocorrelation mechanism that prefers sequentially masked positions.

Each model’s training dataset consists of generations of its own reference model. For each prompt, we generate samples from the corresponding ESM3 model scale using iterative decoding with *L/*4 steps, where *L* is the length of the prompt. We anneal the temperature from 1.0 to 0.5 over the decoding steps.

##### A.4.5. Evaluation Dataset: Tertiary Coordination

Tertiary coordination tasks require the generation of proteins which satisfy challenging tertiary interaction constraints. The model is prompted with the sequence and coordinates of a set of residues which are near in 3D space, but distant in sequence. To evaluate performance on these tasks, we curate a dataset of 46 proteins with ligand binding sites from the Biolip dataset (100). All selected proteins were deposited in the PDB after the training set cutoff date (2020-12-01). The coordinating residues shown to the model are given by the ligand binding sites defined in the Biolip dataset (Table S15).

**Table S15.** Tertiary coordination dataset. Selected PDBs and coordinating residues (along with binding ligand) for each protein sample in the tertiary coordination dataset.

| PDB ID | Coordinating Residues | Ligand ID |
| --- | --- | --- |
| 7map | D25 G27 A28 D29 D30 G48 G49 V50 | 017 |
| 7n3u | I305 F310 V313 A326 K328 N376 C379 G382 D386 F433 | 05J |
| 7exd | D103 I104 C107 T108 I174 H176 T182 W306 F309 E313 Y337 | 05X |
| 8gxp | W317 C320 A321 H323 V376 F377 L396 I400 H479 Y502 | 06L |
| 7n4z | M66 C67 R124 L130 C134 Y135 D152 F155 | 08N |
| 7vrd | A40 S41 H161 Q169 E170 E213 D248 D324 K349 H377 R378 S379 K400 | 2PG |
| 7zyk | V53 V66 V116 H160 N161 I174 D175 | ADP |
| 6yj7 | K23 V24 A25 Y45 T46 A47 F115 I128 | AMP |
| 8ppb | H185 F198 K209 Q249 D250 L251 D262 K336 I415 D416 | ATP |
| 7knv | E33 F94 E95 D125 | CA |
| 7xer | Y466 L505 T525 | CLR |
| 7tj6 | F366 G367 T378 R418 | CMP |
| 6xm7 | H167 H218 H284 H476 | CO |
| 7bfr | Q62 X126 H248 | CO3 |
| 6xlr | X272 Y495 H496 H581 | CU |
| 6tnh | N40 A41 S127 T128 Q187 L191 C201 T202 V236 | DGP |
| 7ndr | F73 S101 F102 D103 R106 | EDO |
| 8axy | H68 H109 E144 | FE |
| 7o6c | E62 E107 Q141 | FE2 |
| 8aul | P31 M32 T33 Q106 H185 R237 S319 G320 G321 G342 R343 F369 Y370 | FMN |
| 7vcp | N37 D38 Q54 F97 S98 R159 D160 E214 Y276 W297 | FRU |
| 7b7f | G167 T168 G189 W195 | FUC |
| 8d0w | F73 L136 E137 F329 | GAL |
| 7yua | T13 T14 I15 D40 H85 S86 D87 D110 N290 | GDP |
| 7w1a | L44 Y88 L91 I212 | GMP |
| 7ljn | G71 S72 D91 K236 S253 V254 D309 R310 | GTP |
| 6s4f | Y84 N87 K88 V131 Q132 L133 D155 F157 I276 P309 G310 G313 P314 V317 | KUN |
| 7mg7 | Y12 G98 L99 Y100 A207 D208 G227 R228 | MAN |
| 7qow | D12 T118 E268 | MG |
| 7dmm | E181 E217 D245 D287 | MN |
| 7qoz | G11 G12 I13 Y34 D35 V36 A86 G87 V126 T127 N128 H185 M235 | NAD |
| 7v2r | G89 F93 K98 F101 E121 Y204 E209 F229 | NAI |
| 7a7b | F51 Y128 K165 N166 S167 Y186 R187 I248 G249 A299 | NAP |
| 7pae | M20 L22 L38 V49 I53 C56 K57 R61 Q78 V80 W90 I109 M117 I129 L147 Y149 | O7T |
| 8egy | H82 K83 S186 G230 S231 N232 E345 S368 G369 | PLP |
| 7qow | S65 R129 D273 H465 | PO4 |
| 7wmk | E77 L124 R129 S174 T189 Q191 W241 D304 E306 K349 D410 W411 Y486 | PQQ |
| 7pl9 | D607 A608 Y637 M638 Y705 G706 M735 K736 | RET |
| 7yf2 | G153 E174 L175 L209 N210 L211 Y295 | SAH |
| 7v6j | G207 D230 L231 D250 M251 K264 | SAM |
| 7ys6 | D106 C110 N288 | SRO |
| 6w8m | A22 A23 G70 S110 T111 G112 V113 Y114 | TJY |
| 8g27 | S258 D294 K435 R717 | UDP |
| 7xyk | R24 C170 R190 S191 D193 N201 H231 Y233 | UMP |
| 8g3s | H224 F228 V249 M250 V253 R263 T266 L267 F270 | YLT |
| 8it9 | T92 P93 R96 Y108 L109 K216 V228 S229 H231 H232 | ZL6 |

ESM3 is prompted with the sequence and coordinates of the residues for a particular ligand binding site. We ask ESM3 to generate novel structures by applying multiple transformations to the prompt. The total sequence length is sampled evenly to be 150, 250, or 350 residues (regardless of the original sequence length). Next, we define a contiguous span of coordinating residues to be prompt residues with fewer than 5 sequence positions between them. The order and the distance between contiguous spans of residues is shuffled. Together, this ensures that, for example, the original protein will no longer satisfy the prompt. We consider a generation a success if backbone cRMSD *<* 1.5A° pTM *>* 0.8.

We construct a total of 1024 prompts for each ligand and generate a completion for each prompt with the model we are evaluating. We report Pass@128, which is an estimate for the fraction of ligands with at least one successful completion after 128 prompts per ligand. We estimate this using an unbiased estimator (Chen et al. (104), Page 3) using the success rate over 1024 prompts. We visualize randomly selected successful generations for both the base model and finetuned model in Fig. S19.

We evaluate RFDiffusion on the tertiary coordination benchmark following a similar methodology. We use the RFDiffusion model finetuned for active site scaffolding. For each ligand, we prompt the model with the same 1024 prompts as ESM3 (maintaining identical protein lengths and active site positions) and generate one structure per prompt using the default parameters provided in the RFDiffusion repository. To generate sequences, we inverse fold each structure once with ProteinMPNN using temperature 0.1, yielding 1024 sequences per ligand. This approach ensures that RFDiffusion and ESM3 are afforded the same sampling budget. These sequences are then scored using the same approach as for ESM3-generated sequences. Pass@128 and the the fraction of tasks solved with at least 8, 16, and 32 distinct solutions (clustered at TM *>* 0.8) are reported in Table S16.

##### A.4.6 Supervised Finetuning

To judge the value of preference tuning, we also train a supervised finetuning (SFT) baseline where we finetune the model to increase likelihood of the high quality samples without the preference tuning loss. The 1.4B, 7B, and 98B models solve 14.2%, 33.7%, and 44.6% of tertiary coordination tasks at 128 generations, respectively, which improves upon the base models but is much lower than their corresponding preference tuned versions.

##### A.4.7 Training Hyperparameters

Each IRPO model is trained for 1000 steps using RMSProp. The learning rates are 1e-5, 1e-5, and 5e-6 for the 1.4B, 7B, and 98B, respectively, annealed using a cosine schedule after a 150 step warmup. Gradient norms are clipped to 1.0.

For all IRPO runs *β* = 0.05 and *α* = 0.8. The SFT baseline uses the same hyperparameters, but with *α* = 0.0 to disregard the preference tuning term.

**Figure S18.**
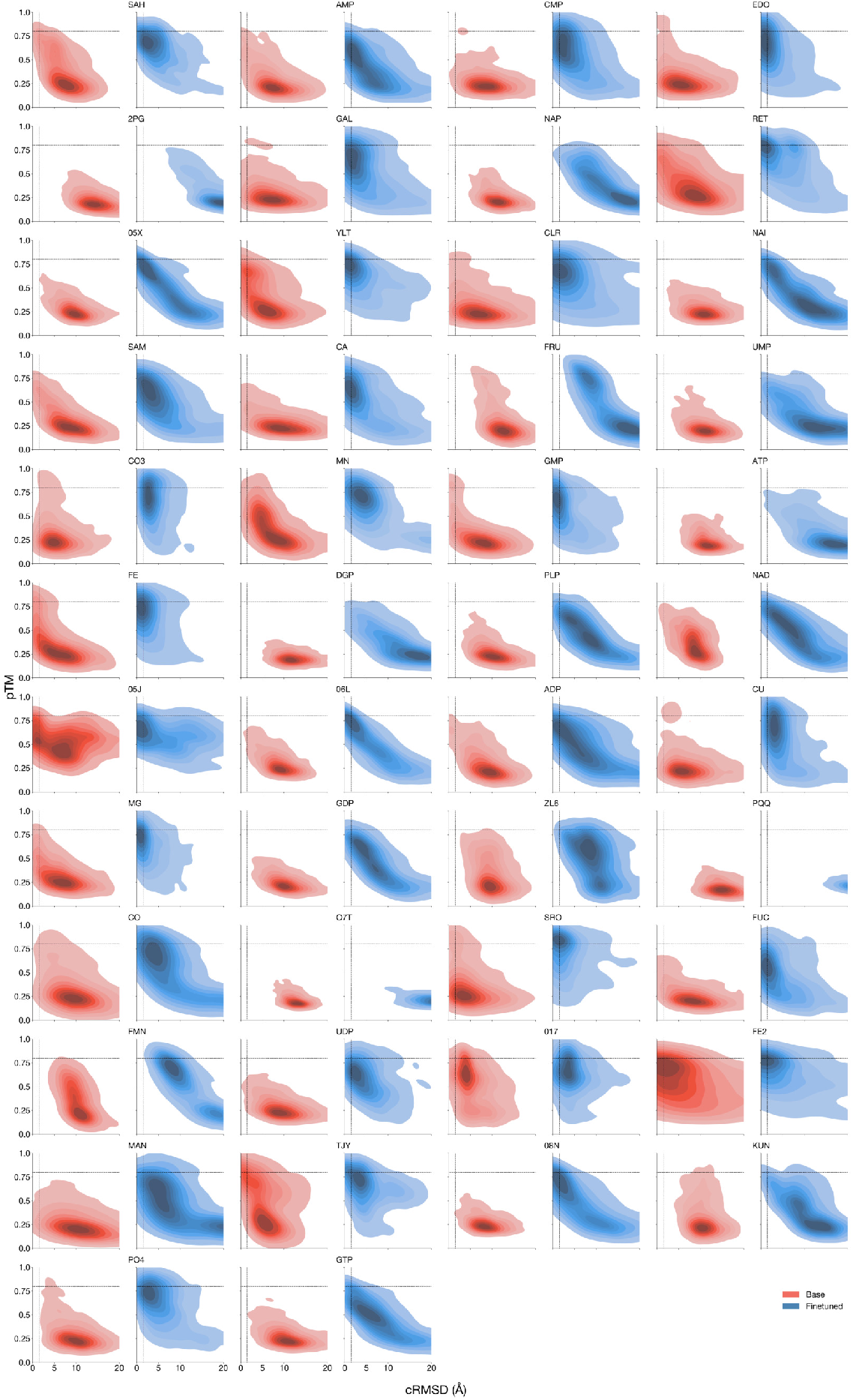
Alignment improves model generations. pTM, backbone cRMSD distributions of generations from the 98B base model and aligned model for all ligands in the tertiary coordination dataset. Each ligand/model pair has 1024 generations.

**Figure S19.**
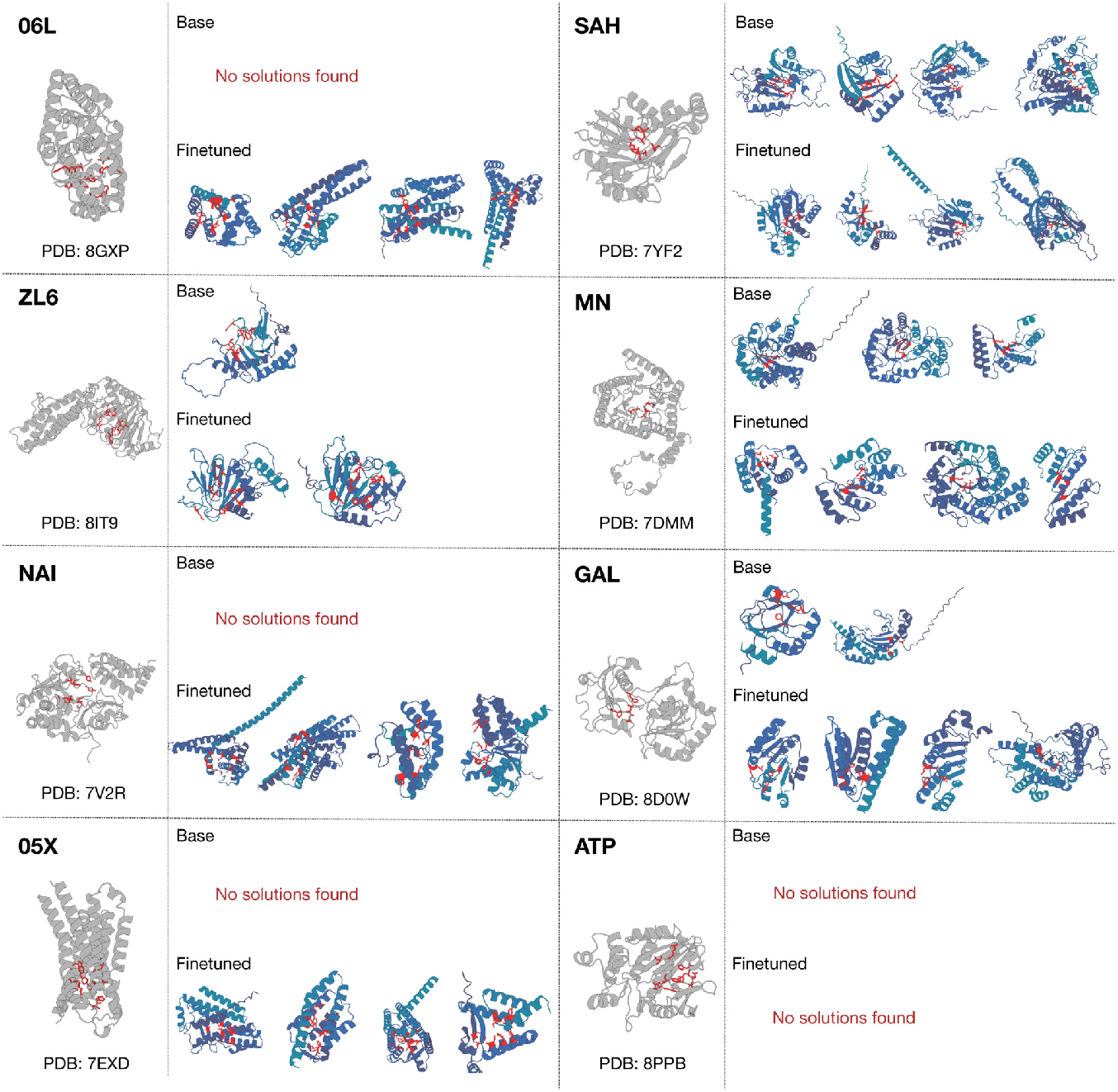
Randomly selected successful generations from the base and finetuned 98B parameter models. A random sample of ligands is selected and visualized with the ground truth PDB chain from which the ligand was taken. Solutions produced by ESM3 are diverse, and the finetuned model gives significantly more successes (out of 1024 total samples).

**Table S16.**
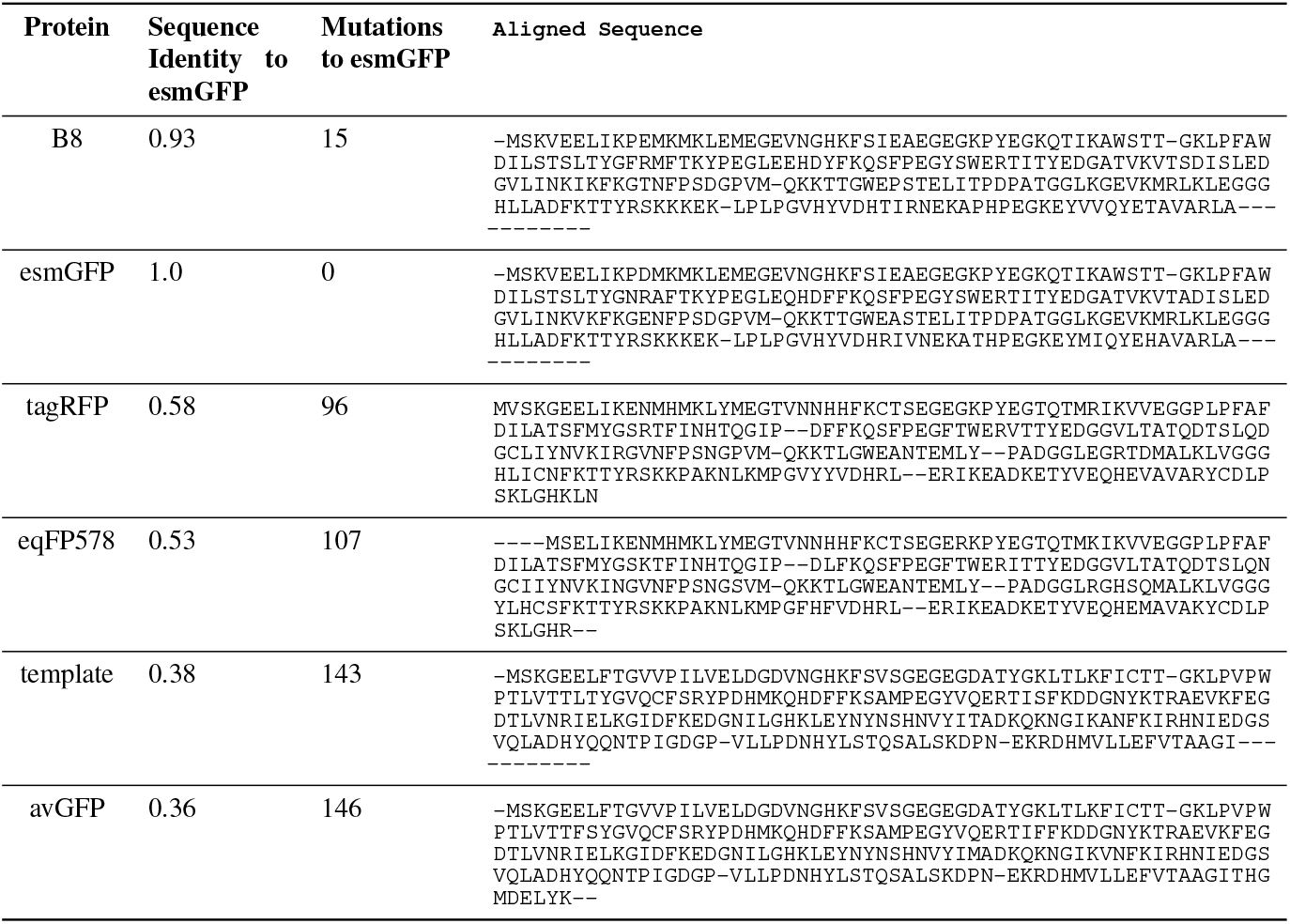
Tertiary coordination benchmark. Fraction of tertiary coordination tasks solved with 128 generations (Pass@128; 2 s.d.), and fraction of tasks with at least 8, 16, and 32 distinct solutions (clustered at TM *>* 0.8) given a budget of 1024 generations. ESM3 models can generate sequences either directly from a prompt (“Without CoT”) or via a chain of thought by first generating structure tokens, then sequence (“With CoT”). Chain of thought improves success rates in finetuned models. The ESM3 98B finetuned model with chain of thought achieves the strongest performance.

#### A.5 GFP

ESM3 generates a dim distant GFP, B8, and a bright distant protein, esmGFP. Details are provided below on computational methods, experimental protocols, results, and post-experiment analyses.

##### A.5.1 Generation and Selection

The base ESM3 7B model generates candidate GFP designs for laboratory testing using a single prompt and a chain of thought over sequence and structure tokens. Candidates are filtered and ranked by metrics at several steps in the process. Experiment 1 tests candidates across a range of sequence identity to a template, yielding multiple GFPs including dim hit B8. Experiment 2 consists of designs starting a chain of thought from the sequence of B8, yielding numerous bright GFPs including C10 which we term esmGFP. This section details the computational protocol that generated and selected candidate GFP designs for Experiments 1 and 2, shown in Fig. 4B. Protocols, metrics, and selection conventions are separately introduced and then synthesized in descriptions of the two experiments, at the end of the section.

###### A.5.1.1 Model

All candidate GFP designs were created using the base ESM3 7B model with no finetuning. Throughout generation, the model is prevented from decoding cysteine residues.

###### A.5.1.2 Prompt

All candidate GFP designs in Experiment 1 are produced with a chain of thought beginning from a single prompt. The goal of the prompt is to capture essential residue identities and structural features needed for chromophore formation and fluorescence, leaving other degrees of freedom open for the model to generate diverse designs.

###### Template

We prompt ESM3 with a minimal set of sequence and structure information from 16 residues near the chromophore formation site. We use a pre-cyclized intermediate crystal structure derived from Barondeau et al. (53), PDB ID 1QY3, as a template. Since this template contains the chromophore maturation slowing mutation R96A, for the prompt we use the unmutated Arg96, constructing coordinates for the unmutated residue by superimposing the backbone atoms of 1GFL (105) onto 1QY3 and replacing 1QY3’s A96 with 1GFL’s R96. We subsequently refer to the full sequence and structure of 1QY3 with mutation A96R as 1QY3 A96R or the template.

###### Sequence prompt

The sequence portion of our prompt consists of 7 template residues: Met1, Thr62, Thr65, Tyr66, Gly67, Arg96, and Glu222. Residues 65-67 form the chromophore. Met1 ensures proper start codon placement. Residues 62, 96, and 222 are described in (53) and other works to have key catalytic roles in chromophore formation.

###### Structure prompt

The structure portion of our prompt consists of structure tokens and backbone atomic coordinates taken from 16 template residues at positions 96, 222, and 58-71 (inclusive) which roughly captures the central alpha helix. The unique geometry of the central alpha helix is known to be crucial for chromophore formation (53).

All other positions and tracks in the prompt are masked. The overall prompt length is 229, matching that of the template. Residue indices are contiguous and begin from 1.

###### A.5.1.3 Joint Sequence Structure Optimization

We employ the following procedure to jointly optimize the sequence and structure of designs throughout our experiments: While annealing temperature linearly from 1 to 0, we perform multiple iterations of first predicting the structure of a designed sequence and subsequently Gibbs sampling each position in the sequence for that predicted structure. In algorithmic form:

###### Algorithm 15

`gibbs_seq_given_struct`

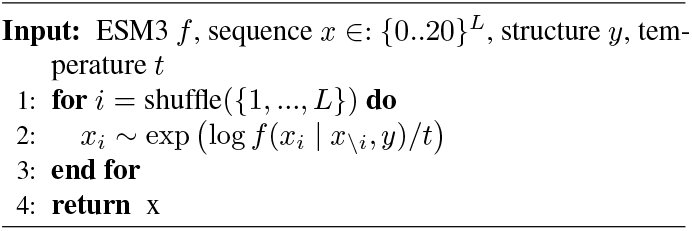

###### Algorithm 16

`joint_optimize`

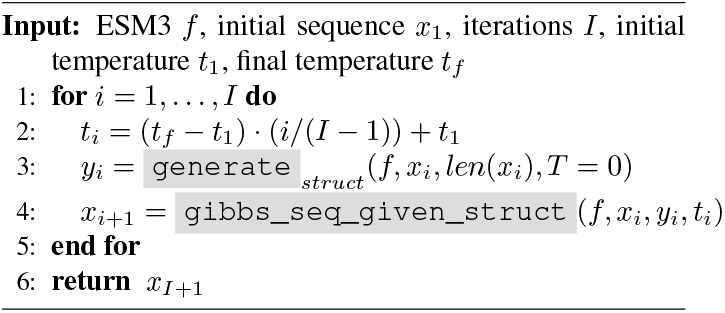

Three variants of `gibbs_seq_given_struct` in `joint_optimize` were employed for Experiments 1 and 2. Joint optimization occasionally produces repetitive spans of amino acids when temperature is annealed to low values. Variant 1 and 2 are intended to address this, in differing ways. Variant 3 is an experiment in biasing the logits with a PSSM of known natural GFPs. Half of the candidates in Experiment 2 were produced using Variant 3. This half did not include esmGFP, which was generated by Variant 2 of joint optimization.

1. **Variant 1: Negative Local Sequence Guidance** We bias the logits of the model away from those produced just from a highly local span of the sequence. Specifically, we use classifier-free guidance (106):

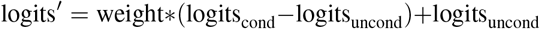

but push away from the logits produced by inputting just 7 residues centered on the position being sampled, with weight 2 and nothing else. All other sequence positions and all other model inputs are left blank.

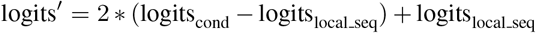
2. **Variant 2: Max Decoding Entropy Threshold** We optionally skip resampling of sequence during Gibbs sampling at positions whose entropy over sequence tokens exceeds a user specified threshold.
3. **Variant 3: PSSM Bias** In Experiment 2 only, we experiment with both including and excluding a PSSM-based bias during Gibbs sequence sampling. Specifically, we add a PSSM constructed from 71 natural GFPs (see Appendix A.5.1.4 for details) directly to the sequence output logits of the model, with a user-specific weight. esmGFP did not use this option; it was produced with weight 0.

###### A.5.1.4 Metrics

GFP designs are produced and scored by a number of ESM3-derived and independent metrics. Unless otherwise noted, designed structures are predicted using ESM3 with only sequence as input, using iterative decoding of structure tokens with temperature 0 and subsequent decoding of backbone coordinates with an older version of the structure token decoder.

The following is an exhaustive list of metrics used. An exact break down of where and how specific metrics are used can be found in Appendix A.5.1.5, Appendix A.5.1.6 and Appendix A.5.1.7.

**Template Chromophore Site RMSD** is calculated via an optimal alignment (107) of N, C, CA, and inferred CB atoms at positions 62, 65, 66, 67, 96, and 222 in the predicted structure of a design and the template (crystal) structure.

**Template Helix RMSD** is calculated in the same way, but for N, C, CA atoms only, at design and template positions 58-71 (inclusive).

**1EMA Helix RMSD** is a metric proposed in (108). An RMSD is calculated between alpha helix residues in the predicted designed structure and a specific crystal structure of avGFP, PDB ID `1EMA`. Our calculation differs slightly from (108). We calculate RMSD for N, C, CA and inferred O atoms, and consider only positions 60-64 and 68-74 (both ranges inclusive) to exclude chromophore positions 65-67.

**Sequence Pseudo-perplexity** is calculated as defined in (109). Given a protein sequence, positions are masked one at a time, negative log-likelihoods of input tokens at masked positions are averaged across all positions in the sequence, and the result is exponentiated.

**Round-trip Perplexity** is calculated for a designed sequence via predicting its structure with ESM3, and then evaluating the perplexity of the sequence given that predicted structure under a single forward pass of ESM3.

**N-gram Score** is calculated as the *E*_ngram_ term defined in (11). This score assesses the divergence between the N-gram frequencies of residues in the designed sequence and those found in a background distribution, derived from UniRef50 2018 03. Specifically, for a function ngram_*i*_ that takes in a sequence *x* and an N-gram order *i*, and a precomputed distribuion of background N-gram frequencies ngram_*i,bg*_, the score is calculated as:

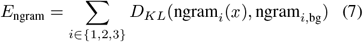

**PSSM** A position-specific scoring matrix (PSSM) is constructed from a MSA of 71 natural GFPs (110). Specifically, at positions aligned to our template, frequencies for the 20 canonical amino acids (excluding gaps) are transformed to log odds via dividing by the uniform background (*p*(*aa*) = 0.05), adding an epsilon of 1e-9, and applying log base 2. This produces a matrix of scores of size 229 × 20.

**PSSM score** We extract from the PSSM values at (position, amino acid) pairs occurring in an input sequence. These are averaged to produce a score.

**N-terminus Coil Count** is metric intended to measure structural disorder at the N-terminus of a design. We observed that predicted structures have various levels of disorder in this region. To quantify it for possible filtering, we apply mkdssp (83) to the ESM3-predicted structure of a design, and record how many of the first 12 positions are reported as having SS8 labels in {S,T,C}.

###### A.5.1.5 Selection Criteria

Among Experiment 1 and 2, designs are selected for testing by first applying a set of filters, and then selecting the top-N designs according to a score-based ranking. Scores are calculated by summing the values of several metrics, which are each normalized across designs to have zero mean and unit variance and which are negated when appropriate so that lower values are always better.

###### Common Filters

The following filters are applied in both Experiments 1 and 2.

- Template Chromophore Site RMSD *<*1.5A°
- Template Helix RMSD *<*1.5A°
- N-gram Score *<*5

###### Common Score Terms

The following score terms are used in both Experiments 1 and 2.

- Sequence Pseudo-perplexity
- Round-trip Perplexity
- ESM3 pTM

###### A.5.1.6 Generation and Selection of Designs for Experiment 1

In this experiment, we generate a set of GFP designs for experimental testing with a range of sequence identities to our template. Designs are generated by a chain of thought: from the prompt, ESM3 decodes all masked structure tokens, then all masked sequence tokens. Lastly, sequence and structure tokens are jointly optimized.

###### Initial Generation

Starting from the prompt, we first generate 38k structures by decoding masked structure tokens one at a time using a fixed temperature sampled uniformly from the range (0, 1.25) for each generation. To focus compute on the most promising structures, we filter according to Template Chromophore Site RMSD *<*1A°, yielding 24k selected structures. We next generate ≈ 4 sequences for each structure with a temperature uniformly sampled from the range (0, 0.6), yielding 92k total sequences.

###### Selection

We select a subset of promising initial generations for further optimization by applying Common Filters with N-gram score’s threshold modified to *<*5.5, ranking designs according to {Common Score Terms, mean ESM3 pLDDT, mean ESMFold pLDDT, and ESMFold pTM}, and selecting the best 40 designs in each interval of 0.1 sequence identity to the template sequence in [0.2, 1.0], 320 in total.

###### Joint Sequence Structure Optimization

We then jointly optimize the sequence and structure of designs. Using 30 iterations in each case, we run 5 seeds of optimization with max decoding entropy threshold = 1.5 and 2 seeds of optimization with negative local sequence guidance = 2.0, yielding 67k total designs. Designs from every iteration are included in this pool.

###### Selection

To select a set of designs for laboratory testing, we apply {Common Filters, N-terminus Coil Count *<*6}, rank designs according to {Common Score Terms, ESMFold pTM, 15 * PSSM Score}, and select the best 88 designs across 8 buckets of sequence identity to our template among intervals of width 0.1 in range [0.2, 1].

###### A.5.1.7 Generation and Selection of Designs for Experiment 2

In this experiment, we perform further refinement of the dim, distant GFP found in Experiment 1, B10. To produce a diversity of designs, we sweep over a number of settings: two variations of refinement are performed, and 2 selection protocols are used.

###### Local Joint Optimization

Starting from our dim GFP design, B10, we perform `joint_optimize` using a full grid sweep of the following sets of settings: Initial temperatures {0.001, 0.01, 0.05, 0.1, 0.5}, PSSM bias weights {0, 0.01, 0.05, 0.1, 0.5}, Max decoding entropy thresholds {0.8, 1, 1.25, 1.5, 2.0}. For each unique settings combination, we use 20 iterations of optimization with 3 seeds, continuing the final step of Gibbs sampling until convergence. After accounting for some distributed system machine failures, this yields 6.3k total candidate designs.

###### Selection

We select two sets of 45 designs for laboratory testing via two filters and a shared set of ranking criteria.

1. **Set 1:** We filter according to {PSSM Bias ≠ 0, Common Filters, RMSD to starting structure *<*1A°, identity to starting sequence in (0.7, 1.0)}.
2. **Set 2:** We filter according to {PSSM Bias = 0 (no bias), Common Filters, RMSD to starting structure *<*1A°, Identity to starting sequence in (0.9, 1.0)}. esmGFP comes from this pool.

For each set, we rank according to Common Score Terms, 8 * PSSM Score, 15 * 1EMA Helix RMSD and select 45 designs each for testing.

##### A.5.2 Experimental Methods and Data Analysis

###### A.5.2.1 Strains and plasmids

We designed a custom bacterial expression vector containing an Ampicillin-resistance gene, the BBa R0040 TetR promoter, the BBa B0015 terminator, and a Bsa-I golden gate site between the promoter and terminator. GFP designs were codon optimized for *E. coli* expression, compatible golden gate overhangs were added, and sequences were ordered from IDT (Integrated Device Technology Inc.). They were then cloned by golden gate assembly into the vector. We evaluated our GFP designs in the *E. coli* host Mach1 (Invitrogen).

###### A.5.2.2 Fluorescence assays of GFP designs

To evaluate the fluorescence of our GFP designs, we transformed our designs into Mach1 cells. For each of two replicates of a design, a colony was seeded into a 1 mL TB culture containing 50 μg/mL carbenicillin. Cultures were grown in 96 deep well blocks at 37 °C in an Infors HT Multitron Shaker with a shaking speed of 1000 RPM for 24 hours. After 24 hours, 1 μL of the cultures were diluted in 200 μl of 0.2 μm filtered DPBS.

Fluorescence intensity of the samples was then quantified at the single cell level using a NovoCyte Quanteon Flow Cytometer (Fig. S20).

The remaining cultures were spun down at 4000 g for 10 minutes, resuspended and lysed with 300 μL lysis buffer (1x bugbuster, 500 mM NaCl, 20 mM Tris-HCl pH 8, 10% glycerol, cOmplete™, EDTA-free Protease Inhibitor Cocktail), incubated at room temperature on a Belly Dancer Orbital Shaker for 10 minutes, and lysate clarified by centrifugation at 4000 g for 20 minutes. 100-120 μl lysate was transferred to a 96 well black clear-bottom plate, and GFP fluorescence was measured using a Tecan Spark Reader. Fluorescence emission was captured at 515 nm with a 10 nm bandwidth and excited with 485 nm with a 10 nm bandwidth. Absorbance was captured at 280 nm with a 3.5 nm bandwidth to assess total protein content per well. For longer time points, plates containing lysate were sealed and incubated at 37°C for up to 7 days prior to measuring fluorescence. GFP fluorescence values were first ratio normalized within a well by their absorbance at 280 nm, and then further ratio normalized across wells using the measured values from a negative control *E. coli* containing vector without GFP. Data from two replicates was then averaged for (Fig. 4B bottom) and (Fig. 4C).

Overview photos of the plates (Fig. 4B top) were taken with an iPhone 12 mini under blue light illumination from an Invitrogen Safe Imager 2.0 Blue Light Transilluminator.

For excitation spectra, emission was captured at 570 nm with a 50 nm bandwidth, while the excitation wavelength was varied from 350 to 520 nm with a 10 nm bandwidth. For emission spectra, an excitation wavelength of 430 nm was used with a 50 nm bandwidth, while emission was captured at varying wavelengths from 480 to 650 nm with a 10 nm bandwidth. Excitation and emission spectra were normalized by their maximum values (Fig. 4C).

###### A.5.2.3 Additional GFP experiments

Plate overview photographs (Fig. 4B top) were taken over two weeks since the initial lysate was created and over one week after the final plate reader quantification was done, and so possibly show additional brightness from slow chromophore maturing designs. We observed some low level contamination of wells H11 (vector with no GFP or designs) and H12 (lysis buffer only) in the photograph of Experiment 1 (Fig. 4B top left). Some of this contamination is already visible in well H12 during the initial plate reader quantification (Fig. 4B bottom left). To address potential contamination concerns we performed an additional replication of B8 and observed a similar level of brightness to Experiment 1 (50x less bright than natural GFPs) (Fig. S21).

**Figure S20.**
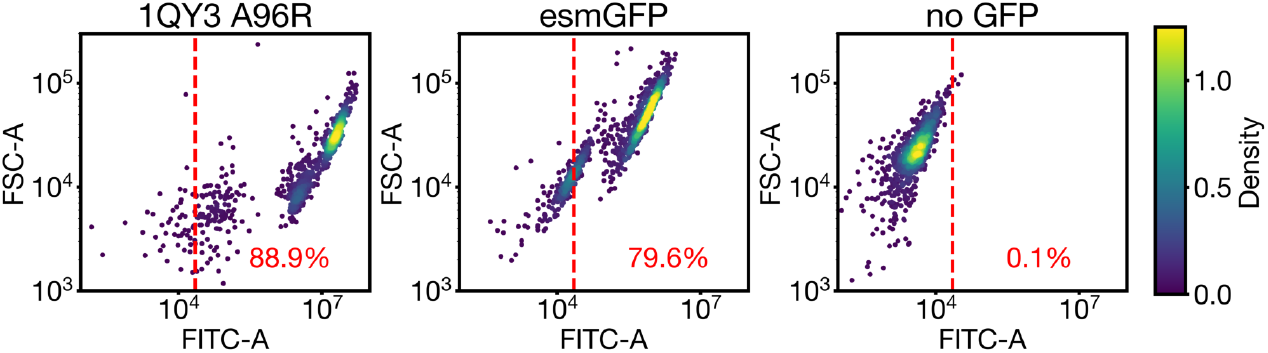
Flow cytometry data confirms cells expressing esmGFP can be detected at the single cell level. Forward Scatter-Area (FSC-A), a measure of cell size vs Fluorescein Isothiocyanate-Area (FITC-A), a measure of GFP-like fluorescent signal, for expressing 1QY3 A96R, esmGFP, and a negative control that does not express any GFP. A gate was set at the 99.9% quantile for the negative control data, and the fraction of cells passing the gate were quantified for each sample.

**Figure S21.**
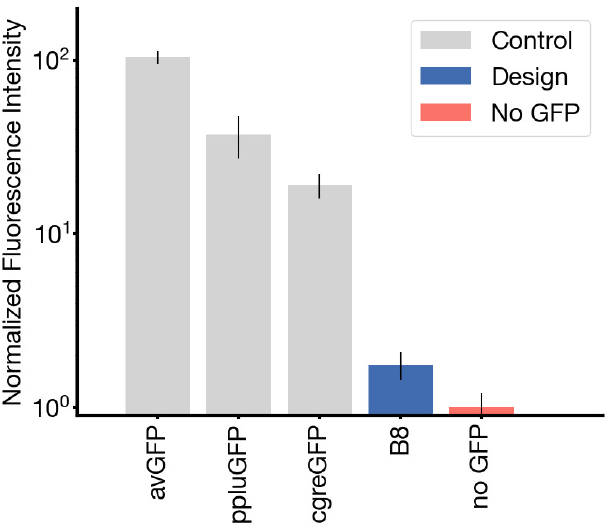
Replication of design B8 and select controls. Results are averages of eight wells across two plates.

Chromophore knockout versions of 1QY3 A96R and esmGFP were created through additional T65G and Y66G mutations. These variants, along with 1QY3 and esmGFP, were synthesized and measured as part of an independent replicate performed by Genscript following the *E. Coli* based fluorescent plate reader assay described above. Normalization was performed with an OD600 measurement of the cells prior to lysis. Analysis otherwise proceeded as above. Two replicates were performed for each design and results were averaged. Chromophore knockout reduced fluorescence to background levels (Fig. S22).

**Figure S22.**
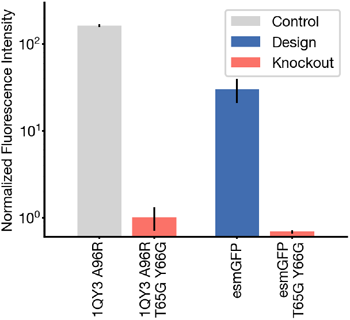
Chromophore knockout mutations. T65G and Y66G reduces fluorescence of both 1QY3 A96R and esmGFP to background levels.

##### A.5.3 Sequence searches and comparisons

###### BLAST nr search

esmGFP’s sequence was searched with BLAST’s online server using the non-redundant sequences database `nr` with all default settings. tagRFP’s sequence was taken from the top hit. The exact top hit found was `TagRFP [Cloning vector pLX-B2-TagRFP-T`, Sequence ID `ASG92118.1` and is shown in its entirety in Table S17.

###### Train set search

MMseqs2 (80), version 15.6f452, was used to search all datasets that ESM3 was trained on at the maximum available expansion level; for cluster resampling datasets all cluster members are searched, not just cluster centers. The goal is to search against every possible sequence that ESM3 may have seen during pretraining. Settings are selected for conducting a high sensitivity search: `-s 6 -a --max-seqs 10000`.

###### A.5.3.2 Sequence Identity calculations

To calculate sequence identities involving the two highlighted GFP designs (B8, esmGFP) and select reference proteins, the following procedure is used. MAFFT (111) v7.525 is applied with all default settings to the sequences of B8, esmGFP, the top tagRFP sequence found by BLAST, eqFP578 (from FPBase (112)), the template (PDB ID 1QY3, with mutation A96R), and avGFP (from FPBase). Identities between two sequences are calculated as the number of matching non-gap residues at aligned positions divided by the minimum non-gapped length of the query and target protein. This is the same sequence identity formula used in Appendix A.5.4. Aligned sequences and identities and mutation counts to esmGFP are provided in Table S17.

###### A.5.3.3 Inner-barrel mutation count

Positions in esmGFP are described as internal if they have SASA *<* 5 in their predicted structure. SASA is calculated as in Appendix A.2.1.6) from the all-atom structure of esmGFP, predicted with ESM3 7B.

##### A.5.4 Phylogenetic Analysis

Sequences and metadata of natural and designed fluorescent proteins were obtained from FPBase (112). An initial set of 1000 proteins was filtered according to the following criteria: presence of a specified parent organism, an amino acid sequence between 200 and 300 residues long, a specified emission maximum, and no cofactors. NCBI taxonomy database was used to obtain taxonomic information about each species. These sequences were further filtered according to keep those that had species found by NCBI and were Eukaryotic but not from Chlorophyta (to exclude Channel-rhodopsin like proteins). The 648 sequences that passed these criteria, along with the sequence for esmGFP, were aligned to a multiple sequence alignement using MAFFT and sequence idenity was computed between each pair of sequences as described above. All pairs within and across taxa were considered for (Fig. 4F). All designed sequences were considered to belong to the species annotated as their parent organism.

**Figure S23.**
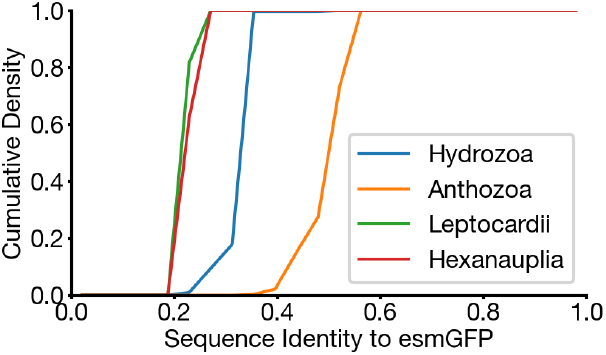
Sequence identity of esmGFP. Comparison with natural and designed GFPs from the four major classes found in nature.

All 648 used sequences belonged to the Leptocardii (e.g. laGFP), Hexanauplia (e.g. ppluGFP), Hydrozoa (e.g. avGFP), or Anthrozoa (e.g. efasGFP) classes. The sequence identity of esmGFP was computed to each protein in these classes (Fig. S23). esmGFP was found to be closest to Anthrozoan GFPs (average sequence identity 51.4%) but also shares some sequence identity to Hydrozoan GFPs (average sequence identity 33.4%).

To estimate the millions of years of evolutionary distance by time between esmGFP and known fluorescent proteins we built an estimator to go from sequence identity between pairs of GFPs to millions of years (MY) apart. We used the following six Anthozoan species *Acropora millepora, Ricordea florida, Montastraea cavernosa, Porites porites, Discosoma sp*., *Eusmilia fastigiata* along with the six GFPs amilGFP, rfloGFP, mcavGFP, pporGFP, dis3GFP, efasGFP respectively. These species and GFPs were chosen because they were annotated in both a recent time calibrated phylogenetic analysis of the Anthozoans (57) and a recent study of GFPs (48). Each of these species contains multiple GFP like sequences including red and cyan FPs. These particular GFPs were chosen as they were annotated to be the main GFP in each species. The millions of years between each species was estimated as twice the millions of years to the last common ancestor annotated in the time calibrated phylogenetic analysis. Using statsmodels (113), a line of best fit was fit between MY and sequence identity. The line was required to pass through a sequence identity of 1.0 and 0 MY. The MY to esmGFP was then estimated using this line and the sequence identity of esmGFP to the nearest known protein.

#### A.6 OPEN MODEL

We are releasing the ESM3 source code and model weights of an open model, ESM3-open. ESM3-open is a 1.4B-parameter model we trained without OAS antibody sequences and with precautionary risk mitigations for release to the academic research community.

As part of this release, we follow guidance from the Principles for the Responsible Development of AI for Biological Design (114). We adopted precautionary risk mitigations, described in Appendix A.6.1, and performed risk evaluations, detailed in Appendix A.6.2. Additionally we conducted a review of the risks and benefits of releasing ESM3-open with experts from the scientific community. We provided reviewers access to ESM3-open, along with a detailed technical report on our risk evaluations. We received unanimous feedback from our reviewers that the benefits of releasing the model greatly outweigh any potential risks.

We see this release as a first step and plan to work with the scientific community to continue to improve processes around responsible development. Open models enable the scientific community to better understand and reduce any potential risks of biological design tools. As our understanding develops alongside the capabilities of future models, we plan to continuously improve our evaluation frameworks, safeguards, and mitigation strategies.

##### A.6.1 ESM3-open Mitigations

As a precaution, we filtered the training data of ESM3-open to minimize model performance on sequences of potential concern while otherwise maintaining performance. We also removed the capability for the model to follow prompts related to viruses and toxins.

###### Filtering sequences of potential concern

Previous work has shown that the performance of protein language models is closely related to the number of similar sequences present in the training data (5). We therefore removed sequences aligned to potentially-concerning proteins from the training data in order to reduce the capability of ESM3-open on these sequences.

We identified and removed sequences unique to viruses, as well as viral and non-viral sequences from the Select Agents and Toxins List (115) maintained by the CDC and USDA. The U.S. Department of Health & Human Services recommends filtering based on the Select Agents list as part of their Screening Framework Guidance for Providers and Users of Synthetic Nucleic Acids (116).

To filter data, we create two denylists: the Viral Denylist and the Select Agent Denylist. We then remove all sequences from the training set that are detected to align to those in the denylists by MMseqs2 at or above a given sequence identity threshold.

To create the Viral Denylist, we identify ∼ 4M sequences that are annotated as viral in UniProt and align almost exclusively to other viral sequences in UniProt. This gives us a procedure that removes viral proteins with both high sensitivity and specificity (as measured by UniProt taxonomic annotations). To create the Select Agents Denylist we identify all sequences in UniProt belonging to organisms on the Select Agents and Toxins List (115). This process gives us 147K non-viral sequences and 40K additional viral sequences.

For each denylist, MMseqs was used to query against the full set of training databases, (including PDB, UniRef, MGnify, and JGI) and all hits were removed from the training set. This filter removes a total of 10.6M sequences across all training sets.

###### Removal of keywords of concern

There are a number of keyword prompts associated with viruses and toxins that we aim to remove. We first identify a list of harmful keywords with the following steps:

1. We curate a list of filter terms associated with viruses and toxins. The full filter term list is available upon request.
2. We then identify all InterPro tags whose free-text term names contain at least one of the filter terms.
3. We identify keywords that are associated with flagged InterPro tags but that are not associated with nonflagged InterPro tags. We remove those keywords. Keywords which are associated with both flagged and non-flagged InterPro tags (e.g. “extracellular region”) are not removed.
4. We additionally remove all keywords that themselves directly contain one of the filter terms

Of the original 68,103 keywords that ESM3 is trained with, this filter removes a total of 9,462 (14%), creating a new vocabulary of 58,641 keywords.

The function vocabulary is defined via vectors representing Term Frequency Inverse Document Frequency (TF-IDF) which are then tokenized using Locality Sensitive Hashing (LSH), as previously described in Appendix A.1.8. To remove flagged keywords, they are first removed from the TF-IDF vocabulary by removing the entries corresponding to flagged keywords. This reduces the TF-IDF vector size to 58,641. The LSH tokenization is defined by 64 hyperplanes, each defined in the TF-IDF space, i.e. a Euclidean space with one dimension per keyword. We redefine the hyperplanes to be in the reduced space by removing the dimensions corresponding to the flagged keywords. This permanently removes the information required for tokenization of the flagged keywords. This mitigation is highly selective and does not change the tokenization for any non-flagged keywords.

**Table S17.**
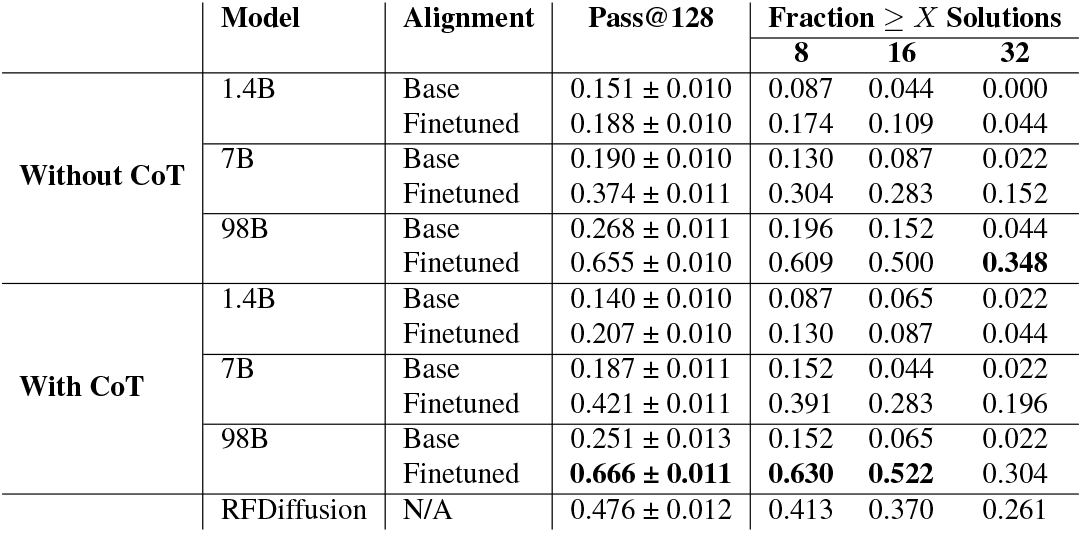
Multiple sequence alignment of select GFP designs (B8, esmGFP) and reference proteins. Template is the full sequence of our template structure (PDB ID 1QY3), with chromophore slowing mutation A96R removed. tagRFP is the full sequence of the top hit returned by BLAST search of the nonredundant database nr, avGFP and eqFP578 are from FPBase. Sequence identities for GFP designs are in general calculated as the number of non-gap matches at aligned positions, divided by the minimum length of the query and target ungapped sequences. Here, only sequence identities to esmGFP are shown. Similarly, the number of mutations to esmGFP are calculated as the number of mismatches at aligned positions where esmGFP does not have a gap.

**Figure S24.**
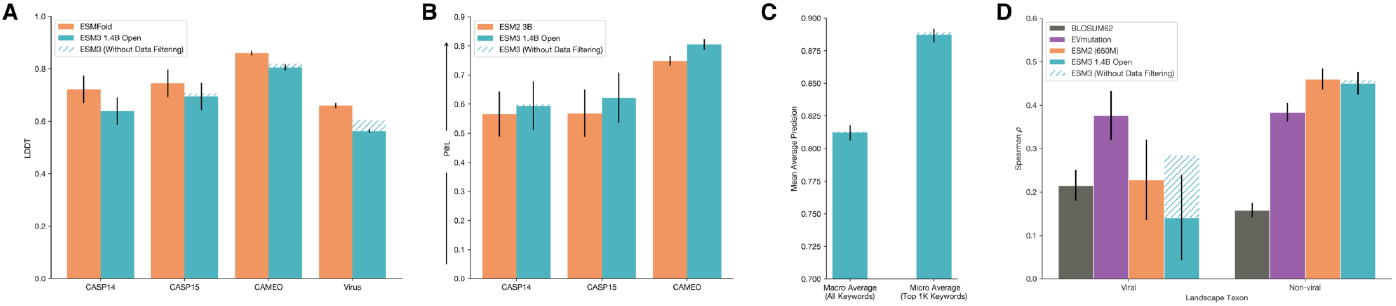
ESM3-open is a powerful predictor of structure and function trained for open release. A: Structure Prediction Single-pass structure prediction for ESM3-open (blue; quadratic time complexity) compared with ESMFold (orange; cubic time complexity). See Appendix A.3.4 for details on this evaluation. B: Representation Learning ESM3-open (blue) is competitive with ESM2-3B (orange) on representation learning as measured by contact prediction P@L for finetuned representations. See Appendix A.3.3 for details on this evaluation. C: Function Keyword Prediction. ESM3-open function prediction performance, as measured by Mean Average Precision across function keywords. ESM3-open achieves 0.81 precision across all keywords, and 0.89 for the top 1K most prevalent keywords in the test set (Appendix A.3.2). We use the same evaluation framework as in Appendix A.1.8.2.2. We report both the macro and micro averages as in Fig. S8. In each of the preceding evaluations, the data mitigation minimally impacted performance, as compared to a compute-matched model without data mitigations (hatched blue). D: Zero-shot Fitness Prediction. Fitness prediction performance as measured by correlation (Spearman *ρ*) across 217 Deep Mutational Scanning datasets collated in ProteinGym. Left and right subplots indicate viral (left) and non-viral (right) DMS datasets. The four columns per group indicate different models. ESM3-open performs substantially worse than EVMutation (purple) on viral fitness prediction, while being competitive with ESM2 (orange) on non-viral fitness prediction. Viral fitness prediction was substantially impacted by the data mitigation, while non-viral fitness prediction was not (hatched blue).

##### A.6.2 ESM3-open Evaluations

In the section below, we outline our evaluations of ESM3-open performance. When appropriate, we compare ESM3-open to either existing open models, (e.g. ESM2 or ESM-Fold), or to a compute-matched version of ESM3-open, trained without any data mitigations.

###### Structure Prediction

In Fig. S24A, we show that ESM3-open achieves competitive performance on structure prediction as measured by LDDT on CASP14, 15 and CAMEO, showing very slight degradation from our compute-matched 1.4B model without data filtering. The evaluation framework is described in Appendix A.3.4.

We also measure the ability of ESM3 to predict the structure of a subset of viral proteins. In Fig. S24A we evaluate structure prediction on a set of structures derived from viruses that were purged from the PDB training set. For the chains in PDB that were *>* 70% sequence identity hits to the Viral Denylist, we cluster at 40% sequence identity and then select the longest chain (with length ≤ 1024) from each cluster. ESM3-open has an average LDDT of 0.56 on the viral structures. Without the data mitigation, a compute-matched ESM3-open has an average LDDT of 0.60.

###### Representation Learning

ESM3-open achieves strong performance on representation learning, slightly outperforming ESM2 (3B) on contact prediction as measured by precision at L (P@L) on structures derived from CASP14/15, and CAMEO, see Fig. S24B. The evaluation framework is described in Appendix A.3.3.

###### Function Keyword Prediction

ESM3-open is able to predict function keywords for proteins in a validation set derived from UniRef and annotated with InterProScan, see Fig. S24C. ESM3-open achieves a Mean Average Precision for all keywords of 0.81 (macro average), and a precision of 0.89 (micro average) for the top 1000 keywords, discarding common terms such as “the”. The evaluation framework is the same as that described in Appendix A.1.8.2.2.

###### Zero-shot Viral Fitness Prediction

We measure the ability of ESM3 to identify viable sequences and understand the effects of mutations on viral proteins. The evaluation consists of the single mutant variants from 217 Deep Mutational Scanning (DMS) datasets collected in ProteinGym (117). This includes 28 DMS landscapes from viral proteins and 189 from other proteins. We evaluate the correlation (Spearman *ρ*) between the predicted variant effect and measured variant effect. The predicted variant effect is measured as the difference between the logit value for the variant allele and the logit value of the wildtype allele at a given masked position (18).

First, we compare the performance of ESM3-open to a compute-matched version of ESM3-open which did not undergo any data filtering. We also compare the performance of ESM3-open to existing open model baselines. Fig. S24D assesses performance relative to the EVMutation (118) baseline. EVMutation is a Markov Random Field model (not deep learning-based) trained on a multiple sequence alignment of the target protein. BLOSUM62 is a baseline based on amino acid substitution frequencies. Before mitigations, ESM3-small outperforms ESM2 on viral landscapes, but falls short of EVMutation. EVMutation uses lower sequence diversity thresholds when generating MSAs for viral proteins, while ESM2 and ESM3 use fixed 50% and 70% clustering for their training sets respectively, which may explain the difference in capability. On non-viral landscapes, ESM3 performs on par with ESM2. Applying data filtering as a mitigation reduces average Spearman *ρ* performance on viral fitness prediction from 0.28 (ESM3-small) to 0.17 (ESM3-open), to the level of a BLOSUM prior, which incorporates no knowledge of the statistics of the protein family. This indicates that the model contains essentially no information about the evolutionary statistics of the evaluated viral protein families. Performance on non-viral proteins is not adversely affected, changing from 0.46 (ESM3-small) to 0.45 (ESM3-open).

